# Chemical genetics reveals *Leishmania* KKT2 and CRK9 kinase activity is required for cell cycle progression

**DOI:** 10.1101/2025.09.08.674917

**Authors:** Juliana B. T. Carnielli, James A. Brannigan, Nathaniel G. Jones, Priscila Z. Ramos, Rafael M. Couñago, Peter Sjö, Ana Paula C. A. Lima, Anthony J. Wilkinson, Jeremy C. Mottram

## Abstract

Protein kinases are key regulators of the eukaryotic cell cycle and have consequently emerged as attractive targets for drug development. Their well-defined active sites make them particularly amenable to inhibition by small molecules, underscoring their druggability. The *Leishmania* kinome, shaped by diverse evolutionary processes, harbours a unique repertoire of potential drug targets. Here, we used the cysteine-directed protein kinase probe SM1-71 to identify four essential protein kinases MPK4, MPK5, MPK7 and AEK1 as candidates for covalent kinase inhibitor development, as well as CLK1/CLK2 for which covalent inhibitors have already been identified. We leveraged the absence of natural analog-sensitive (AS) kinases in *L. mexicana* to establish an *in vivo* chemical-genetic AS kinase platform for investigating essential functions of protein kinases. Using CRISPR-Cas9-mediated precision genome editing, we endogenously engineered two kinetochore-associated protein kinases, KKT2 and KKT3, and cyclin-dependent kinase CRK9, to generate AS kinases. We show that KKT2 and CRK9 kinase activities are essential for both promastigote and intracellular amastigote survival; KKT2 kinase activity being required for progression through mitosis at a stage preceding mitotic spindle assembly, while CRK9 kinase activity is required for S phase, consistent with its role in trans-splicing. This study demonstrates the utility of AS chemical genetics in *Leishmania* and identifies KKT2 and CRK9 as having critical roles in *Leishmania* cell cycle regulation and therefore being promising drug targets.

## Introduction

Leishmaniasis are a group of neglected tropical diseases caused by protozoan parasites of the genus *Leishmania*, transmitted to mammalian hosts by the bite of infected female sandflies. The clinical manifestations of leishmaniasis are species-specific and primarily include cutaneous leishmaniasis (CL), mucocutaneous leishmaniasis (MCL), and visceral leishmaniasis (VL). Strongly associated with poverty, leishmaniasis is endemic in 99 countries, placing over one billion people at risk and resulting in an estimated one million new cases annually worldwide [1, 2].

In the absence of an effective vaccine, the limited available chemotherapies play a critical role in the control of the leishmaniasis. Currently approved treatments include pentavalent antimonials, amphotericin B, miltefosine, paromomycin, and pentamidine. However, these drugs are associated with severe shortcomings, including toxicity, the emergence of drug-resistant parasite, and restricted accessibility in low-resource settings. Moreover, the mechanisms of action of most anti-leishmanial agents remain incompletely understood, impeding the rational design of combination therapies and the development of robust drug resistance surveillance strategies. To address these gaps, drug discovery efforts have increasingly leveraged phenotypic screening coupled with target deconvolution, leading to the identification of promising candidates with known mechanisms of action. Notable examples include selective inhibitors of the kinetoplastid proteasome (GNF6702 and GSK3494245) [3, 4]; the pyrazolopyrimidine compound GSK3186899, which exerts its activity through inhibition of the parasite’s cyclin-dependent kinase CRK12 [5]; the amidobenzimidazole AB1 from Novartis, which selectively targets the parasite kinetochore kinases CLK1/CLK2 [6, 7]; the benzoxaborole derivative DNDI-6148, which acts by inhibition of *Leishmania* cleavage and polyadenylation specificity factor (CPSF3) endonuclease [8]; and, more recently, the GSK pyrrolopyrimidine derivative, which inhibits parasite growth via targeting of mitochondrial cytochrome b [9]. Despite these advances, target deconvolution can be a major challenge, particularly when compounds act on non-protein targets or exhibit polypharmacology [10, 11]. In this context, target-based drug discovery offers a complementary strategy, as it provides direct mechanistic insight and facilitates detailed structure-activity relationship studies. Nevertheless, the success of such approaches relies heavily on the availability of well-validated drug targets. For trypanosomatids, progress in this area has been limited, with only a few targets meeting established chemical and genetic criteria for target validation. Consequently, the identification of novel therapeutic targets and elucidation of their biological functions within the parasite are essential for advancing drug discovery and improving disease management.

The human kinome has been extensively explored as a therapeutic landscape, leading to the successful development of kinase inhibitors for the treatment of different types of cancers, as well as inflammatory, autoimmune, and neurodegenerative diseases [12]. This remarkable clinical impact has catalysed interest in exploring the kinome of other organisms, including protozoan parasites such as *Leishmania*, as sources of novel drug targets. The kinome of *L. mexicana*, with 193 eukaryotic protein kinases (ePKs) predicted in its genome, is roughly one-third the size of the human kinome [13, 14]. Based on conserved catalytic domain sequences, ePKs are broadly classified into two major superfamilies: serine/threonine (Ser/Thr) kinases, which are ubiquitous across eukaryotes; and tyrosine (Tyr) kinases whose receptor and receptor-like members are absent in *Leishmania*. However, these organisms possess dual-specificity kinases and atypical tyrosine kinases, which likely account for protein tyrosine phosphorylation observed in their proteomes [15–18]. The *Leishmania* kinome includes homologs of the five major ePK groups found in higher eukaryotes – AGC, CAMK, CK1, CMGC, and STE – as well as members of the NEK family, which represents a distinct evolutionary branch of the human kinome. Additional kinases that do not fit into these canonical groups are categorized as “others” and include key regulators of eukaryotic cell signalling such as Aurora and Polo-like kinases, as well as the highly divergent kinetoplastid kinetochore proteins KKT2 and KKT3. Furthermore, atypical protein kinases (aPKs), which lack a conserved ePK catalytic domain but retain kinase activity, have also been identified in the *Leishmania* genome. Among these are members of the phosphatidylinositol 3′-kinase-related kinase (PIKK) group, which resemble lipid kinases [13].

Our recent functional study of the *Leishmania* kinome showed that approximately 21% of *Leishmania* protein kinases are required for promastigote survival, highlighting their potential as drug targets [14]. Notably, two preclinical candidates have provided proof-of-concept for kinase-targeted therapy in *Leishmania*: GSK3186899, a pyrazolopyrimidine that targets CRK12 [5]; and the amidobenzimidazole AB1, which selectively targets the kinetochore kinases CLK1/CLK2 [7]. These advances underscore the therapeutic promise of the *Leishmania* kinome and provide a level of validation for protein kinases as a viable class of targets for anti-leishmanial drug development.

The successful development of kinase-targeting therapeutics relies heavily on robust preclinical target validation, which involves elucidating the biological functions and signalling roles of the target protein kinase to assess its suitability as a drug target. In the absence of selective inhibitors for individual kinases, the study of kinase function has been significantly advanced by the use of the chemical-genetic analog-sensitive (AS) kinase approach. This enables precise and reversible inhibition of kinase activity. This strategy involves mutating the conserved, bulky gatekeeper residue within the ATP-binding pocket of the kinase to an amino acid with a smaller side chain, such as glycine or alanine. This substitution enlarges the ATP-binding pocket, rendering the engineered kinase uniquely sensitive to cell-permeable “bumped” kinase inhibitors (BKIs), structurally modified ATP analogs that do not bind to the wild-type kinase ATP-binding pocket, but can selectively inhibit the analog-sensitive variant [19–21].

AS kinase technology has been successfully applied to investigate protein kinase signalling in diverse cellular processes such as cell cycle [22, 23], immunological T and B cell activation [24, 25], and apicomplexan parasite egress [26]. In kinetoplastid parasites, this approach brought to the light the biological role of the cyclin-dependent kinases CRK1 and CRK9 in *Trypanosoma brucei*. CRK1 has been shown to be involved in anterograde protein trafficking [27], and to promote the G1/S phase transition of the cell cycle, in coordination with global protein translation [28], while CRK9 is essential for pre-mRNA processing [29]. Additionally, AS-mediated inhibition of polo-like kinase has revealed its critical role in flagellar attachment and cytokinesis in *T. brucei* [30]. Importantly, this chemical-genetic strategy has also contributed to drug discovery by validating AGC essential kinase 1 (AEK1) as a potential drug target in *T. brucei* [31] and *T. cruzi* [32].

Equally important to the identification of potential drug targets is the ability to rationally design selective inhibitors against them. In this context, substantial progress has been achieved through the development of targeted covalent kinase inhibitors. These inhibitors are typically engineered by combining a kinase-binding scaffold with an electrophilic warhead capable of forming an irreversible covalent bond with nucleophilic amino acid residues [33]. The most commonly targeted residue is cysteine, due to its high intrinsic nucleophilicity [34]. Notably, the eleven FDA-approved covalent kinase inhibitors operate through a shared mechanism involving the Michael addition of a protein cysteine thiolate to an acrylamide or acrylamide-like electrophile [35]. This strategy has also shown promise against trypanosomatid parasites, where CLK1/CLK2 inhibition by AB1 illustrates the potential of covalent kinase inhibitors in the context of trypanosomatid drug discovery [6]. Therefore, the identification of reactive cysteine residues within protein kinases has emerged as a valuable approach to guide rational covalent inhibitor design. One powerful tool in this effort has been the promiscuous covalent kinase inhibitor SM1-71, originally developed to target cysteines located immediately N-terminal to the conserved DFG motif (DFG–1) in human protein kinases. The use of SM1-71 and its biotinylated analog has enabled systematic mapping of reactive cysteine residues across the human kinome, providing key insights into targetable nucleophilic sites for covalent ligand development [36, 37].

In this study, we explored the active sites of the *Leishmania* kinome using the SM1-71 probe. We identified 10 kinases with targetable cysteine residues, five of which are known to be essential for parasite survival. Our bioinformatic and structural analysis indicate an absence of naturally occurring AS kinases that have glycine or alanine gatekeeper residues, making this parasite a suitable model for the AS chemical genetics strategy in the promastigote and, importantly, the intracellular amastigote stages. We showed that BKI-mediated inhibition of KKT2 AS resulted in mitotic arrest at a stage prior to spindle assembly, whilst inhibition of CRK9 AS led to impaired DNA synthesis, resulting in cell death.

## Results

### *Leishmania* kinome, a source of attractive drug targets

From our previous systematic analysis of the *Leishmania* kinome, 43 protein kinases were highlighted as potentially being essential in the promastigote stage, since a null mutant could not be recovered [14]. Here, we reanalysed the Bar-Seq dataset generated from the pooled barcoded library of kinase deletion mutants, to identify protein kinases required for amastigote-stage survival. The proportion of barcodes in the metacyclic promastigote stage, as well as at two post-infection time points – in *in vitro* macrophage (12 and 72 hours) and in *in vivo* mouse footpad (3 and 6 weeks) experiments – was used to calculate fold changes in barcode representation relative to the preceding time point. A protein kinase was classified as required for amastigote survival if the corresponding mutant cell line exhibited the following trajectory in barcode representation: (i) a significant ≥2-fold reduction in barcode abundance at the first post-infection time point, followed by an additional decrease of ≥30% at the second time point; or (ii) no significant change in the first time point, but a significant ≥50% reduction at the final time point (Supplementary Fig 1). This scoring approach was expected to exclude kinases primarily involved in parasite infection or differentiation. This analysis identified 21 protein kinases required for amastigote survival. These proteins were distributed across kinase families as follows: AGC (1), CAMK (2), CK1 (2), CMGC (9), NEK (2), STE (2), and Other (3). Furthermore, our previously determined subcellular localization data for *L. mexicana* protein kinases [14] revealed that the protein kinases assigned as required for amastigote survival in this study are predominantly localized in the cytoplasm (57.1%) and, to a lesser extent, the nucleus (14.3%) of the parasite (Table 1, and Supplementary Data 1).

**Table 1.** *L. mexicana* protein kinases required for amastigote survival.

| Gene ID | Name | Group/Family | Amastigote survival requirement |  | Primary localization <sup>c</sup> |
| --- | --- | --- | --- | --- | --- |
|  |  |  | inMAC <sup>a</sup> | FP <sup>b</sup> |  |
| LmxM.28.1670 | ZFK | AGC | **Required** | Dispensable | Lysosome |
| LmxM.19.1470 |  | CAMK/CAMKL | Dispensable | **Required** | Cytoplasm |
| LmxM.22.0810 | SOS2 | CAMK | Dispensable | **Required** | Basal body |
| LmxM.04.1210 | CK1.3 | CK1/CK1 | Dispensable | **Required** | Cytoplasm |
| LmxM.33.3020 |  | CK1/TTBK | **Required** | Dispensable | Flagellar pocket |
| LmxM.09.0400 | CLK1 (KKT10) | CMGC/CLK | **Required** | Not known | Nucleus |
| LmxM.09.0410 | CLK2 (KKT19) | CMGC/CLK | **Required** | Not known | Nucleus |
| LmxM.13.1640 | MPK7 | CMGC/MAPK | Dispensable | **Required** | Cytoplasm |
| LmxM.20_36.6470 | MPK1 | CMGC/CDKL | **Required** | Dispensable | Flagellum |
| LmxM.21.1650 | PK4 | CMGC | Dispensable | **Required** | Cytoplasm |
| LmxM.25.1560 |  | CMGC/CLK | Dispensable | **Required** | Cytoplasm |
| LmxM.29.0370 | MPK12 | CMGC/MAPK | Dispensable | **Required** | Cytoplasm |
| LmxM.29.2910 | MPK5 | CMGC/MAPK | Dispensable | **Required** | Pellicular membrane |
| LmxM.36.4250 |  | CMGC/GSK | Dispensable | **Required** | No signal |
| LmxM.10.0830 |  | Other/Orphan | Dispensable | **Required** | Nucleus |
| LmxM.33.0030 |  | Other/NAK | Dispensable | **Required** | Cytoplasm |
| LmxM.33.0940 | WEE1 | Other/WEE | Dispensable | **Required** | Cytoplasm |
| LmxM.08_29.2570 |  | NEK | Dispensable | **Required** | Cytoplasm |
| LmxM.30.3160 |  | NEK | Dispensable | **Required** | Cytoplasm |
| LmxM.07.0250 |  | STE | Dispensable | **Required** | Cytoplasm |
| LmxM.08_29.2320 | MKK1 | STE | Dispensable | **Required** | Cytoplasm |

<sup>a</sup> inMAC, *in vitro* infected macrophage. <sup>b</sup> FP, *in vivo* mouse footpads. <sup>c</sup> protein localisation as previous described by Baker et al., 2021 [14].

From our comprehensive analysis of the kinase requirements in *Leishmania*, 21 protein kinases were identified as essential for the survival of the amastigote stage, in addition to the 43 previously shown to be required in promastigotes [14]. This brings the total to 64 protein kinases that represent potential drug targets. Among these, particular attention was drawn to the four Kinetoplastid Kinetochore Proteins (KKTs) – CLK1 (LmxM.09.0400, also annotated as KKT10), CLK2 (LmxM.09.0410, also annotated as KKT19), KKT2 (LmxM.36.5350), and KKT3 (LmxM.34.4050) – due to their divergent evolutionary roots in the conserved eukaryotic chromosome segregation process, resulting in a unique and druggable repertoire specific to trypanosomatids. We have previously validated CLK1 and/or CLK2 as druggable targets in trypanosomatids through the development of AB1 [6, 7, 38] and the third-generation EGFR inhibitor WZ8040, using a bioluminescence resonance energy transfer (BRET)-based target engagement assay in live cells [39].

### *L. mexicana* protein kinase function investigated using chemical-genetics

Given that the most robust target validation strategies enable assessment of drug effects *in vivo*, we sought to engineer analog-sensitive kinases directly in *Leishmania*. Genetic modification of the gatekeeper residue was introduced in the amenable promastigote stage, followed by differentiation into intracellular amastigotes – the clinically relevant, drug-targetable form in the mammalian host. This strategy allows evaluation of the functional requirement of the engineered kinases across both life stages using viability assays. If the target kinase activity is essential, AS kinase-expressing parasites would be expected to exhibit increased susceptibility to BKIs relative to the wild-type T7/Cas9 line.

To assess the feasibility of using AS kinase technology to investigate the requirement of protein kinases for *Leishmania* survival, we first screened the *L. mexicana* kinome for the presence of naturally occurring AS kinases that might interfere with our approach. A natural AS kinase is defined as a kinase that has a glycine or alanine residue in the gatekeeper position. To address this, we performed a comprehensive protein sequence and structural analysis of all 183 *bona fide* ePKs described in the *L. mexicana* kinome to determine the identity of their gatekeeper residues. Of the 193 predicted ePKs previously annotated in the *L. mexicana* kinome [14], 10 are classified as pseudokinases as they lack all three of the conserved catalytic residues (K D D) required for kinase activity (Fig 1a). Given their lack of enzymatic function and demonstrated non-essentiality, pseudokinases were not analysed for gatekeeper residue identity.

**Fig 1.**
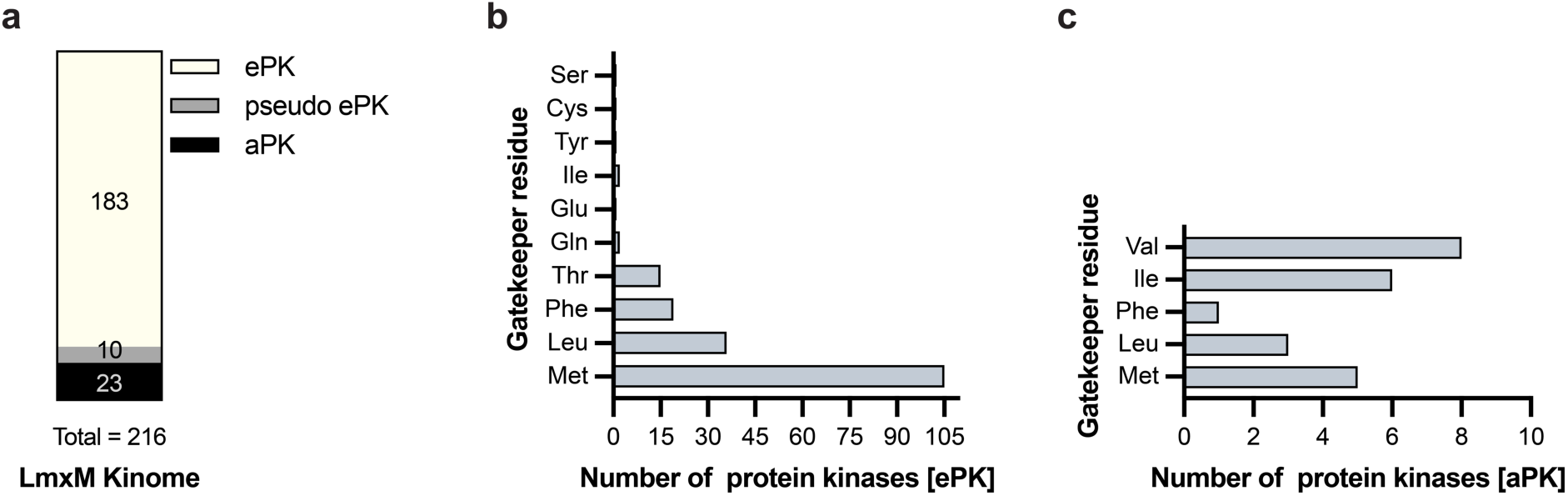
The *L. mexicana* kinome lacks natural analog-sensitive kinases. (a) Schematic overview of the *L. mexicana* kinome, comprising eukaryotic protein kinases (ePKs) and atypical protein kinases (aPKs). ePKs lacking all three key residues (K D D) required for catalytic activity were classified as pseudo ePKs. (b) Distribution of gatekeeper residues among the 183 bona fide *L. mexicana* ePKs. (c) Gatekeeper residue distribution among the 23 *L. mexicana* aPKs.

Our analysis revealed that methionine is the predominant gatekeeper residue, present in 105 of the 183 genuine ePKs (57.4%). Leucine (19.7%), phenylalanine (10.4%), and threonine (8.2%) were the next most abundant gatekeeper residues. Notably, the AGC kinase subgroup was the only family in which methionine was not the most frequent gatekeeper; instead, leucine was most prevalent (63.6%). The two smallest gatekeeper residues identified were: a cysteine in the member of the family STE LmxM.36.0910; and a serine in LmxM.36.4250, a member of the CMGC family (Fig 1b, Supplementary Fig 2, and Supplementary Data 1).

We also analysed the gatekeeper residues of 23 atypical protein kinases. Interestingly, valine was the most frequent gatekeeper residue among aPKs, a residue not observed at this position in any ePK (Fig 1c, Supplementary Fig 2, and Supplementary Data 1). However, due to their distinct structural features, aPKs are inhibited by a different set of BKIs than those used for analog-sensitive ePKs (e.g., LY294002 analogues targeting PI3K-like kinases) [40].

Together, these findings demonstrate that *L. mexicana* lacks natural analog-sensitive kinases, supporting the suitability of this parasite as a model for functional dissection of protein kinase activity using AS technology.

We selected the kinetochore kinases CLK1/CLK2, KKT2 and KKT3 for functional validation using the chemical-genetic AS kinase strategy. Additionally, we have also selected the cyclin-dependent kinase CRK9 (LmxM.27.1940) to target, since this essential protein kinase was shown to be amenable to the AS kinase approach in the closely relate trypanosomatid *T. brucei* [29]. Orthologs of *L. mexicana* CLK1/CLK2, KKT2, KKT3 and CRK9 are present and highly conserved in all high-quality sequenced genomes of the order *Trypanosomatida* that are known human pathogens (Supplementary Fig 3), making studies on *Leishmania* relevant to multiple clinically relevant trypanosomatid parasites. Our gatekeeper residue analysis revealed that a bulky methionine occupies the gatekeeper position of the ATP-binding site of these five selected kinase targets (Fig 2a).

**Fig 2.**
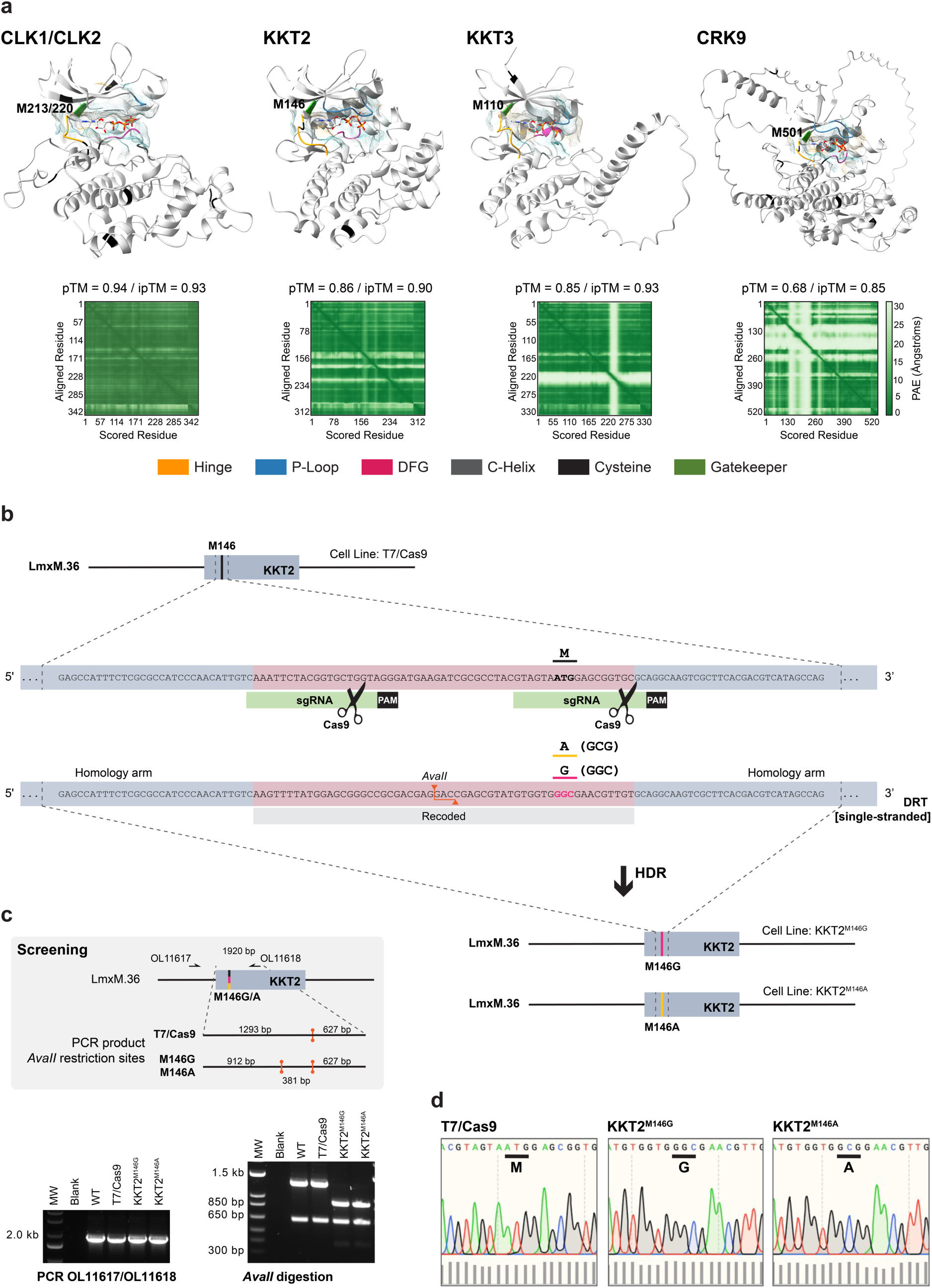
Marker-free CRISPR-Cas9–mediated precision genome editing to engineer analog-sensitive kinases in *Leishmania*. (a) Structural model of the kinase domains of CLK1/CLK2, KKT2, KKT3, and CRK9 generated using AlphaFold 3 [41] and visualized with ChimeraX v1.9. The ATP ligand is displayed as a stick model, with heteroatom-based colouring. A semi-transparent molecular lipophilicity potential (MLP) surface is overlaid, with colour ranging from dark cyan (most hydrophilic) to white to dark goldenrod (most lipophilic). AlphaFold confidence metrics are displayed below each structure, including the predicted template modelling score (pTM), the interface predicted template modelling score (ipTM) between the kinase and ATP, and the predicted aligned error (PAE) plot. (b) Schematic representation of the CRISPR-Cas9 strategy used to engineer analog-sensitive kinases by substituting the gatekeeper methionine (M) with glycine (G) or alanine (A). The example illustrates the replacement of the gatekeeper residue in KKT2. Linear DNA fragments for *in vivo* transcription of two single guide RNAs (sgRNAs) and a 120 bp DNA repair template containing silent recoding mutations and the gatekeeper substitution were used. The mutations introduced an *Ava*II restriction site, enabling genotypic screening of edited clones. PAM, protospacer adjacent motif; HDR, homology-directed repair; LmxM.36, *L. mexicana* chromosome 36. (c) Genotyping workflow (top grey box) and PCR-restriction digest results (bottom) for selected analog-sensitive clones. (d) Sanger DNA sequencing of the engineered KKT2 locus confirms the substitution of the gatekeeper methionine with glycine or alanine in the KKT2^M146G^ and KKT2^M146A^ lines, respectively. Sequencing chromatograms were visualized in SnapGene v7.2; bar graphs below indicate per-base quality scores.

### CRISPR-Cas9–mediated precision genome editing to engineer *Leishmania* protein kinases

The *L. mexicana* cell line stably expressing T7 RNA polymerase and Cas9 endonuclease [42] was used as the parental strain for engineering AS protein kinases via CRISPR-Cas9–mediated precision genome editing. Analog-sensitive kinase mutants were generated by introducing specific point mutations at the endogenous kinase *loci*, replacing the native bulky gatekeeper residue with either glycine or alanine. This was achieved using a 120 bp single-stranded DNA repair template lacking selectable drug resistance markers. To direct the desired edits, two single guide RNAs (sgRNAs) were designed to flank the gatekeeper codon. In addition to the codon substitution, the DNA repair template was engineered to contain silent mutations within the protospacer adjacent motif (PAM) and sgRNA target sites to prevent its cleavage by Cas9 (Fig 2b). These silent mutations also introduced/removed restriction endonuclease cleavage sites, enabling discrimination between wild-type and mutant alleles for downstream screening. Ten clones per transfection were screened by PCR followed by restriction enzyme digestion and then confirming by Sanger DNA sequencing (Fig 2c – d). CLK1 and CLK2 share 86% sequence identity, and their kinase domains, which comprise the gatekeeper residue (M213 in CLK1 and M220 in CLK2), are identical (Supplementary Fig 4a). As a result, it is not feasible to selectively target CLK1 without also affecting CLK2, and vice versa. Therefore, both kinases were simultaneously targeted in this study. Precision editing of CLK1 or CLK2 was not achieved, possibly because the AS mutation is not tolerated by the kinase (Supplementary Fig 4b – e). This strategy, however, was successful in the generation of AS kinase mutants for KKT2, KKT3 and CRK9 with 3/10 success rate for the KKT2^M146G^ substitution, 1/10 for KKT2^M146A^, 1/10 for KKT3^M110G^, and 1/10 for both CRK9^M501G^ and CRK9^M501A^ variants (Supplementary Fig 5 – 7). These findings demonstrate that *L. mexicana* is amenable to marker-free precision genome editing using the CRISPR-Cas9 system, with high editing efficiency achieved across multiple protein kinase targets.

### KKT2 and CRK9 protein kinase activity is essential for *Leishmania*

The requirement for kinase activity was initially evaluated in the promastigote stage of the *Leishmania* parasite by comparing the sensitivity of parasites expressing either wild-type or analog-sensitive kinases to the bumped kinase inhibitors 1NM-PP1 and 1NA-PP1 (Supplementary Fig 8a). Using a resazurin-based viability assay, the effects of the tested BKIs on *Leishmania* promastigote viability revealed that 1NM-PP1 exhibited stronger inhibitory activity, particularly against AS kinases with a glycine residue at the gatekeeper position, suggesting more efficient binding in these mutants. On the other hand, 1NA-PP1 displayed reduced specificity, showing comparable inhibition of AS kinases with either glycine or alanine at the gatekeeper site. Parasites expressing AS variants of KKT2 and CRK9 exhibited marked growth inhibition at BKI concentrations that had minimal effect on the wild-type line (T7/Cas9 IC_10_). Furthermore, the AS mutants exhibited significantly increased sensitivity to BKIs, as indicated by lower half-maximal inhibitory concentration (IC_50_) values compared to the wild-type line. Our result on the sensitivity of CRK9 AS to BKIs in promastigote stage was previously published by Jones N.G. et al., 2023 [43]. Using that dataset, we calculated IC_50_ values and compared them to those of the parental line T7/Cas9, which displayed IC_50_ values of 11.26 µM for 1NM-PP1 and 12.80 µM for 1NA-PP1. The CRK9^M501A^ showed reduced IC_50_ values of 6.22 µM for 1NM-PP1 (p-value = 0.0469) and 3.36 µM for 1NA-PP1 (p-value = 0.0469), while the CRK9^M501G^ exhibited even greater sensitivity, with IC_50_s of 0.63 µM for 1NM-PP1 (p-value = 0.0071) and 2.32 µM for 1NA-PP1 (p-value = 0.0231). These findings indicate that the kinase activity of KKT2 and CRK9 is essential for promastigote survival (Fig 3a and Supplementary Fig 8b).

**Fig 3.**
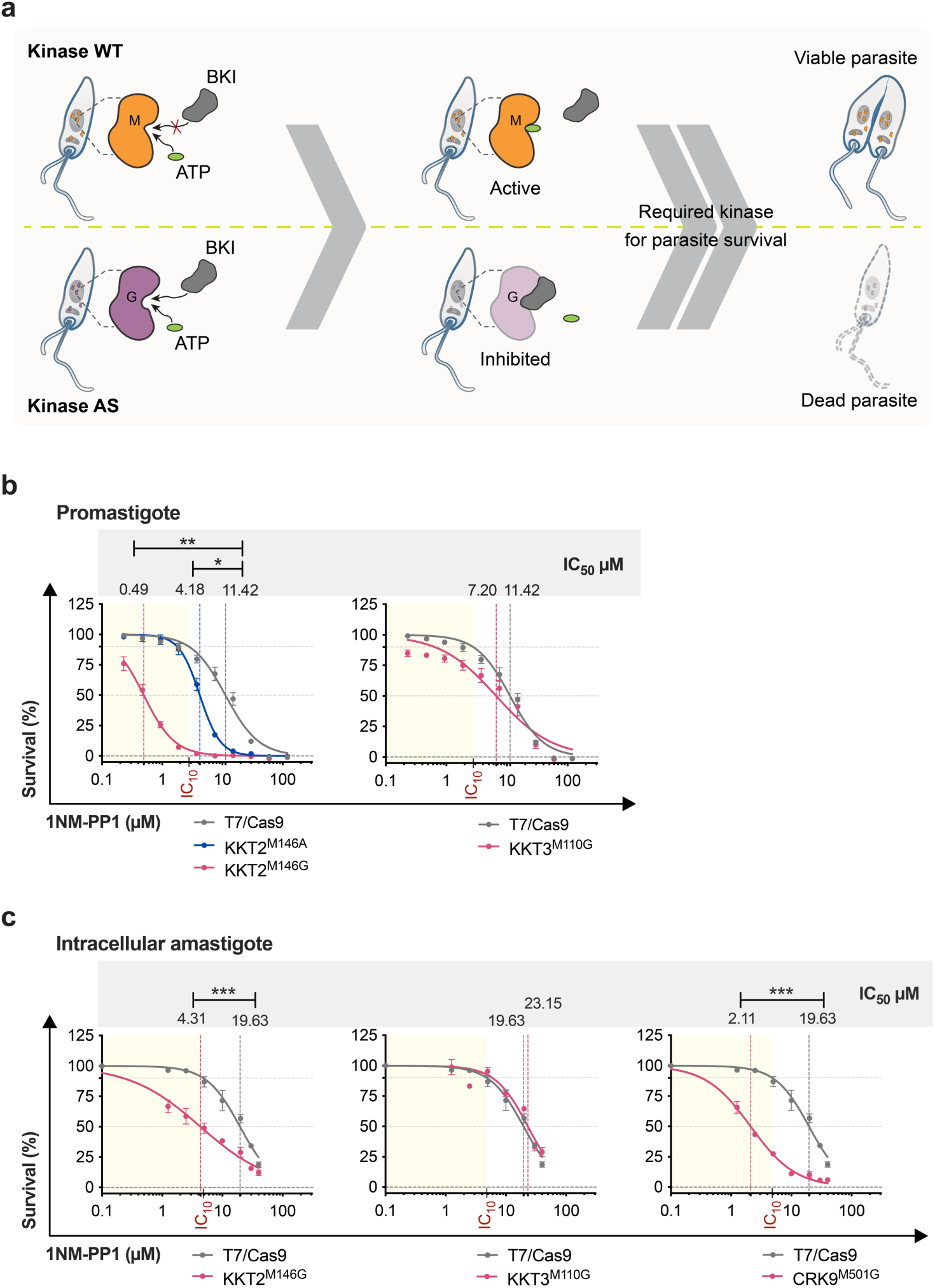
The kinase activities of KKT2 and CRK9, but not KKT3, are essential for *Leishmania* survival. (a) Schematic representation of the viability assay performed on parasites expressing either the wild-type or AS variant, illustrating the expected outcome if the targeted protein kinase is essential for parasite survival following treatment with a bumped kinase inhibitor. The same experimental principle was applied to the amastigote viability assay. WT, wild type; AS, analog-sensitive; Cas9, endonuclease; sgRNA, single guide RNA; PAM, protospacer adjacent motif; M, methionine; G, glycine; BKI, bumped kinase inhibitor. Parasite viability under 1NM-PP1 treatment was assessed in *L. mexicana* lines expressing AS variants of KKT2, KKT3, and CRK9, as well as the wild-type T7/Cas9 line. Dose-response curves were fitted using GraphPad Prism v10.4.1, with viability normalized to untreated controls (set at 100% for each cell line). Statistical significance was evaluated using unpaired two-tailed Student’s t-tests (*p-value <0.05; **p-value <0.01; ***p-value <0.001). The IC10 value denotes the concentration of 1NM-PP1 that reduces the viability of the wild-type T7/Cas9 line by 10%. (b) Susceptibility of promastigotes to 1NM-PP1 was measured using a resazurin-based viability assay. Data represent mean ± SEM from three biological replicates. (c) Susceptibility of intracellular amastigotes to 1NM-PP1 was assessed by quantifying the percentage of infected macrophages. Data represent mean ± SEM from four biological replicates.

Although the AS kinase mutants retained sufficient activity to support promastigote growth in culture, we assessed whether the introduced mutations affected parasite fitness. Growth rate comparisons between AS lines carrying glycine gatekeeper mutations and the parental line revealed no significant differences (Supplementary Fig 9), confirming that the increased drug sensitivity observed in viability assays was due to specific inhibition of kinase activity, rather than a fitness defect.

Given the increased susceptibility of parasites expressing analog-sensitive kinases with a glycine gatekeeper to BKIs, these lines were selected for intracellular amastigote susceptibility assays. Notably, 1NA-PP1 exhibited substantially lower activity against the intracellular amastigote stage compared to 1NM-PP1 (Fig 3b and Supplementary Fig 8c), possible due to reduced stability of 1NA-PP1 within the acidic environment of the macrophage parasitophorous vacuole. Consistent with observations in the promastigote stage, parasites expressing AS variants of KKT2 and CRK9 showed significantly increased susceptibility to 1NM-PP1, indicating that the kinase activity of these proteins is also critical for survival of the intracellular amastigote stage (Fig 3b).

The susceptibility of parasites expressing AS variant of KKT3 to the BKIs did not differ from that of the wild type line in either the promastigote or intracellular amastigote stages (Fig 3 and Supplementary Fig 8), suggesting that KKT3 kinase activity is not essential for parasite survival. To reassess the essentiality of *KKT3*, we attempted CRISPR-Cas9-mediated gene deletion using single guide RNAs (sgRNAs) designed with the updated LeishGEdit algorithm for Bar-Seq applications [44]. Puromycin and blasticidin double-resistant parasites were recovered in two of three independent transfections. However, PCR analysis revealed that the KKT3 coding sequence was still present in recovered clones, suggesting that the gene or chromosome may have been duplicated to retain an essential function. Whole-genome sequencing of three recovered clones confirmed gene duplication of the *KKT3* locus, supporting the hypothesis that KKT3, but not its kinase activity, is essential for promastigote viability (Supplementary Fig 10).

To gain deeper insights into the essential roles of KKT2 and CRK9 we exploited the ability to selectively inhibit their kinase activity to investigate their biological functions in *Leishmania*.

### KKT2 kinase activity is required for coordinated progression through S and M phases in *Leishmania*

The effect of KKT2 inhibition on cell cycle progression in *Leishmania* promastigotes was assessed by measuring the DNA content via flow cytometry. Cells were stained with propidium iodide (PI), and its distribution across cell cycle phases was determined using the Watson model algorithm in FlowJo v10.10.0. Following treatment with 5 µM 1NM-PP1 for 6 hours, parasites expressing the KKT2^M146G^ analog-sensitive variant exhibited a significant accumulation of cells in the G2/Mitosis phase (30.6%), compared to the untreated control (21.5%) and those expressing wild-type KKT2 (21.3%), indicating cell cycle arrest at this stage (Fig 4a and Supplementary Fig 11a). After 24 hours of treatment, a further increase in cells accumulating in the S phase was detected in the parasites expressing the KKT2 AS variant. While the Watson model predominantly classified these cells as being in S phase, visual inspection of the histograms indicated that the G2/Mitosis population was also enlarged. Furthermore, it was observed that the size of arrested cells did not differ from control cells at the corresponding cell cycle stage (Fig 4a and Supplementary Fig 11b).

**Fig 4.**
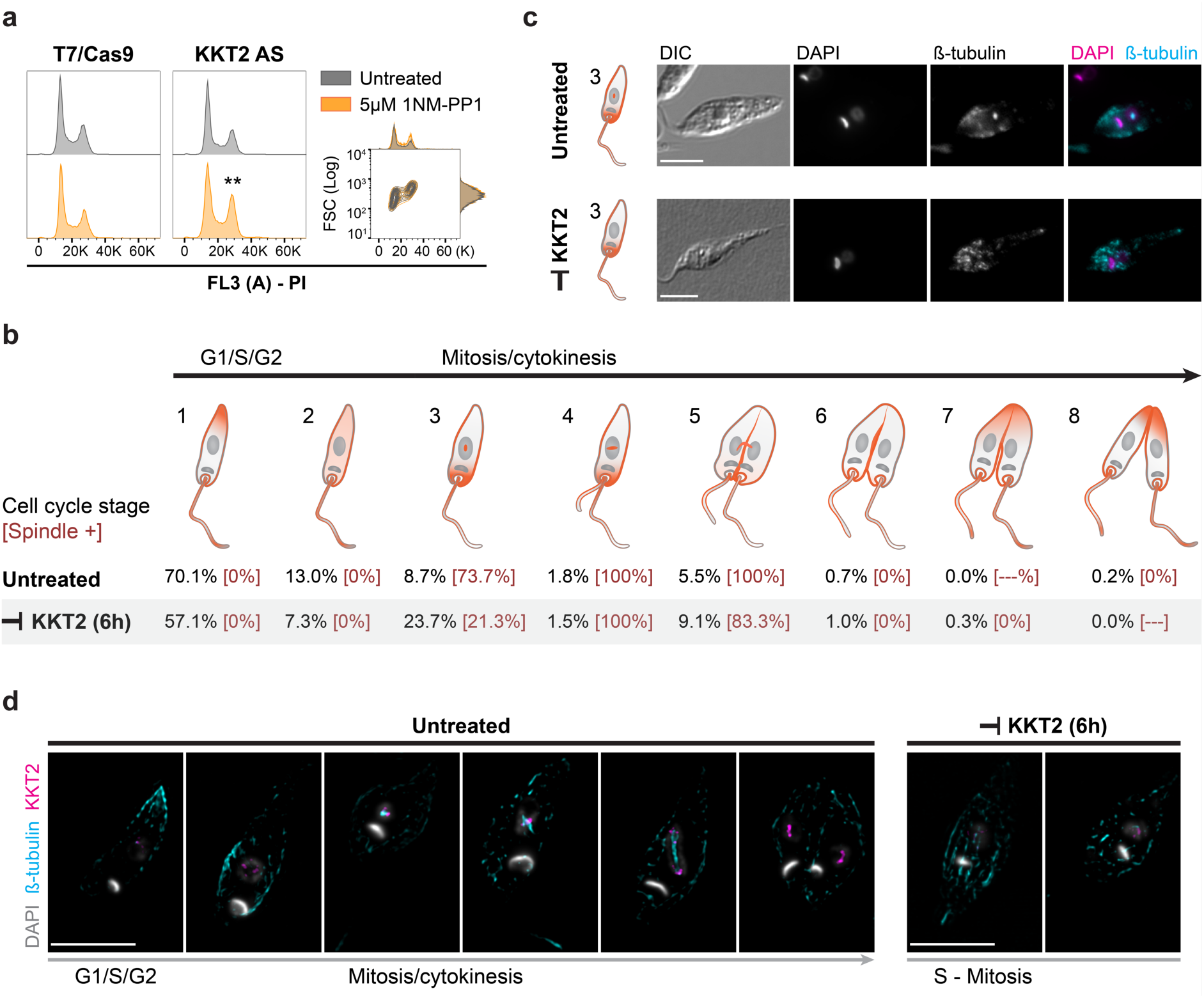
KKT2 kinase activity is required for cell cycle progression through synthesis and mitotic phases in *Leishmania*. (a) Representative cell cycle histograms of parasites stained with propidium iodide (PI) following 6 hours treatment with 5 µM 1NM-PP1. The parental T7/Cas9 line and untreated parasites expressing KKT2^M146G^ cultured under the same conditions were used as controls. Cell cycle phase quantification was performed using the Watson model algorithm in FlowJo v10.10.0. For statistical significance, p-values were calculated using two-tailed Student’s t-tests, comparing the percentage of cells in each phase between treated and untreated populations (**p < 0.01). Right panel: adjunct histograms showing DNA content and forward scatter (FSC) to assess cell size across different cell cycle phases. (b) Schematic representation of *L. mexicana* cell cycle stages based on nuclear and kinetoplast configurations (grey), flagellum number, β-tubulin (orange) distribution, as well as features of cytokinesis. The percentage of cells in each stage are given below with the proportion exhibiting mitotic spindles in square brackets (n = 396 and 438 cells for treated and untreated conditions, respectively). (c) Fluorescence microscopy of promastigote cells treated with 5 µM 1NM-PP1 for 6 hours, stained with the KMX-1 antibody to detect β-tubulin, and counterstained with DAPI to visualize DNA. Representative fluorescence micrographs show cells in stage 3 of the cell cycle (according to panel “b”) under treated and untreated conditions. Scale bars, 5 μm. DIC, the Nomarsky differential interference contrast. (d) Spatial and temporal distribution of *L. mexicana* KKT2 throughout the cell cycle. High-resolution fluorescence microscopy was performed on KKT2_AS::mNG::3xMyc promastigotes cultured with or without 10 µM 1NM-PP1 for 6 hours. Cells were stained with KMX-1 antibody to detect β-tubulin, anti-Myc antibody to visualize KKT2, and counterstained with DAPI for DNA visualization. Representative fluorescence micrographs show the localization of KKT2 and β-tubulin across different *Leishmania* cell cycle stages. The colours used for each marker are indicated to the left of the images. Scale bars, 5 μm. Additional individual and merged channel images are presented in Supplementary Fig 13.

To further investigate the effects of KKT2 inhibition on the *Leishmania* cell cycle, we employed fluorescence microscopy to monitor cell cycle progression by staining for β-tubulin (a mitotic spindle component) and DNA. This approach enabled us to classify *Leishmania* promastigotes into eight distinct cell cycle stages based on nuclear and kinetoplast configuration, flagellum number, the presence or absence of a mitotic spindle, β-tubulin distribution, and cytokinesis features (Table 2). Using this classification system, treatment of parasites expressing the KKT2 AS variant with 5 µM 1NM-PP1 for 6 hours led to an accumulation of cells exhibiting one nucleus, one kinetoplast, one flagellum, and β-tubulin concentrated in the anterior region of the cell – features characteristic of stage 3 (23.7% compared to 8.7% in the untreated control). However, unlike the untreated control in which the majority of stage 3 cells (73.7%) exhibited a visible mitotic spindle, only 21.3% of stage 3 cells under KKT2 inhibition displayed spindle formation, indicating that KKT2 activity is required either before or during mitotic spindle assembly (Fig 4b – c). In addition, a small increase in the proportion of cells in stage 5 was observed following KKT2 inhibition (9.1% compared to 5.5% in the untreated control), suggesting that KKT2 activity also plays a role in anaphase progression (Fig 4c).

**Table 2.**
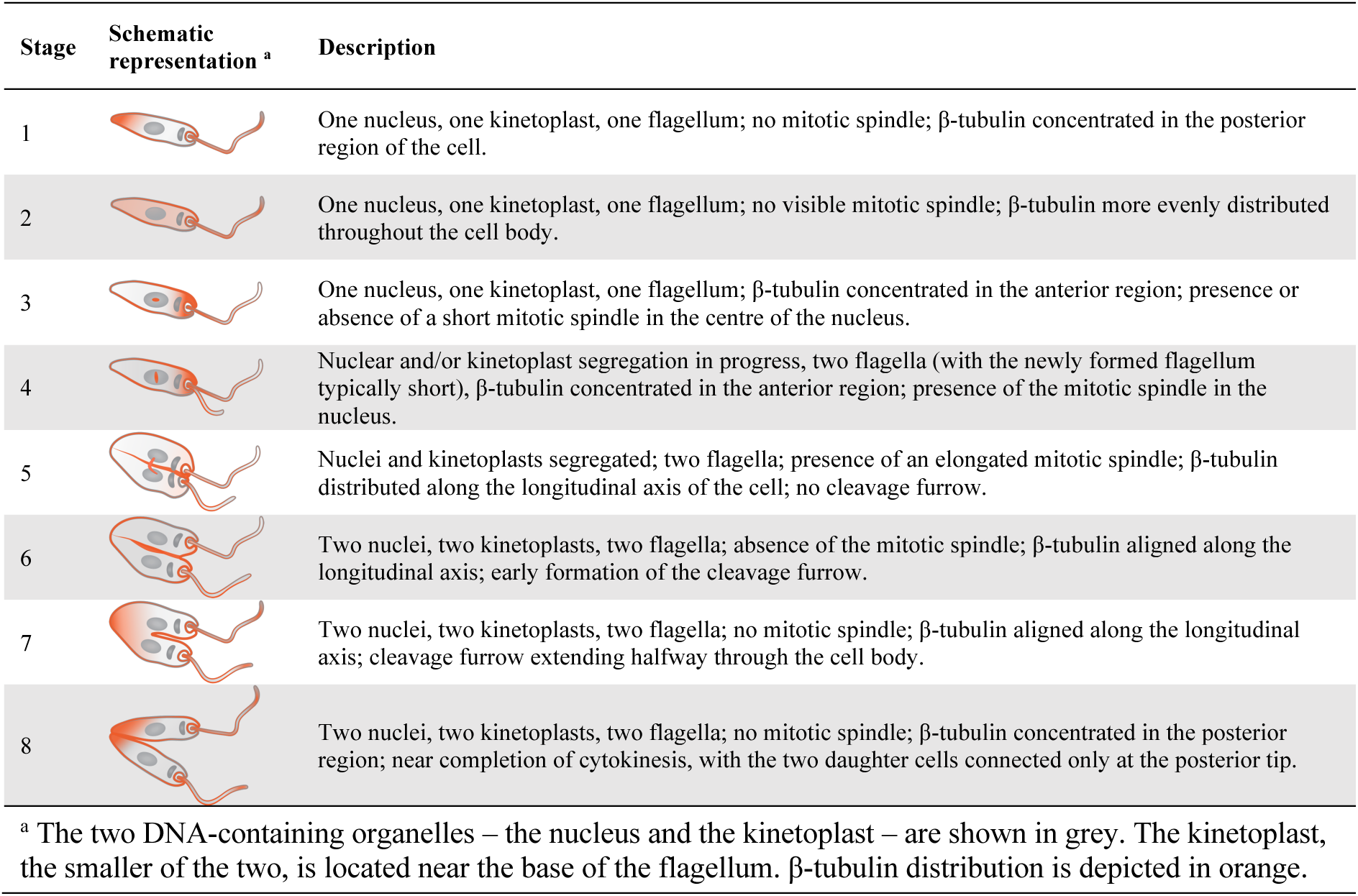
Definition of cell cycle stages.

| Stage | Schematic representation <sup>a</sup> | Description |
| --- | --- | --- |
| 1 |  | One nucleus, one kinetoplast, one flagellum; no mitotic spindle; β-tubulin concentrated in the posterior region of the cell. |
| 2 |  | One nucleus, one kinetoplast, one flagellum; no visible mitotic spindle; β-tubulin more evenly distributed throughout the cell body. |
| 3 |  | One nucleus, one kinetoplast, one flagellum; β-tubulin concentrated in the anterior region; presence or absence of a short mitotic spindle in the centre of the nucleus. |
| 4 |  | Nuclear and/or kinetoplast segregation in progress, two flagella (with the newly formed flagellum typically short), β-tubulin concentrated in the anterior region; presence of the mitotic spindle in the nucleus. |
| 5 |  | Nuclei and kinetoplasts segregated; two flagella; presence of an elongated mitotic spindle; β-tubulin distributed along the longitudinal axis of the cell; no cleavage furrow. |
| 6 |  | Two nuclei, two kinetoplasts, two flagella; absence of the mitotic spindle; β-tubulin aligned along the longitudinal axis; early formation of the cleavage furrow. |
| 7 |  | Two nuclei, two kinetoplasts, two flagella; no mitotic spindle; β-tubulin aligned along the longitudinal axis; cleavage furrow extending halfway through the cell body. |
| 8 |  | Two nuclei, two kinetoplasts, two flagella; no mitotic spindle; β-tubulin concentrated in the posterior region; near completion of cytokinesis, with the two daughter cells connected only at the posterior tip. |

<sup>a</sup> The two DNA-containing organelles – the nucleus and the kinetoplast – are shown in grey. The kinetoplast, the smaller of the two, is located near the base of the flagellum. β-tubulin distribution is depicted in orange.

To investigate the spatial and temporal distribution of KKT2 throughout the cell cycle and under conditions of kinase activity inhibition, we generated a tagged parasite line in which the C-terminus of both alleles of the KKT2 AS gene was endogenously fused to an mNeonGreen (mNG)::3xMyc epitope (KKT2_AS::mNG::3xMyc; Supplementary Fig 12). Subsequently, parasites were cultured in the presence or absence of 10 µM 1NM-PP1 for 6 hours, and KKT2 localization was examined using high-resolution fluorescence microscopy alongside β-tubulin and DNA staining to define cell cycle stages. When the KKT2 kinase is active in *Leishmania* parasites, the KKT2 protein exhibits a weak and diffuse signal during the early stages of the cell cycle, prior to mitotic spindle formation. As the cell cycle progresses – from spindle assembly to chromosome segregation – the KKT2 signal intensifies and becomes distinctly localized at the ends of the mitotic spindle. To investigate KKT2 distribution further, we analysed stage 3 cells lacking a mitotic spindle. In these cells, KKT2 appeared dispersed in the nucleus, in contrast to the untreated control at the same stage, where cells exhibited a developing mitotic spindle with KKT2 concentrated in close proximity to it (Fig 4d and Supplementary Fig 13).

Taken together, these results indicate that KKT2 kinase activity is required for key events during the coordination of synthesis and mitotic phases of the cell cycle in *Leishmania*. As these experiments were conducted in naturally asynchronous cell populations, which likely explains why the requirement for KKT2 activity was observed across multiple stages of the cell cycle.

### Inhibition of CRK9 kinase activity affects multiple stages of the *Leishmania* cell cycle

We have previously reported that inhibition of *L. mexicana* CRK9 kinase activity disrupts trans-splicing, which is predicted to lead to a widespread reduction in protein expression, resulting in broad cellular defects [43]. To investigate the role of CRK9 kinase activity in cell cycle progression, promastigotes expressing the AS variant of CRK9 were treated with 5 µM 1NM-PP1 for 6 hours. After treatment, a significant accumulation of cells in the G1 and G2/M phases was observed, along with a marked reduction in cells in S phase. Furthermore, CRK9 inhibition led to an increase in the proportion of cells in the Sub G0 phase, which is commonly associated with reduced DNA content, suggesting that CRK9 inhibition leads to DNA abnormalities (Fig. 5a and Supplementary Fig 14a). Notably, after 24 hours treatment, the significant accumulation in the Sub G0 phase persisted, while the other observed differences at the 6-hour time point were no longer present. The sustained increase in the Sub G0 phase after 24 hours indicates that the effects of CRK9 inhibition accumulate over time, leading to irreversible damage to the parasite (Supplementary Fig 14b).

**Fig 5.**
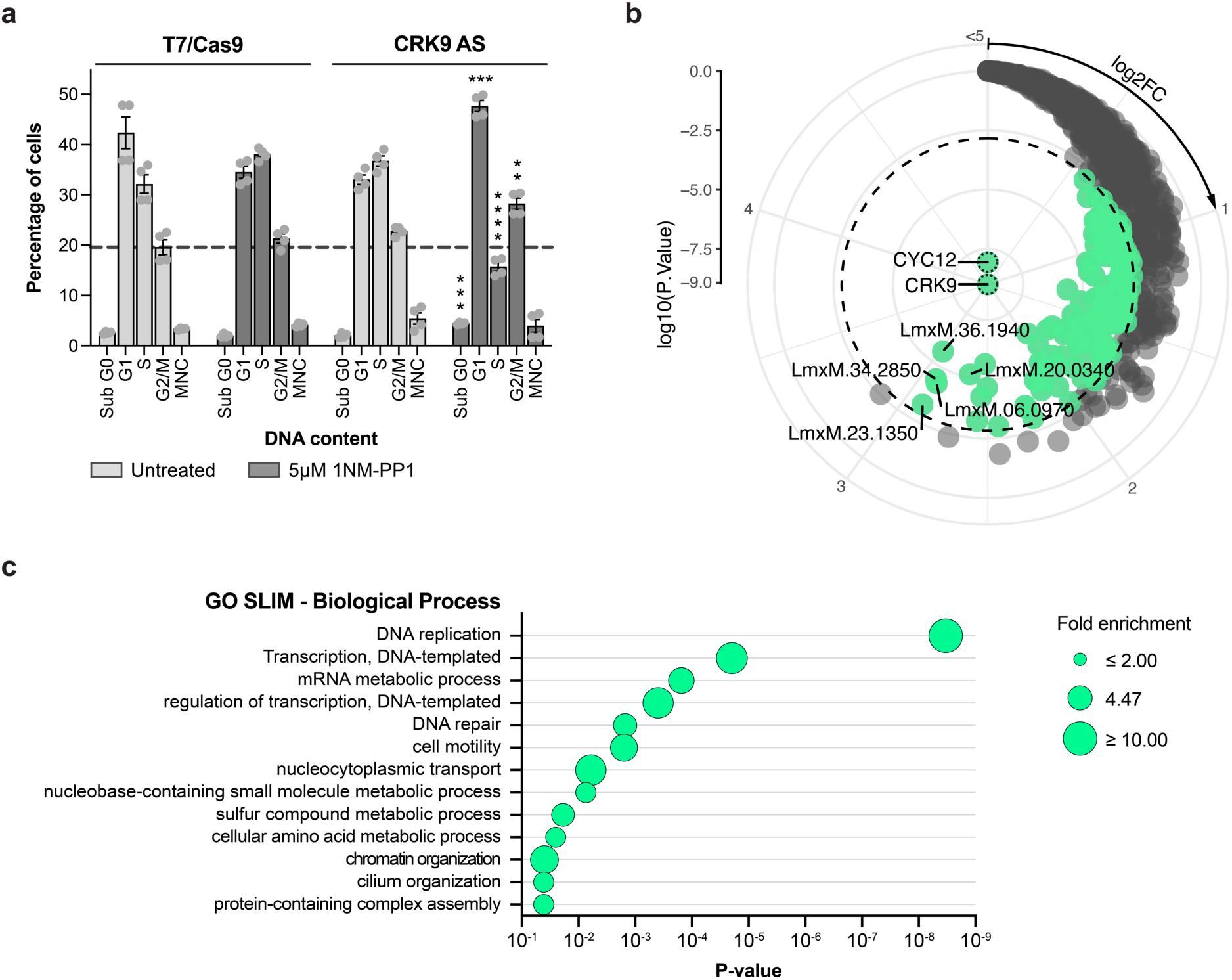
CRK9 protein kinase activity in the *Leishmania* cell cycle and its proximal proteome. (a) Effect of CRK9 inhibition on *Leishmania* cell cycle. Bar graph showing the percentage of cells in each cell cycle phase following 6 hours treatment with 5 µM 1NM-PP1. The parental T7/Cas9 line and untreated parasites expressing CRK9^M501G^ cultured under the same conditions were used as controls. Cell cycle phase quantification was performed using the Watson model algorithm in FlowJo v10.10.0. For statistical significance, p-values were calculated using two-tailed Student’s t-tests, comparing the percentage of cells in each phase between treated and untreated populations (**p < 0.01; ***p < 0.001; ****p < 0.0001). (b) CRK9-proximal proteins. Proximal proteins were determined with the limma package [46], with multiple testing correction performed according to Benjamini & Hochberg. Radial plot show proteins enriched in CRK9 compared to the spatial reference BDF7. The dashed line indicates a 1% false discovery rate (FDR) threshold used to distinguish proximal (green) from non-proximal (grey) proteins. Log2 fold enrichment increases in a clockwise direction; Log10 p-value increases from the centre outward. The six most enriched proteins (log2FC) are labelled in the plot: LmxM.36.5640, cyclin CYC12; LmxM.36.1940, inosine-guanosine transporter; LmxM.34.2850, RNA polymerase-associated protein LEO1; LmxM.23.1350, acetyltransferase-like protein; LmxM.06.0970, Domain of unknown function (DUF3883); LmxM.20.0340, hypothetical protein. (c) Core biological processes of the CRK9-proximal proteome. Bubble plot showing the significantly enriched Gene Ontology (GO) biological processes among CRK9-proximal proteins.

We applied our cross-linking BioID (XL-BioID) proximity labelling strategy to map the CRK9 proximal proteome and uncover potential functional pathways involving CRK9 [7, 43]. To this end, we generated 3xMyc::BirA*::CRK9 and the spatial reference 3xMyc::BirA*::BDF7 (Bromodomain Factor 7) cell lines using CRISPR-Cas9. A total of 137 statistically significant CRK9-proximal proteins were identified, including the previously reported CRK9 interactor, cyclin CYC12 [45] (Fig. 5b). Gene Ontology (GO) enrichment analysis revealed that the CRK9-proximal proteome is predominantly composed of proteins involved in DNA and RNA processing, consistent with CRK9’s role in gene expression regulation. While the exact mechanism for CRK9 regulation of trans-splicing is not fully established, it is postulated that it may not act directly on splicing factors and could lie upstream in a transcription-coupled process, given that splicing and polyadenylation occur co-transcriptionally in kinetoplastids [29, 45]. Indeed, we see enrichment of components of the Polymerase Associated Factor 1 complex (Leo1, CTR9), components that could be consistent with Elongator (Acetyltransferase-like, WD40-like), elongation factors (Spt5, TFIIS), and various components of RNA polymerase complexes (Supplementary Data 3). We also find components involved in DNA mismatch repair (MSH2, MSH3), PCNA and DNA polymerase delta, which could reflect a transcription-coupled mismatch repair pathway, or be involved in repairing damage cause by R-loops. The analysis also highlighted potential involvement of CRK9 in more distinct biological processes, such as cell motility, cilium organization, and amino acid metabolism (Fig. 5c) but these remain to be explored in detail.

### Identification of *Leishmania* protein kinases with targetable cysteines

Having exploited the gatekeeper residue to utilise the AS chemical genetics system in *Leishmania*, we decided to further explore the active site of *Leishmania* protein kinases using a broad-spectrum kinase inhibitor SM1-71. This probe acts by targeting the ATP-binding site and contains an acrylamide moiety that forms covalent adducts with cysteine residues. SM1-71 was originally developed to covalently bind cysteine residues immediately N-terminal to the DFG motif (DFG-1) in human kinases with its biotinylated analog, SM1-71-biotin (Supplementary Fig 15) enabling efficient capture on streptavidin-coated magnetic beads [36, 37].

We initially performed a label-free MS/MS analysis of proteins enriched from promastigote cell lysates incubated with either DMSO or SM1-71-biotin. This analysis identified 868 *Leishmania* proteins, which were consolidated into 802 unique entries based on sequence similarity for relative quantification. Among these, 131 proteins (121 entries) were significantly enriched in the SM1-71-biotin condition. Notably, 29 of the enriched proteins (28 entries) were annotated as protein kinases, and 38 entries were identified as other nucleotide-binding proteins. The enrichment of non-kinase proteins is likely attributed to the probe capturing hyper-reactive cysteines present throughout the proteome [47, 48]. Structural predictions using AlphaFold 3 revealed that 14 of these kinases possess cysteine residues in proximity to the DFG motif – 10 possess a cysteine at the canonical DFG-1 position, and 4 have cysteines located within seven residues of the motif. Additionally, 15 kinases were found to contain cysteine residues within the P-loop region (Supplementary Data 2 and Supplementary Fig 16).

To strengthen these findings, we applied a competitive chemoproteomic approach in which promastigote lysates were pre-treated with unlabelled SM1-71 (lacking biotin but including the acrylamide warhead) prior to incubation with SM1-71-biotin. This approach enabled the identification of more stable and specific probe-protein interactions. Through this strategy, 12 proteins were significantly enriched by SM1-71-biotin, including 11 protein kinases (Supplementary Data 2 and Supplementary Fig 16). Of these, 10 protein kinases overlapped with those identified in the initial enrichment assay. These consistently enriched kinases were classified as high-confidence candidates containing targetable cysteine residues amenable to covalent inhibition.

The sequence and structural prediction revealed the presence of a cysteine at the DFG-1 position in CLK1/CLK2 (C292 in CLK1, which differs from the cysteine C219 targeted by the covalent inhibitor AB1) [7], MPK4 (C173), MPK5 (C193), MPK7 (C272), and the protein kinase LmxM.32.1710 (C172). This DFG-1 cysteine is the most probable site of covalent interaction with the SM1-71 inhibitor, as the probe was specifically designed to target this region of the ATP-binding pocket [37]. Additionally, we observed enrichment of protein kinases that lack the DFG-1 cysteine but contain cysteines either in the P-loop region or in proximity to the DFG motif, sites previously shown to be reactive with the SM1-71 probe [36]. Among them are AEK1, which harbours cysteine C63 in the P-loop and C185 at the DFG-4 position; and the NEK family members LmxM.36.1520 (C52, C66, C98) and LmxM.36.1530 (C103), which contain cysteines in the P-loop region. Curiously, protein kinases STK36 and LmxM.19.0150 contain cysteines only in the C-lobe of the kinase domain, distant from the active site, suggesting the presence of available-reactive cysteines in these proteins. Five of these protein kinases with targetable cysteine residues in the active site were predicted to be essential for promastigote and/or amastigote survival: the kinetochore protein CLK1/CLK2, the mitogen-activated protein kinases MPK4, MPK5, and MPK7, and the AGC essential kinase AEK1 (Table 3 and Supplementary Fig 16).

**Table 3.** *L. mexicana* protein kinases with targetable cysteines.

| Accession | Kinase Group/Family | Kinase Name | Cysteine position in the kinase ATP-binding site <sup>a</sup> |  |  |  | Outside <sup>b</sup> |
| --- | --- | --- | --- | --- | --- | --- | --- |
|  |  |  | P-Loop | C-Helix | Hinge | DFG |  |
| LmxM.09.0400 * | CMGC/CLK | CLK1/KKT10 | C159[P4]<br>C210[P6] |  | **C219** | **C292[DLG-1]**<br>C297[DLG+2]<br>C298[DLG+3] |  |
| LmxM.13.0440 | Other/ULK | STK36 |  |  |  |  | C192[C Lobe]<br>C218[C Lobe] |
| LmxM.13.1640 * | CMGC/MAPK | MPK7 |  |  |  | C267[DFG-6]<br>**C272[DFG-1]** |  |
| LmxM.19.0150 | STE |  |  |  |  |  | C105 |
| LmxM.19.1440 * | CMGC/MAPK | MPK4 | **C41[P2]**<br>C45[P3] |  |  | C168[DFG-6]<br>**C173[DFG-1]** |  |
| LmxM.25.2340 * | AGC | AEK1 | C63[P3] |  |  | C185[DFG-4] |  |
| LmxM.29.2910 * | CMGC/MAPK | MPK5 | C58[P3] | C100[+1]<br>C102[+2] |  | **C193[DFG-1]** |  |
| LmxM.32.1710 | CAMK/CAMKL |  |  |  |  | **C172[DFG-1]** |  |
| LmxM.36.1520 | NEK |  | C52[P3]<br>C66[P4]<br>C98[P5] | C86<br>C90[+2] |  |  | C179<br>C208<br>C222 |
| LmxM.36.1530 | NEK |  | C103[P5] | C90<br>C91<br>C95[+2] |  |  | C184<br>**C213** |

\* *L. mexicana* protein kinase required for promastigote and/or amastigote survival (Supplementary Data 1). <sup>a</sup> The positions of cysteine residues surrounding the ATP-binding site were identified using AlphaFold 3 structural prediction and visualised with ChimeraX v1.9. The ribbon-structure representation of cysteine locations is shown in Supplementary Fig 16. Cysteines located within 6 Å of the ATP ligand are highlighted in bold. <sup>b</sup> Outside, cysteine located outside the ATP-binding site.

## Discussion

Owing to their well-established druggability, protein kinases are among the most intensively investigated targets in pharmaceutical research, with over 80 protein kinase inhibitors approved by the U.S. Food and Drug Administration (FDA) for the treatment of various cancers and other diseases [12, 49]. Recently we genetically validated over 40 protein kinases from the *Leishmania* kinome as they were found to be required for parasite survival [14]. Kinase druggability has also been demonstrated in *Leishmania*, where a pyrazolopyrimidine compound was shown to selectively inhibit CRK12, a member of the CMGC kinase family [5], and the amidobenzimidazole compound AB1 was found to specifically target the kinetochore-associated kinases CLK1/CLK2 [7].

Our kinome-wide bioinformatics analysis of *L. mexicana* revealed the absence of protein kinases with naturally occurring glycine or alanine residues at the gatekeeper position. This finding stands in contrast to several apicomplexan parasites, including *Toxoplasma gondii* [50], *Cryptosporidium parvum* [51], and *Neospora caninum* [52], where glycine residues are found at the gatekeeper position in calcium-dependent protein kinase 1 (CDPK1), thereby enabling selective inhibition by bumped kinase inhibitors and supporting their development in a targeted therapeutic approach. *Leishmania* does not contain this class of calcium-dependent protein kinases. Motivated by the absence of naturally occurring AS-compatible kinases in *L. mexicana*, we adopted a chemical-genetic strategy to conditionally interrogate kinase function. Notably, the smallest gatekeeper residue identified in our kinome analysis was a serine in the CMGC family kinase LmxM.36.4250. This kinase is implicated in amastigote survival in an *in vivo* mouse model, but not in promastigotes or *in vitro* macrophage-internalized amastigotes. This observation is relevant given prior evidence that naturally occurring serine gatekeepers can confer sensitivity to BKIs, as shown for MAPKL-1 (mitogen-activated protein kinase like 1) in *T. gondii* [53] and CDPK4 in *Plasmodium spp* [54]. These findings have important implications for the application of AS kinase technology in *Leishmania*. While *L. mexicana* represents a suitable model for in-cell target validation using AS kinase approaches in promastigote and *in vitro* macrophage-internalized amastigotes, the presence of the natural AS-serine LmxM.36.4250 kinase may confound interpretation in *in vivo* models.

We selected four kinetochore-associated kinases, CLK1/CLK2, KKT2 and KKT3, along with the cyclin-dependent kinase CRK9, for chemical-genetic validation of their kinase activity requirements in *Leishmania* using our inducible AS kinase system. We were unable to generate CLK1/CLK2 AS mutants, probably because the mutation is not tolerated by the kinase. Through this approach, we demonstrated that the kinase activities of KKT2 and CRK9 are essential for parasite survival in both the promastigote stage and, critically, in the clinically relevant intracellular amastigote stage. These findings position KKT2 and CRK9 as promising therapeutic targets for the development of anti-leishmanial kinase inhibitors. Notably, KKT2 lacks orthologs in the human kinome [55], indicating a greater potential for achieving selectivity in drug development. In contrast, CRK9 has sequence similarity to many human CDKs, with CDK2 showing the highest sequence identity (37%; Supplementary Fig 17). Supporting our findings, CRK9 kinase activity has previously been validated as essential for parasite survival in a murine model of *T. brucei* infection [45]. Importantly, we also showed that the in-cell AS kinase strategy is compatible with the intra-macrophage amastigote stage, enabling conditional chemical modulation of kinase activity within an intracellular environment. This represents a significant advance, as direct genetic manipulation of intracellular amastigotes remains technically challenging, limiting functional assessment of kinase essentiality for parasite survival.

In contrast, AS-mediated inhibition of KKT3 did not result in any observable growth defect in *Leishmania*, despite our gene deletion data suggesting KKT3 as an essential gene. These findings align with previous work in *T. brucei*, where deletion of the protein kinase domain showed that KKT3 kinase activity is dispensable for proliferation of procyclic cells [56], whereas RNAi-mediated knockdown of the full-length protein indicated that KKT3 is required for parasite survival [55, 57]. Here, we cannot exclude the possibility that the gatekeeper residue substitution in AS-KKT3 failed to sufficiently expand the ATP-binding pocket to permit effective binding of bumped kinase inhibitors. Therefore, additional investigation is necessary before concluding that KKT3 kinase activity is dispensable for *Leishmania* survival.

We leveraged our inducible AS kinase system to investigate the functional roles of KKT2 and CRK9 in *Leishmania*, aiming to gain mechanistic insights into the biology of these proteins. KKT2 is a non-canonical protein kinase and a core component of the kinetoplastid kinetochore [55] – a specialized macromolecular complex that, together with centromeres and spindle microtubules, drives chromosome segregation in eukaryotes [58]. In *T. brucei*, RNAi-mediated depletion of KKT2 has been shown to impair proliferation in both procyclic and bloodstream forms, and to disrupt the localization of the kinetochore protein KKT14 [57, 59]. Subsequent studies demonstrated that KKT2 kinase activity is specifically required for proliferation in the bloodstream stage [38] and for accurate chromosome segregation in the procyclic form [56]. However, KKT2 kinase activity was found to be dispensable for the localization of KKT2 itself, as well as for the proper localization of the kinetochore proteins KKT1, KKT9, and KKT14 [38, 56]. KKT2 is a multidomain protein and its unique centromere localization domain in the middle, and its divergent polo-box domain at the C-terminus have been shown to be sufficient for its correct localization [59, 60]. Our findings revealed that KKT2 kinase activity is essential for mitotic progression in *L. mexicana*, as evidenced by an increased proportion of cells arrested at an early mitotic stage (stage 3) lacking a detectable mitotic spindle upon inhibition of AS mutant. This observation suggests that KKT2 activity may be required for mitotic spindle assembly. However, given that KKT2 function has primarily been linked to kinetochore assembly [38, 55, 59], and that we did not directly assess the impact of KKT2 inhibition on kinetochore assembly, we cannot exclude the possibility that the spindle defects observed are secondary to impaired kinetochore function resulting from loss of KKT2 activity. In addition, we demonstrate that KKT2 activity is required for proper coordination of S-phase progression, providing new insights into the regulation of cell cycle transitions in kinetoplastids. We found that KKT2 is constitutively expressed throughout the *Leishmania* cell cycle and displays a punctate nuclear signal. Prior to mitosis, KKT2 exhibits a more diffuse nuclear distribution, whereas during mitosis it becomes concentrated at the ends of the mitotic spindle, suggesting a spatially dynamic role linked to mitotic progression.

To further dissect essential regulatory nodes in the *Leishmania* cell cycle, we investigated CRK9, a member of the cyclin-dependent kinases, which play central roles in regulating cell cycle progression and gene expression in eukaryotes [61]. Although trypanosomatid cyclin-dependent kinases retain a conserved domain architecture, their primary sequences are divergent, making it difficult to unambiguously identify their counterparts in human and yeast [13]. Like other cyclin-dependent kinases, CRK9 functions in complex with a cyclin (CYC) partner, CYC12, an L-type cyclin, and together with CRK9-associated protein (CRK9AP), forms an unusual tripartite complex [45]. Although CRK9AP was not detected in our proximity labelling dataset, CYC12 was the most enriched CRK9-proximal protein. In *T. brucei*, CRK9 has been implicated in the regulation of pre-mRNA processing [62], an effect later shown to depend directly on its kinase activity [29]. Consistent with this role, our previous work demonstrated that inhibition of CRK9 kinase activity in *Leishmania* leads to accumulation of unspliced pre-mRNAs, indicating a conserved role in RNA processing among kinetoplastids [43]. In the present study, we found significant enrichment of CRK9-proximal proteins involved in DNA and RNA processing pathways. Furthermore, we show that AS-mediated inhibition of CRK9 in *Leishmania* results in broad disruption of cell cycle progression, marked by accumulation of cells in Sub-G0, G1, and G2/M phases, and a corresponding reduction in S-phase cells. This pleiotropic phenotype is consistent with the expected consequences of inhibition of trans-splicing, which disrupts global mRNA maturation and protein synthesis, ultimately leading to broad cellular dysfunction. Notably, the increase in the Sub-G0 population, often associated with DNA fragmentation, persisted after 24 hours of treatment, suggesting that CRK9 inhibition may lead to DNA content dysregulation. Consistent with our findings, RNAi-mediated depletion of CRK9 in *T. brucei* has been shown to impair mitosis and cytokinesis in the procyclic form, although no significant impact on proliferation was observed in the bloodstream stage [63].

In addition, our kinome-wide bioinformatic analysis of gatekeeper residues also identified alternative opportunities for chemical-genetic intervention. Notably, we found that the protein kinase LmxM.36.0910, a member of the STE kinase family, harbours a cysteine at the gatekeeper position – a feature that renders it susceptible to covalent inhibition by electrophilic compounds specifically designed to target gatekeeper cysteines [64]. This discovery provides a potential route to functionally interrogate LmxM.36.0910 using covalent chemical tools, however, functional genetics analyses revealed that this kinase is non-essential across all life cycle stages of *Leishmania* parasite, indicating limited therapeutic relevance despite its biochemical tractability.

In the context of covalent chemical tools, our chemoproteomic profiling using the electrophilic probe SM1-71 identified five essential kinases, CLK1/CLK2, MPK4, MPK5, MPK7, and AEK1, that harbour reactive cysteine residues amenable to covalent inhibition. Covalent kinase inhibitors represent an emerging class of therapeutics, with eleven FDA-approved compounds employing irreversible targeting of cysteine residues through Michael addition chemistry [49, 65]. In line with this strategy, our previous work demonstrated that the amidobenzimidazole compound AB1 covalently binds to a hinge-region cysteine in CLK1 of trypanosomatids [6, 7]. These findings support the further validation of MPK4, MPK5, MPK7, and AEK1 as promising new candidates for the development of covalent kinase inhibitors in *Leishmania*.

To enable functional interrogation of protein kinase activity in *Leishmania*, in this study, we successfully adapted a CRISPR-Cas9-based genome editing strategy to introduce single-codon substitutions directly into endogenous *loci* without the use of drug selection markers. Our marker-free approach employed 120-nt single-stranded oligonucleotide donor DNA containing short homology arms, the desired target mutation, and silent shield mutations to prevent further Cas9-mediated cleavage. Two sgRNAs flanking the target codon were used in our strategy. Using this system, we achieved an average editing efficiency of 11.7% for the successful substitution of the codon encoding the gatekeeper residue in both alleles of *KKT2*, *KKT3*, and *CRK9*, demonstrating that exhaustive screening was not required to isolate correctly edited clones for these genes. Comparable approaches have been reported in other kinetoplastids. In *T. brucei*, a similar strategy was used to introduce point mutations into the aquaglyceroporin gene (AQP2) using a 48-nt single-stranded oligonucleotide repair template and a sgRNA expressed from a vector, yielding an editing efficiency of approximately 8% [66]. In *T. cruzi,* CRISPR-Cas9-mediated editing using single-stranded oligonucleotide DNA donors has also been used to introduce premature stop codons, although this system relied on the delivery of recombinant Cas9 ribonucleoprotein complexes rather than endogenous Cas9 expression [67]. In *Leishmania*, marker-free genome editing was first demonstrated in *L. donovani*, where a single-stranded oligonucleotide was used to introduce a stop codon into the miltefosine transporter gene, which conferred resistance to miltefosine [68]. Unlike our strategy, in that system the sgRNA was expressed from an episomal vector and clones carrying the desired mutation were enriched by miltefosine selection. A subsequent adaptation of this approach employed a co-targeting strategy, in which the miltefosine transporter gene was simultaneously targeted to allow for selection-based enrichment of edited cells. This co-selection significantly improved editing efficiency and enabled the deletion of multicopy gene families in *L. donovani* [69]. Furthermore, a marker-free CRISPR-Cas9 editing strategy was described in *L. major* to introduce a single amino acid substitution in the calcium-dependent kinase SCAMK, with both the sgRNA and DNA donor delivered via an episomal expression vector [70]. Unlike these systems, our approach is fully compatible with the high-throughput CRISPR-Cas9 toolkit developed by Beneke et al. [42], which enables streamlined genome editing without the need for custom vector construction. This compatibility, combined with the absence of selection markers, makes our strategy well-suited for scalable functional genomic studies in *Leishmania*.

Altogether, this study establishes a powerful framework for dissecting protein kinase function in *Leishmania*, integrating chemical-genetic tools with CRISPR-mediated precise genome editing to enable in-cell interrogation of targeted kinases. Through this approach, we identify KKT2 and CRK9 as promising therapeutic targets for antileishmanial drug development. Notably, the evolutionary conservation of these protein kinases across clinically relevant trypanosomatid species raises the prospect of developing a single therapeutic agent with efficacy against multiple kinetoplastid diseases, including leishmaniasis, Chagas disease, and African trypanosomiasis.

## Materials and Methods

### Cell culture

*Leishmania mexicana* promastigotes (strain MNYC/BZ/62/M379) were cultured at 25°C in HOMEM medium (modified Eagle’s medium; Gibco, ThermoFisher Scientific) supplemented with 10% heat-inactivated fetal calf serum (hi-FCS) (Gibco, ThermoFisher Scientific) and 100 U penicillin – 100 μg mL^-1^ streptomycin (Sigma-Aldrich), pH 7.2. Where applicable, selective antibiotics were added at the following concentrations: Hygromycin B (InvivoGen, ant-hg) at 50 μg mL^-1^; Nourseothricin (Jena Bioscience, AB-101) at 50 µg mL^-1^; Blasticidin (InvivoGen, ant-bl) at 10 μg mL^-1^; Puromycin (InvivoGen, ant-pr) at 30 μg mL^-1^. Bone marrow-derived macrophages (BMDM) isolated from BALB/c mice were differentiated in DMEM (Dulbecco’s Modified Eagle Medium) medium supplemented with 10% hi-FCS and macrophage colony-stimulating factor secreted by L929 cells [71]. BMDM was maintained in culture in DMEM medium supplemented with 10% hi-FBS and 10 mM L-glutamine (Gibco, ThermoFisher Scientific) at 37°C, in an atmosphere of 5% CO_2_. All experiments were conducted according to the Animals (Scientific Procedures) Act of 1986, United Kingdom, and had approval from the University of York Animal Welfare and Ethical Review Body (AWERB) committee.

### Growth curve

*L. mexicana* promastigotes were inoculated at a density of 4×10^4^ parasites mL^-1^ in HOMEM medium supplemented with 10% hi-FCS. The cumulative cell growth was monitored daily by manual counting using a Neubauer hemocytometer. Growth rate calculations were performed using data from the logarithmic phase of the growth curve (0 – 96 h).

### Identification of kinase gatekeeper residues in the *Leishmania* kinome

The kinase domains of the 216 predicted protein kinases in the *L. mexicana* genome were identified using InterPro domain analysis [72]. The gatekeeper residue was identified by multiple sequence alignment of each kinase family. Where the alignment was ambiguous, or an unusual residue was predicted, the gatekeeper was confirmed by 3D structure comparison with a kinase of known structure and a well-defined gatekeeper (eg. PDB code 3F3W). When a suitable structure model (e.g. of its paralog from *L. infantum*) for the kinase in question was not available in the AlphaFold database, a model was generated using AlphaFold 3 [41]. Structures were aligned using secondary structure matching [73] and the superposition was checked by confirming the overlap of conserved motifs around the active site Lys and the conserved Asp residues in the sequence motifs HRD and DFG. The gatekeeper residue was identified as being at the end of a β-strand immediately before the hinge region that separate the N- and C-lobes of the protein kinase. The predicted models were visualized in ChimeraX v1.9.

### *L. mexicana* genome editing by CRISPR-Cas9

To achieve precise genome editing, we utilized the CRISPR-Cas9 system in *L. mexicana* strain MNYC/BZ/62/M379, which constitutively expresses Cas9 and T7 RNA polymerase, as previously described [42, 74]. To introduce single-codon mutations without the use of drug selection markers, two single-guide RNAs (sgRNAs) targeting sequences flanking the codon of interest was employed. The sgRNAs were designed using the Eukaryotic Pathogen CRISPR guide RNA/DNA Design Tool (http://grna.ctegd.uga.edu) with default parameters (SpCas9: 20 nt gRNA, NGG PAM on 3’end). Primers for sgRNA synthesis were manually designed to align with the high-throughput system outlined by Beneke et al. [42]. The sgRNA templates were generated via PCR using a forward oligonucleotide encoding the T7 promoter sequence (5’-GAAATTAATACGACTCACTATAGG-3’), followed by 20 nt corresponding to the target-specific guide sequence and the complementary sequence (5’-GTTTTAGAGCTAGAAATAGC-3’) to the 3’ end of the reverse oligonucleotide (OL6137), which contain sgRNA Cas9 scaffold sequence [42]. The resulting PCR products were purified using the QIAquick PCR Purification Kit (Qiagen, Cat. 28106) and prepared in ultra-pure water at a final concentration of ∼1 µg µL^-1^. The DNA repair template (DRT) used to introduce gatekeeper mutations in kinases were designed as 120 nt single-stranded DNA oligonucleotide. Each template contained the desired codon mutation, silent mutations at the protospacer adjacent motif (PAM) and sgRNA binding sites to prevent re-cutting, and 20 – 30 nucleotides of homology flanking both sides of the Cas9-induced double-strand break (DSB). The single-stranded DRT were resuspended in nuclease-free water at a final concentration of 2.2 µg µL^-1^. Approximately 5 µg of sgRNA and 11 µg of DRT were combined and transfected into 5×10^6^ promastigote-stage of the *L. mexicana* cell line stably expressing T7 RNA polymerase and Cas9 endonuclease (T7/Cas9) [42]. Transfections were performed using the P3 Primary Cell 4D-Nucleofector Kit (Lonza, Cat. V4XP-3024) with a single pulse of program FI-115 in a final volume of 110 µL. Post-electroporation, cells were immediately transferred to pre-warmed HOMEM medium supplemented with 20% hi-FCS and 10 µM 6-biopterin. After a 16 hours recovery period, transfected cells were cloned into 96-well plates at a ratio of one cell per two wells.

Generation of gene knockout or endogenously tagged *Leishmania* lines was carried out using the CRISPR-Cas9 genome editing toolkit for kinetoplastids developed by Beneke et al. [42]. Primers for sgRNA and repair template construction were designed using the LeishGEdit platform (http://leishgedit.net/). For generation of null knockout mutants, two distinct repair templates – differing only in their drug resistance markers – were employed. Parasites were transfected with purified sgRNA and corresponding DNA repair templates as described above. Sixteen hours post-transfection, selective drugs (blasticidin and/or puromycin) were added, and lines were cloned by limiting dilution. Genomic DNA from recovered clones was extracted, and diagnostic PCRs were performed, followed by restriction enzyme digestion where required. Digestions were incubated for 16 h using the appropriate restriction enzymes and buffers recommended by New England Biolabs. Oligonucleotide sequences used to generate and validate CRISPR-Cas9-edited lines are provided in Supplementary Tables 1 – 3.

Null mutants were validated by whole-genome sequencing using the Illumina platform. Paired-end reads (150 bp) were aligned to the *L. mexicana* MNYC/BZ/62/M379 Cas9/T7 reference genome [74], using Minimap2 v2.26-r1175. Read alignments were visualized with IGV v2.16.1. Genome coverage was assessed using Mosdepth v0.3.3, which calculated read depth in 500 bp windows and mean coverage across all chromosomes. Gene coverage was also determined by Mosdepth based on annotations from the MNYC/BZ/62/M379 Cas9/T7 reference. Sample coverage was normalised per chromosome, and the coverage change was calculated from the relative difference between the comparison and reference. 0.5 was used as a cut off, which equates to one copy change. The Illumina sequencing data were deposited under the SRA Bioproject accession number PRJNA1303394.

### *In vitro* susceptibility assay of *Leishmania* promastigotes

A dose response curve was set in a 96-well plate with 1×10^4^ parasites mL^-1^ treated with two-fold increasing concentrations of the bumped kinase inhibitor (1NA-PP1 or 1NM-PP1). The viability of treated and untreated control was assessed after 96 hours by addition of 50 µL of 0.0125% (w/v) resazurin (Alamar Blue) prepared in PBS. Cells were incubated for an additional 2 – 4 hours at 37°C, after which fluorescence was measured using a CLARIOstar^®^ plate reader (BMG LABTECH) with excitation at 540 nm and emission at 590 nm. Fitting of dose-response curves and IC_50_ calculation were carried out using GraphPad Prism v9.3.1, with viability normalized to untreated controls (set as 100%) for each cell line.

### *In vitro* susceptibility assay of intracellular *Leishmania* amastigote

Promastigotes were cultured in HOMEM medium supplemented with 10% hi-FCS until reaching late-logarithmic phase. Differentiated BMDM were plated on 16 well Labtek tissue culture slides (Nunc, NY, USA) and then infected at a ratio of 10 promastigotes per macrophage. After 18 hours incubation at 37°C in 5% CO₂ using DMEM supplemented with 5% hi-FCS, non-internalized promastigotes were removed by washing, and infected macrophages were treated with serial dilutions of either BKIs 1NA-PP1 or 1NM-PP1 prepared in DMEM supplemented with 2% heat-inactivated horse serum. Following 96 hours treatment, cells were fixed with methanol and stained with Giemsa. Infected cells were quantified by light microscopy using a Zeiss Axiolab-5 microscope, with 100 macrophages counted per well to determine the percentage of infected macrophages. Parasite viability was calculated as the percentage of infected macrophages in treated wells relative to untreated controls. Fitting of dose-response curves and IC_50_ calculation were carried out using GraphPad Prism v10.1.0, with viability normalized to untreated controls set as 100%.

### Cell cycle analysis

Promastigote cells were cultured in the presence or absence of 5 µM 1NM-PP1 for 6 or 24 hours. Following treatment, cells were washed with PBS containing 5 mM EDTA (PBS-EDTA) and resuspended in 70% methanol. After overnight incubation at 4°C, cells were washed once with PBS-EDTA and resuspended in 1 mL PBS-EDTA containing 10 µg mL^-1^ of propidium iodide and 10 µg mL^-1^ of RNase A. Following a 45-minute incubation at 37°C in the dark, DNA content was analysed by flow cytometry using a CyAn^TM^ ADP cytometer (Beckman Coulter). Cell cycle distribution was determined using the Watson model in FlowJo^TM^ v10.6.2 software.

### Immunofluorescence microscopy

Promastigote cells growing in the presence or absence of 5 or 10 µM 1NM-PP1 for 6 or 24 h were washed twice with PBS (1,400 g for 10 minutes at room temperature). Approximately 10^6^ were resuspended in PBS and allowed to adhere for 15 minutes at 37°C onto poly-L-lysine-coated high-precision coverslips (thickness No. 1.5H [0.170 mm ± 0.005 mm], MARIENFELD: cat. 0107222). *In vivo* cross-linking was performed by incubating the adhered cells with 1 mM disuccinimidyl suberate (DSS) in PBS for 10 minutes at 37°C. Cells were fixed at room temperature with 4% paraformaldehyde in PBS for 15 minutes, followed by quenching with 0.1 M glycine in PBS (pH 7.6) for 5 minutes. After two washes with PBS, cells were permeabilized with 0.5% Triton X-100 in PBS for 15 minutes. Blocking was performed by incubating cells in blocking buffer (5% BSA, 0.01% saponin in PBS) for 1 hour at room temperature. Primary immunostaining was carried out for 1 hour at room temperature using mouse anti-β-tubulin KMX-1 antibody (Sigma-Aldrich, MAB3408) diluted 1:1000 in blocking buffer. After three washes with 0.1% Triton X-100 in PBS, cells were incubated for 1 hour with Alexa Fluor^TM^ 647-conjugated goat anti-mouse IgG secondary antibody (Abcam ab150119) diluted 1:1000 in blocking buffer. Following three washes with 0.1% Triton X-100/PBS, cells were counterstained with 20 µg mL^-1^ DAPI in PBS for 30 minutes, followed by a final PBS wash. Coverslips were mounted on glass slides using ProLong diamond antifade mountant (Invitrogen™), according to the manufacturer’s instructions.

For detection of endogenously tagged KKT2, the above protocol was followed using the following antibody combinations: mouse anti-β-tubulin KMX-1 [diluted 1:800], and rabbit anti-Myc-Tag (clone 71D10, Cell Signaling Technology mAb #2278) [diluted 1:200] as primary antibodies; and Alexa Fluor™ 568-conjugated goat anti-mouse IgG (Invitrogen A-11031) [diluted 1:800], and Alexa Fluor™ 488-conjugated donkey anti-rabbit IgG (Invitrogen A-21206) [diluted 1:200] as secondary antibodies.

### Microscopy and image analysis

Widefield fluorescence imaging was performed using a Zeiss Axio Observer7 microscope in z-stack mode (20 optical sections were captured per field). Images were processed using Fiji v2.14.0/1.54f, employing the Microvolution blind deconvolution module. Maximum intensity projections were subsequently generated using selected z-stack layers retrieved with the “z-project” function in Fiji.

Super-resolution structured illumination microscopy (SR-SIM) was performed on a Zeiss Elyra 7 system using the Lattice SIM modality. 3D acquisition in 3 colour was performed [DAPI Excitation 405nm Emission 420-480nm; AF488 Excitation 488nm Emission 490-560nm; AF568 Excitation 561nm Emission 570-630nm] with a z-stack of 50 slices captured at 0.091 µm intervals. Zeiss Zen Black v3.0 software was used to reconstruct the images by SIM^2^ processing using different pre-set processing parameter depending on the fluorescent signal intensities [fixed standard for β-tubulin_AF568 and KKT2_Myc_AF488; and low contrast for DAPI]. To ensure accurate colour alignment, SR-SIM images were taken of TetraSpeck fluorescent microspheres with 200 nm diameter (Thermo Fisher Scientific). Then the Zen Black Channel Alignment module was used to calculate a correction matrix that was applied to the experimental images.

### CRK9-proximal proteins affinity purification

The cell lines *L. mexicana* T7/Cas9 3xMyc::BirA*::BDF7, 3xMyc::BirA*::CRK9 were generated using Cas9 directed endogenous tagging with pPlot BirA* Puro [42]. Cultures in 100 mL HOMEM 10% FBS were set up in quadruplicate and grown until early log stage, approximately 3×10^6^ cells mL^-1^, at this point d-biotin was supplemented to the media at 150 µM for 18 hours at 25°C. After biotinylation cells were harvested by centrifugation (1,500 g for 10 minutes) washed twice in PBS then resuspended in pre-warmed PBS at a density of 4×10^7^ cells mL^-1^. Cells were treated with 1 mM DSP crosslinker (Thermo) for 10 minutes at 25°C; this crosslinking was quenched with Tris-HCl pH7.5 to a concentration of 20 mM. The cells were then collected by centrifugation and stored at −80 °C until lysis. Samples were lysed with 500 µL ice cold RIPA buffer (25 mM Tris-HCl pH 7.6, 150 mM NaCl, 1% NP-40, 1% sodium deoxycholate, 0.1% SDS) containing 2x HALT protease inhibitor cocktail (Thermo) and 1x PhosSTOP (Roche). To each tube of lysate, 1µL of BaseMuncher Endonuclease (250 units, Abcam) was added and nucleic acids digested at room temperature for 10 minutes. The samples were then sonicated using a BioRuptor Pico (3x cycles, 30 seconds on, 30 seconds off, 4°C) and clarified by centrifugation in Protein LoBind tubes (Eppendorf) at 10, 000 g for 10 minutes at 4 °C. Biotinylated proteins were then enriched using 100 μL of magnetic streptavidin bead suspension (1 mg of beads, ResynBioscience) for each affinity purification from 4×10^8^ parasites. Binding was performed overnight at 4°C with end-over-end rotation. Beads were then washed in 500 μL of the following buffers for 5 min each: RIPA for 4x washes; 4 M urea in 50 mM triethyl ammonium bicarbonate (TEAB) pH 8.5; 6 M urea in 50 mM TEAB pH 8.5; 1 M KCl, 50 mM TEAB pH 8.5. Beads from each affinity purification were then resuspended in 200 μL 50 mM TEAB pH 8.5 containing 0.01% ProteaseMAX (Promega), 10 mM TCEP, 10 mM Iodoacetamide, 1 mM CaCl_2_ and 500 ng Trypsin Lys-C (Promega). Digest was conducted overnight in a 37°C shaking heat block, at 900 rpm. The treatment with reducing agent in this step also cleaves the DSP crosslinker. The supernatant was recovered from the beads which were then washed with 50 µL water for 5 minutes to maximise recovery of peptides. Digests were acidified with trifluoroacetic acid (TFA) to a final concentration of 0.5% and centrifuged for 10 minutes at 17,000 g to remove insoluble material. The digested peptides were desalted using Strata C_18_-E columns (55 µm, 70 Å, 50 mg, 1 mL tubes – Phenomenex), elution volume was 3x 90 µL acetonitrile, peptides were dried down using a miVac Centrifugal Concentrator (Barnstead).

### Mass spectrometry data acquisition and analysis of CRK9-proximal proteins affinity purification

After Samples were loaded onto a nanoAcquity UPLC system (Waters) equipped with a PharmaFluidics µPAC C_18_, Trapping column and a PharmaFluidics 50 cm µPAC C_18_ nano-LC column (5 µm pillar diameter, 2.5 µM inter-pillar distance). The trap wash solvent was 0.1% (v/v) aqueous formic acid and the trapping flow rate was 10 µL min^-1^. The trap was washed for 5 minutes before switching flow to the capillary column. Separation used a gradient elution of two solvents (solvent A: aqueous 0.1% (v/v) formic acid; solvent B: acetonitrile containing 0.1% (v/v) formic acid). The analytical flow rate was 1 µL min^-1^ and the column temperature was 50°C. The gradient profile was linear 2.5-30% B over 30 minutes then linear 30-90% B over 5 minutes. All runs then proceeded to wash with 90% solvent B for 5 minutes. The column was returned to initial conditions and re-equilibrated for 5 minutes before subsequent injections.

The nanoLC system was interfaced with a maXis HD LC-MS/MS system (Bruker Daltonics) with CaptiveSpray ionisation source (Bruker Daltonics). Positive ESI-MS and MS/MS spectra were acquired using MRM mode to define data independent acquisition (DIA) windows with a width of 5 Th between *m/z* 450-650. Instrument control, data acquisition and processing were performed using Compass 1.7 software (microTOF control, Hystar and DataAnalysis, Bruker Daltonics). Instrument settings were: ion spray voltage: 1,450 V, dry gas: 3 L min^-1^, dry gas temperature 150°C, ion acquisition range: *m/z* 280-1,600, spectra rate: 15 Hz, quadrupole low mass: 322 *m/z*, collision RF: 1,400 Vpp, transfer time 120 ms. The collision energy was set to 22 for DIA windows below m/z 600 and to 24 for higher m/z windows.

LC-MS data, in Bruker.d format, were converted to .mzML format using MSConvert (ProteoWizard) before analysing using DIA-NN (1.8.1) with searching against and in-silico predicted spectral library, derived from the LmexCas9T7-prot database appended with common proteomic contaminants. Search criteria were set to maintain a false discovery rate (FDR) of 1%. Peptide-centric output in .tsv format, was pivoted to protein-centric summaries using KNIME and data filtered to require protein q-values < 0.01 and a minimum of two peptides per accepted protein. Protein intensities were log2 transformed and proximal proteins were determined with the limma package [46] using options trend = TRUE and robust = TRUE for the eBayes function. Protein intensities in CRK9 samples were compared to those in the spatial reference BDF7 to determine proximal proteins. Multiple testing correction was carried out according to Benjamini & Hochberg, the false discovery rate for identified proximals was 1%.

### SM1-71 affinity purification

Cell pellets (in triplicate for each sample) of 3×10^8^ parasites washed with PBS were used for each affinity purification and lysed in 300 μL ice-cold lysis buffer (0.1% sodium dodecyl sulphate (SDS), 0.5% sodium deoxycholate, 1% IGEPAL CA-630, 0.1 mM EDTA, 125 mM NaCl, 50 mM Tris-HCl pH 7.5) containing 0.1 mM 4-(2-aminoethyl)benzenesulfonyl fluoride, 1 μg mL^-1^ pepstatin A, 1 μM E-64 and 0.4 mM 1-10 phenanthroline. In addition, every 10 mL of lysis buffer was supplemented with 100 μL protease inhibitor cocktail (abcam) plus 1 tablet of cOmplete protease inhibitor cocktail and PhosSTOP (Roche). Cells were lysed using a Bioruptor sonicator (Diagenode) for 3 cycles (30s on/off) in 1.5 mL microtubes containing silica beads to aid shearing. 1 μL of BaseMuncher endonuclease (250 Units, abcam) was added to each lysate and incubated on ice for 1 hour to digest nucleic acids. Lysates were clarified by centrifugation at 10,000 g for 10 minutes at 4°C and the supernatant transferred to LoBind Eppendorf microtubes. Lysates were pre-treated with SM1-71 (final concentration 4 uM) or DMSO (to 0.2%) for 1 hour before adding SM1-71-biotin for 2 hours. For enrichment of biotinylated material, 100 μL of magnetic streptavidin bead suspension (1 mg of beads, Resyn Biosciences, washed in PBS) was used to affinity purify by end-over-end rotation at 4°C overnight. Beads with bound proteins were washed in 500 μL of the following for 5 minutes each: lysis buffer (plus protease inhibitors); lysis buffer; PBS + 0.025% Tween-20 then PBS/Tween + 2% SDS (55°C) before rinsing with PBS/Tween and transferring the beads to clean LoBind tubes. Further washes using 4 M urea, 6 M urea, 1 M KCl and 50 mM triethylammonium bicarbonate (TEAB) pH 8.5 were performed and the beads stored at −20°C.

### Mass spectrometry data acquisition and analysis of SM1-71 affinity purification

After rinsing with 50 mM TEAB, proteins were denatured, reduced and enzymatically digested on-bead with the addition of 100 µL of 50 mM TEAB containing the following: 0.01% (w/w) ProteaseMAX surfactant (Promega); 2.9 mg mL^-1^ tris(2-carboxyethyl)phosphine; 100 mM CaCl_2_ and 2.5 µg sequencing grade trypsin/Lys-C mix (Promega). Beads were shaken for 5 minutes at 850 rpm before incubation overnight at 37°C. Supernatant containing peptide was acidified to contain 0.5% trifluoracetic acid before spinning at 17,000 g for 10 minutes to pellet precipitated ProteaseMAX. A 40 µL aliquot of soluble material was taken per sample for LC-MS acquisition. Resulting peptides were desalted with C_18_ ZipTip (0.2 uL, Millipore) before being re-suspended in aqueous 0.1% (v/v) trifluoroacetic acid.

Peptides were loaded onto an mClass nanoflow UPLC system (Waters) equipped with a nanoEaze M/Z Symmetry 100 Å, C_18_, 5 µm trap column (180 µm x 20 mm, Waters) and a PepMap, 2 µm, 100 Å, C_18_ EasyNano nanocapillary column (75 µm x 500 mm, Thermo). The trap wash solvent was aqueous 0.05% (v/v) trifluoroacetic acid and the trapping flow rate was 15 µL min^-1^. The trap was washed for 5 minutes before switching flow to the capillary column. Separation used gradient elution of two solvents: solvent A, aqueous 0.1% (v/v) formic acid; solvent B, acetonitrile containing 0.1% (v/v) formic acid. The flow rate for the capillary column was 300 nL min^-1^ and the column temperature was 40°C. The linear multi-step gradient profile was: 3-10% B over 7 minutes, 10-35% B over 30 minutes, 35-99% B over 5 minutes and then proceeded to wash with 99% solvent B for 4 minutes. The column was returned to initial conditions and re-equilibrated for 15 minutes before subsequent injections.

The nanoLC system was interfaced with an Orbitrap Fusion Tribrid mass spectrometer (Thermo) with an EasyNano ionisation source (Thermo). Positive ESI-MS and MS^2^ spectra were acquired using Xcalibur software (version 4.0, Thermo). Instrument source settings were: ion spray voltage, 1,900 V; sweep gas, 0 Arb; ion transfer tube temperature; 275°C. MS^1^ spectra were acquired in the Orbitrap with: 120,000 resolution, scan range: *m/z* 375-1,500; AGC target, 4e^5^; max fill time, 100 ms. Data dependant acquisition was performed in top speed mode using a 1 s cycle, selecting the most intense precursors with charge states >1. Easy-IC was used for internal calibration. Dynamic exclusion was performed for 50 s post precursor selection and a minimum threshold for fragmentation was set at 5e^3^. MS^2^ spectra were acquired in the linear ion trap with: scan rate, turbo; quadrupole isolation, 1.6 *m/z*; activation type, HCD; activation energy: 32%; AGC target, 5e^3^; first mass, 110 *m/z*; max fill time, 100 ms. Acquisitions were arranged by Xcalibur to inject ions for all available parallelizable time.

LC-MS chromatograms in .raw format were imported into Progenesis QI (Waters, v.3.3) for peak picking and alignment. A concatenated MS^2^ peak list was exported and searched against the *L. mexicana* subset of the TryTripDB database (v.54) appended with common proteomic contaminants using Mascot Server (Matrix Science v.2.5). Database search criteria specified: Enzyme, trypsin; Max missed cleavages, 1; Variable modifications, Oxidation (M); Peptide tolerance, 3 ppm; MS/MS tolerance, 0.5 Da. Peptide identifications were filtered through the Percolator algorithm to achieve a false discovery rate of 1% as assessed empirically against a reversed database search. Peptide identifications were reimported into Progenesis QI and associated with precursor intensity signals and identifications mapped between runs. Relative peptide abundance was obtained by integrating areas under identified MS^1^ signals and inference from peptide values. Normalisation was applied between runs on the basis of total observed peptide signal. Protein quantifications were filtered to require a minimum of two peptides per protein. A multi-way ANOVA was applied within Progenesis QI to call differing abundance with the Hochberg and Benjamini multiple test correction applied and q<0.05 set as significance level.

### Statistics

Data were collected from at least three independent experiments, unless otherwise indicated. Statistical analyses were performed using GraphPad Prism v10.2.0. The appropriate tests were conducted and are detailed in the corresponding figure legends. Results are presented as the mean ± standard error of mean (SEM).

## Acknowledgements

This work was supported by funding from MRC GCRF (MR/P027989/1) and Wellcome Trust (223045/Z/21/Z). The funders had no role in study design, data collection, data analysis, interpretation, and writing of the manuscript. We thank our colleagues in the Bioscience Technology Facility of University of York who provided insight and expertise that greatly assisted our microscopy, flow cytometry, genomic data analysis, and mass spectrometry research. The York Centre of Excellence in Mass Spectrometry was created thanks to a major capital investment through Science City York, supported by Yorkshire Forward with funds from the Northern Way Initiative, and subsequent support from EPSRC (EP/K039660/1; EP/M028127/1).

## Author contributions

Conceived the project: J.B.T.C., J.C.M., A.J.W., and R.M.C. Supervised the project: J.C.M. and A.J.W. Conceived and designed the experiments: J.B.T.C., J.A.B., R.M.C., A.P.C.A.L., N.G.J., and P.S. Performed the experiments: J.B.T.C., J.A.B., N.G.J., and P.Z.R. Analysed data: J.B.T.C., J.A.B., A.P.C.A.L., N.G.J., A.J.W., and J.C.M. Acquired funding: J.C.M., A.P.C.A.L., and A.J.W. Wrote the manuscript: J.B.T.C and J.C.M. All authors read and approved the final manuscript.

## Supporting information captions

**Supplementary Information:** Supplementary Tables and Figures supporting the main text.

**Supplementary Data 1:** Mapping of gatekeeper residues across the *L. mexicana* kinome.

**Supplementary Data 2:** Identification of protein kinases with targetable cysteines by the multi-targeted SM1-71-biotin probe.

**Supplementary Data 3:** Identification of CRK9-proximal proteome.

