## Supplementary Information for "Chemical genetics reveals *Leishmania* KKT2 and CRK9 kinase activity is required for cell cycle progression"

**Supplementary Table 1. Sequence of oligonucleotides used to generate sgRNA and DNA repair template for CRISPR-Cas9 edited lines.**

| Oligo ID | Engineered cell line | Sequence (5' → 3') | Description |
|---|---|---|---|
| OL6137 (R) | N/A | AAAAGCACCGACTCGGTGCCACTTTTTCAAGTTGATAACGGACTAGCCTTATTTTAACTTGCTATTTCTAGCTCTAAAAC | Reverse oligo to generate 5'sgRNA and 3'sgRNA |
| OL11240 | AS CLK1$^{M213}$ / AS CLK2$^{M220}$ | GAAATTAATACGACTCACTATAGGCGCTATTTCCAGAACGACAGGTTTTAGAGCTAGAAATAGC | 5'sgRNA |
| OL11241 | | GAAATTAATACGACTCACTATAGGCAAGTACGGCCCCTGCCTGCGTTTTAGAGCTAGAAATAGC | 3'sgRNA |
| OL11613 (F) | AS KKT2$^{M146}$ | GAAATTAATACGACTCACTATAGGCAAATTCTACGGTGCTGGTAGTTTTAGAGCTAGAAATAGC | 5'sgRNA |
| OL11614 (F) | | GAAATTAATACGACTCACTATAGGGTAGTAATGGAGCGGTGCGCGTTTTAGAGCTAGAAATAGC | 3'sgRNA |
| OL11607 (F) | AS KKT3$^{M110}$ | GAAATTAATACGACTCACTATAGGACAGCGGACTTGATCGTTATGTTTTAGAGCTAGAAATAGC | 5'sgRNA |
| OL11608 (F) | | GAAATTAATACGACTCACTATAGGAGAAGGTCGTGGAGCGTGCTGTTTTAGAGCTAGAAATAGC | 3'sgRNA |
| OL11601 (F) | AS CRK9$^{M501}$ | GAAATTAATACGACTCACTATAGGGAAGGACGTCTTCCTTGTAAGTTTTAGAGCTAGAAATAGC | 5'sgRNA |
| OL11602 (F) | | GAAATTAATACGACTCACTATAGGACTACTGTCCCTACGACCTGGTTTTAGAGCTAGAAATAGC | 3'sgRNA |
| OL14683 (F) | Δ*kkt3* | GAAATTAATACGACTCACTATAGGCTATGTGTGGTGGGCGATTTGTTTTAGAGCTAGAAATAGC | 5'sgRNA |
| OL14685 (F) | | GAAATTAATACGACTCACTATAGGTTCCCGTCGTTTTCGCTCACGTTTTAGAGCTAGAAATAGC | 3'sgRNA |
| OL14681 (F) | | AGATTTTGTTGTCTATCACAATCAAAGTAGGTGTATCGGATGTCAGTTGCGTGGCTGATGTCCGTATGTATAATGCAGACCT GCTGC | Upstream forward to generate repair template |
| OL14684 (R) | | CTTTTTTTTTCCGCATCCACGACCTCTGCACCAATTTGAGAGACCTGTGC | Downstream reverse to generate repair template |
| OL6945 (F) | KKT2::mNG::3xMyc | GAAATTAATACGACTCACTATAGGGAGGACAGCGCGTCGTGAGTGTTTTAGAGCTAGAAATAGC | 3'sgRNA |
| OL6943 (F) | KKT2_AS::mNG::3xMyc | GAGACGGTCCTCAACAACAATTTCCGGAGAGGTTCTGGTAGTGGTTCCGG | Downstream forward to generate repair template |
| OL6944 (R) | KKT2_AS::3xMyc::mT | AGAAGATACACGTAAACACGAACTCAACCACCAATTTGAGAGACCTGTGC | Downstream reverse to generate repair template |

R, reverse oligo; F, forward oligo; N/A, not applicable; AS, analog-sensitive kinase; *Δ*, knockout target gene.

The R sgRNA oligo OL6137 holds the Cas9 scaffold (black) and the complementary sequence to the forward primer (blue). The F oligo sequences to generate sgRNA are coloured as follows: T7 promoter in black; sgRNA target sites in red; complementary sequence to sgRNA scaffold in blue. To generate linear DNA fragment for *in vivo* transcription of the single guide RNA, Q5® High-Fidelity DNA Polymerase (New England BioLabs®inc. Cat. M0491L) was used in a 40 µL reaction mix containing 1x Q5 Reaction buffer, 200 µM dNTPs, 2 µM of the sgRNA scaffold oligo (OL6137), 2 µM of the forward oligo and 0.8 units of Q5® High-Fidelity DNA Polymerase. Cycling conditions: initial denaturation at 98ºC for 30 seconds, followed by 40 cycles of 98ºC for 10 seconds (denaturation), 60ºC for 30 seconds (annealing), 72ºC for 15 seconds (extension) and a final extension step at 72ºC for 10 minutes.

The upstream and downstream oligo sequences to generate DNA repair template are coloured as follows: 30 bp for homology direct recombination in red; barcode in green; complementary sequence to the template plasmid in blue. DNA repair template was generated using Q5® High-Fidelity DNA Polymerase in a 40 µL reaction mix containing 1x Q5 Reaction buffer, 200 µM dNTPs, 2 µM of each forward and reverse oligos, 0.6 ng of the template and 0.8 units of Q5® High-Fidelity DNA Polymerase. Cycling conditions: initial denaturation at 94ºC for 5 minutes followed by 45 cycles of 94ºC for 30 seconds (denaturation), 65ºC for 30 seconds (annealing), 72ºC for 2.25 minutes (extension) and a final extension step at 72ºC for 10 minutes. Templates used to generate DNA repair template to knockout and endogenously tag genes (sequences available on http://leishgedit.net/): pPLOTv1 blast-mNeonGreen-blast; pPLOTv1 puro-mNeonGreen-puro; pTBlast_v1; pTPuro_v1.

**Supplementary Table 2. Sequence of the DNA repair templates used to engineer analog-sensitive kinases in *Leishmania*.**

| **Engineered cell line** | **Sequence** (ssDRT) |
|---|---|
| AS CLK1$^{M213G}$ / AS CLK2$^{M220G}$ | GACCGCTTCCCGCTGATGAAGATCCAGCGTTACTTTCAAAATGATTCTGGTCATATGTGCATCGTC**GGC**CCCAAATATGGACCGTGTCTCCTAGACTGGATCATGAAGCACGGCCCCTTC |
| AS CLK1$^{M213A}$ / AS CLK2$^{M220A}$ | GACCGCTTCCCGCTGATGAAGATCCAGCGTTACTTTCAAAATGATTCTGGTCATATGTGCATCGTC**GCG**CCCAAATATGGACCGTGTCTCCTAGACTGGATCATGAAGCACGGCCCCTTC |
| AS KKT2$^{M146G}$ | GAGCCATTTCTCGCGCCATCCCAACATTGTCAAGTTTTATGGAGCGGGCCGCGACGAGGACCGAGCGTATGTGGTG**GGC**GAACGTTGTGCAGGCAAGTCGCTTCACGACGTCATAGCCAG |
| AS KKT2$^{M146A}$ | GAGCCATTTCTCGCGCCATCCCAACATTGTCAAGTTTTATGGAGCGGGCCGCGACGAGGACCGAGCGTATGTGGTG**GCG**GAACGTTGTGCAGGCAAGTCGCTTCACGACGTCATAGCCAG |
| AS KKT3$^{M110G}$ | CGCGTTTCGAGTTTGGAGCGCTCAACAAGACGGCAGATCTCATTGTGATCGGA**GGC**GAACTATGCGTCCCCAGTACTCTGCATGATTTGCTCCTCAGCACTCGTATCACCAGCGAAGCGG |
| AS KKT3$^{M110A}$ | CGCGTTTCGAGTTTGGAGCGCTCAACAAGACGGCAGATCTCATTGTGATCGGA**GCG**GAACTATGCGTCCCCAGTACTCTGCATGATTTGCTCCTCAGCACTCGTATCACCAGCGAAGCGG |
| AS CRK9$^{M501G}$ | CACACGACCGCTGGCCGCCGTCGGCGCTGCAAGCAAGGCGAAAGATGTTTTTCTGGTG**GGC**GATTATTGCCCATATGATCTTGGGAGCTACATGCGGCGGTACGCGACTGTGGCAGAGCT |
| AS CRK9$^{M501A}$ | CACACGACCGCTGGCCGCCGTCGGCGCTGCAAGCAAGGCGAAAGATGTTTTTCTGGTG**GCG**GATTATTGCCCATATGATCTTGGGAGCTACATGCGGCGGTACGCGACTGTGGCAGAGCT |

AS, analog-sensitive kinase; ssDRT, single-stranded DNA repair template.

Repair template sequence is coloured as follows: homology arm in black, recoded codons in blue, and mutated target codon in red.

**Supplementary Table 3. Sequence of oligonucleotides used to screen the CRISPR-Cas9 engineered *L. mexicana* cell lines.**

| Oligo ID | Engineered cell line | Sequence | Description |
|---|---|---|---|
| OL11917 (F) | CLK1 | ATGTCGCGCAGCCAGAGC | Specific amplification of CLK1 to then screen the analog sensitive by additional PCR or restriction site digestion – Sanger sequencing. |
| OL8401 (R) | | GCAGCTCGCTGTGAAAGTAG | |
| OL11802 (F) | CLK2 | AGGAAACCGCTGTGTCAAGCACC | Specific amplification of CLK1 to then screen the analog sensitive or resistant by additional PCR or restriction site digestion – Sanger sequencing. |
| OL8401 (R) | | GCAGCTCGCTGTGAAAGTAG | |
| OL11592 (F) | AS CLK1$^{M213G}$ / AS CLK2$^{M220G}$ | CCAGAACGACAGCGGCCA | Semi-nested PCR to specifically amplify the wildtype CLK1 or CLK2 genome. |
| OL8401 (R) | | GCAGCTCGCTGTGAAAGTAG | |
| OL12375 (F) | AS CLK1$^{M213G}$ / AS CLK2$^{M220G}$ | TCAAAATGATTCTGGTCATATGTGCATCGTCG | Semi-nested PCR to specifically amplify the AS CLK1 or AS CLK2 genome. |
| OL8401 (R) | | GCAGCTCGCTGTGAAAGTAG | |
| OL11617 (F) | AS KKT2$^{M146}$ | CTTCGCGTTAACGTGGATTT | PCR followed by restriction site digestion to screen analog sensitive mutant – Sanger sequencing |
| OL11618 (R) | | TGCAACCTCTGAGACCAGTG | PCR followed by restriction site digestion to screen analog sensitive mutant – Sanger sequencing |
| OL12165 (R) | | CCTCGAGATACGGATTCATGCG | Sanger sequencing |
| OL11611 (F) | AS KKT3$^{M110}$ | CATCGTGCTCTGCACTCTGT | PCR followed by restriction site digestion to screen analog sensitive mutant – Sanger sequencing |
| OL11612 (R) | | TAAGCGACTGTTCAGCAACG | PCR followed by restriction site digestion to screen analog sensitive mutant – Sanger sequencing |
| OL11605 (F) | AS CRK9$^{M501}$ | CTGCCGGGATGTGAACTACT | PCR followed by restriction site digestion to screen analog sensitive mutant – Sanger sequencing |
| OL11606 (R) | | GCGGTTCCTCCTTGAAGAAT | PCR followed by restriction site digestion to screen analog sensitive mutant – Sanger sequencing |
| OL14686 (F) | Δ*kkt3* | TCGTGCTCTGCACTCTGTTT | CRISPR-Cas9 Knocked out cell line screening |
| OL14687 (R) | | CCAGCGGGAGGGAGGATAAA | CRISPR-Cas9 Knocked out cell line screening |
| OL14688 (F) | | GTCACAGTGCAGGTGTACGA | CRISPR-Cas9 Knocked out cell line screening |
| OL14689 (R) | | CACGCATGCTCGATCAACAG | CRISPR-Cas9 Knocked out cell line screening |
| OL12735 (F) | KKT2::mNG::3xMyc | TGCAACGCTATGGTGGACAT | CRISPR-Cas9 endogenous tagged cell line screening |
| OL12736 (R) | KKT2_AS::mNG::3xMyc | TCATGGTCACGCATCTAGCG | CRISPR-Cas9 endogenous tagged cell line screening |
| OL14186 (R) | KKT2_AS::3xMyc::mT | CCTTGCGCTATTCGCTGTTG | CRISPR-Cas9 Knocked out cell line screening – Sanger sequencing |
| OL14188 (R) | | TTGAGGAAGTGCGGTACTGG | PCR followed by restriction site digestion to screen analog sensitive mutant – Sanger sequencing |
| OL9369 (R) | N/A | GCAGCAGGTCTGCATTATAC | Integration of the DNA repair template in the CRISPR-Cas9 knocked-out cell lines |
| OL12757 (F) | N/A | GCACAGGTCTCTCAAATTGG | Integration of the DNA repair template in the CRISPR-Cas9 C-terminus endogenous tagged cell lines |

R, reverse oligo; F, forward oligo; N/A, not applicable; AS, analog-sensitive kinase; *Δ*, knockout target gene.

Screening PCRs were performed using Platinum® Taq DNA Polymerase (Invitrogen™, Cat. 10966018) in a 25 µL reaction mix containing 1x PCR buffer, 200 µM dNTPs, 1.5 mM $MgCl_2$, 0.4 µM of each forward and reverse oligos, 6% (v/v) Kb extender and 0.02 U/µL Platinum® Taq DNA Polymerase. Cycling condition used was: initial denaturation at 94ºC for 5 minutes followed by 40 cycles of 94ºC for 30 seconds (denaturation), 60ºC for 30 seconds (annealing), 72ºC for 1 minute per Kb (extension) and a final extension step at 72ºC for 10 minutes.

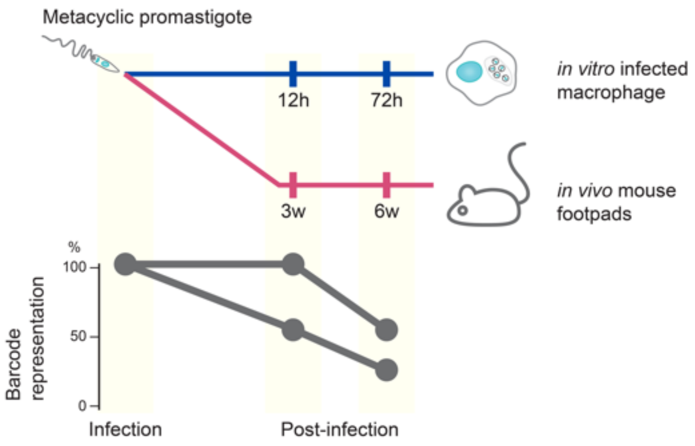

**Supplementary Fig 1. Schematic illustration to identify protein kinases required for *L. mexicana* amastigote survival.** The Bar-Seq dataset generated from the pooled barcoded library of protein kinase deleted mutants [1] was used to identify the protein kinases required for amastigote survival. The proportion of barcodes in the metacyclic promastigote stage and in two post-infection time points – 12 and 72 hours (h) for the *in vitro* infected macrophages, and 3 and 6 weeks (w) for the *in vivo* mouse footpads – was used to calculate fold changes in barcode representation relative to the preceding time point. The line graph shows the trajectories of the barcode representation in a mutant cell line used to classify a protein kinase as required for amastigote survival: (i) a significant ≥50% reduction in barcode abundance at the first post-infection time point, followed by an additional decrease of ≥30% at the second time point; or (ii) no significant change in the first time point, but a significant ≥50% reduction at the final time point. Statistical analysis, comparing the barcode proportion at each time point with those of the preceding time point, was performed by paired Student's t-test.

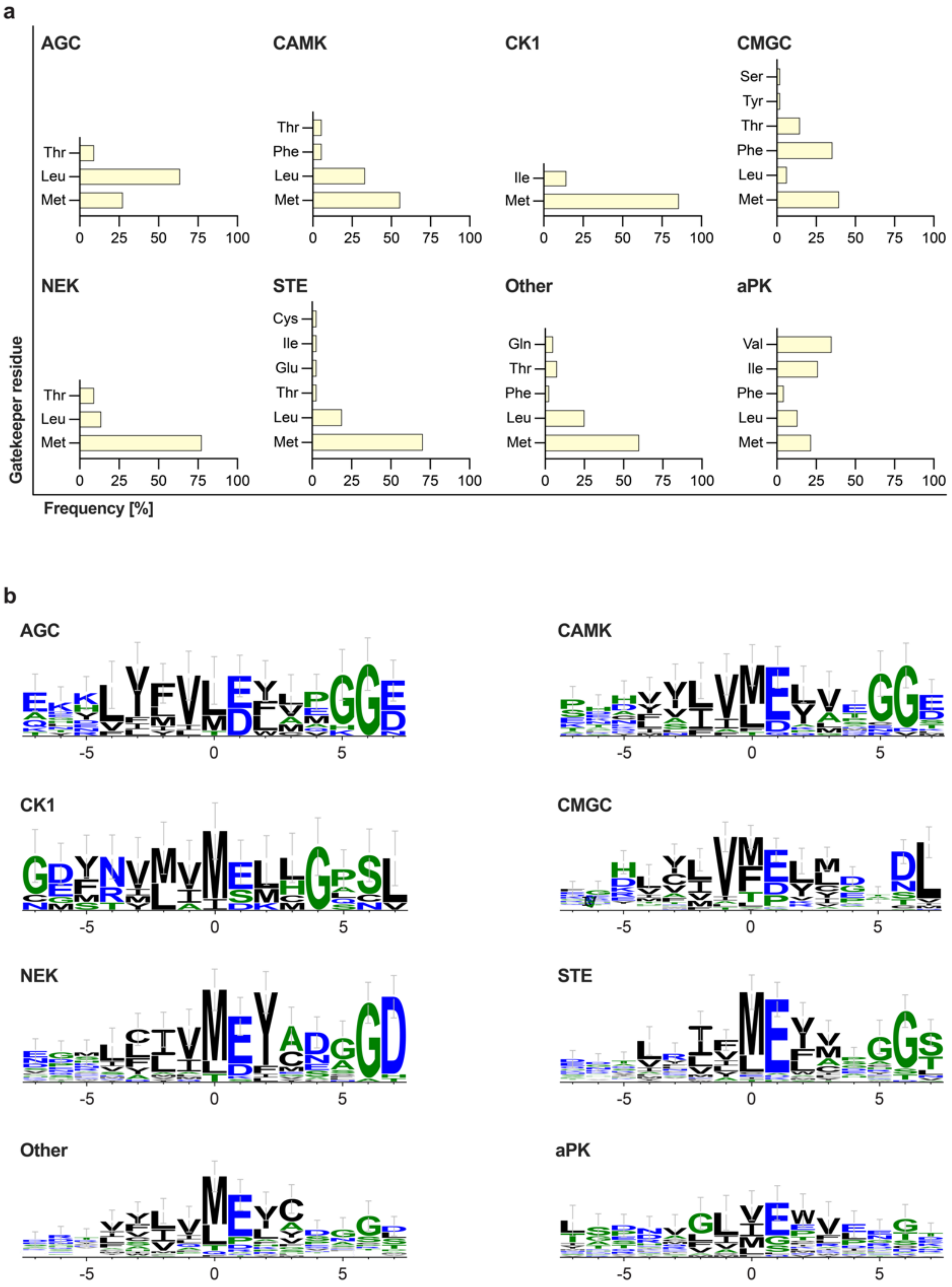

**Supplementary Fig 2. Gatekeeper residues in the *L. mexicana* kinome.** (a) Frequency distribution of gatekeeper residues across kinase groups/families. (b) Amino acid sequence logos depicting the region surrounding the gatekeeper (position 0) for each kinase group/family. The overall height of the stack indicates the level of sequence conservation at that position, while the height of individual symbols within the stack reflects the frequency of each amino acid at that position. Sequence logos were generated using WebLogo 3 [2].

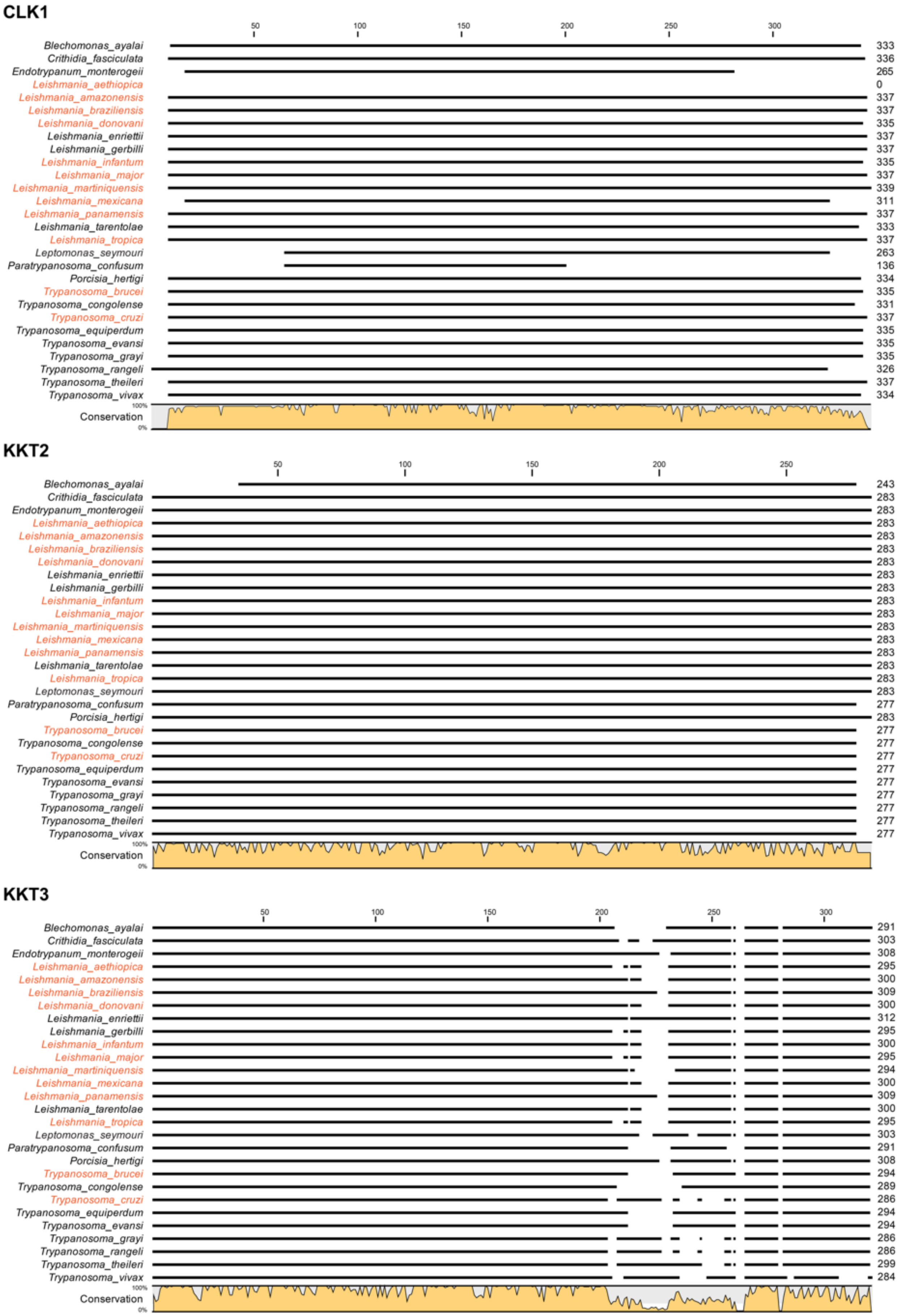

**Supplementary Fig 3. Conservation of CLK1, KKT2, KKT3 and CRK9 across trypanosomatids.**

**CRK9**

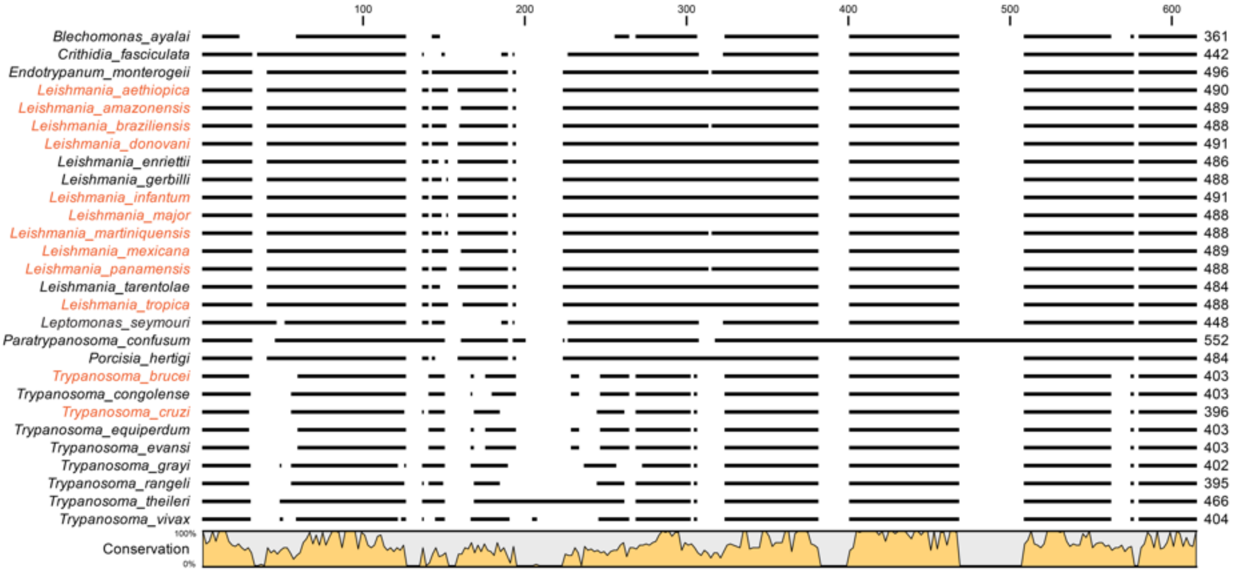

**Supplementary Fig 3. Conservation of CLK1, KKT2, KKT3 and CRK9 across trypanosomatids**. The protein kinase domain of *L. mexicana* CLK1 (LmxM.09.0400), KKT2 (LmxM.36.5350), KKT3 (LmxM.34.4050), and CRK9 (LmxM.27.1940) were aligned with their orthologues from reference trypanosomatid species available in TriTrypDB (https://tritrypdb.org/tritrypdb/app). For *L. donovani*, the CLK1 sequence from strain CL-SL was used due to incomplete sequencing of this gene in the reference strain BPK282A1. Similarly, for *T. cruzi*, KKT3 from the Dm28c 2014 strain was used, as the reference strain CL Brener Esmeraldo-like lacked an annotated KKT3 ortholog. Sequence aligment was performed using the Clustal Omega algorithm in CLC Genomics Workbench v22. The consensus line graph indicating sequence conservation is shown below the alignment. Species known to be human pathogens are highlighted in orange.

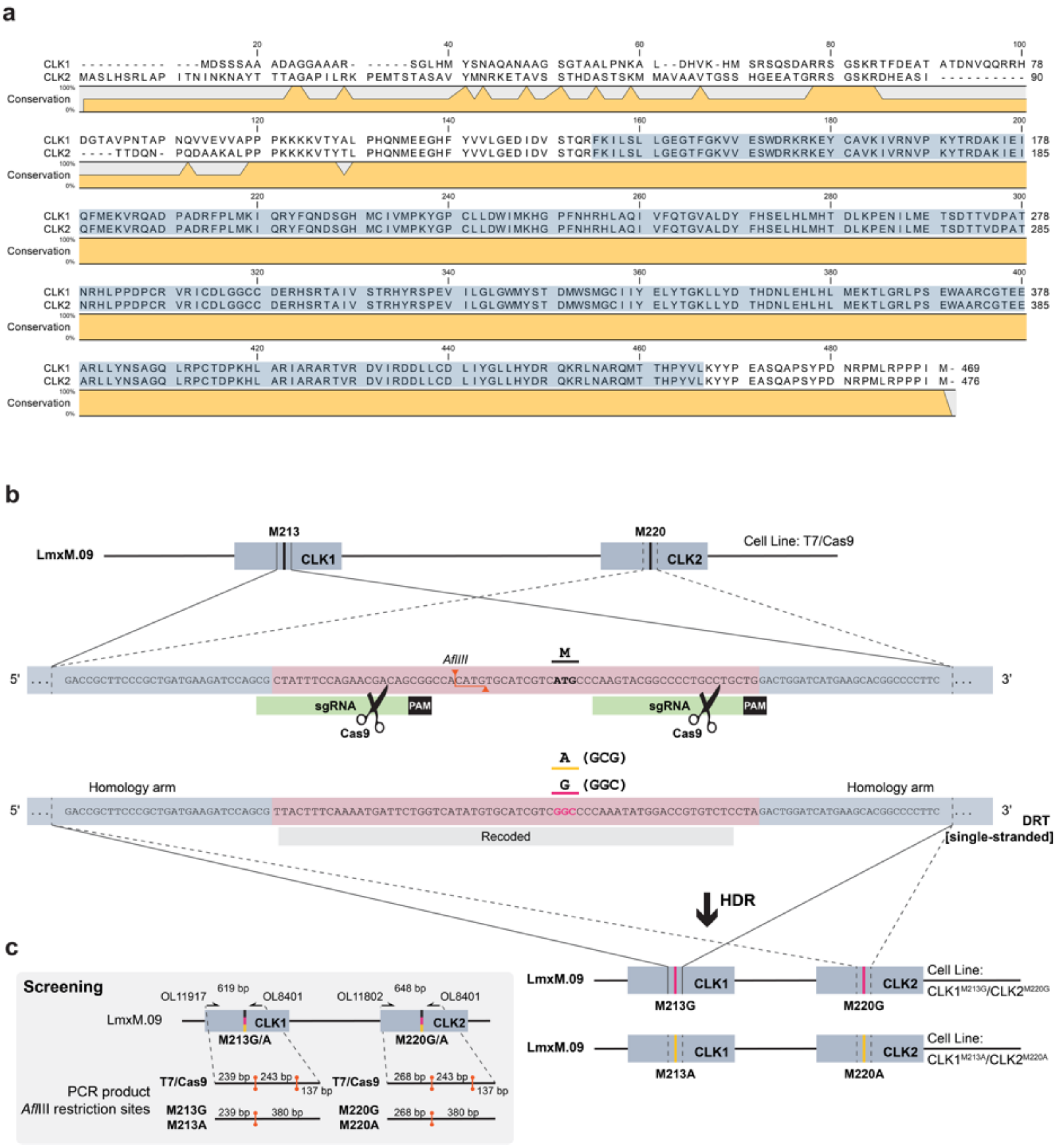

**Supplementary Fig 4. CRISPR-Cas9–mediated engineering of analog-sensitive CLK1/CLK2 in *Leishmania*.**

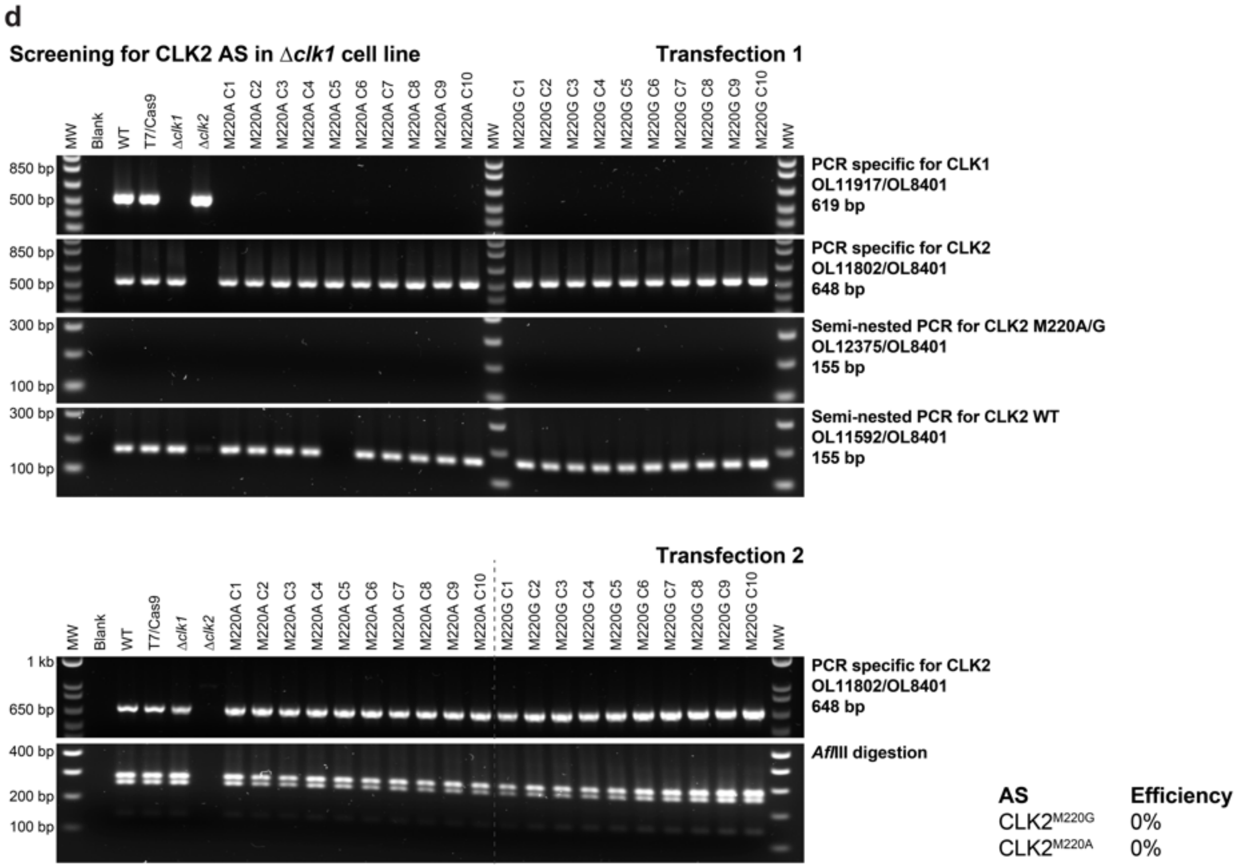

**Supplementary Fig 4. CRISPR-Cas9–mediated engineering of analog-sensitive CLK1/CLK2 in *Leishmania*.**

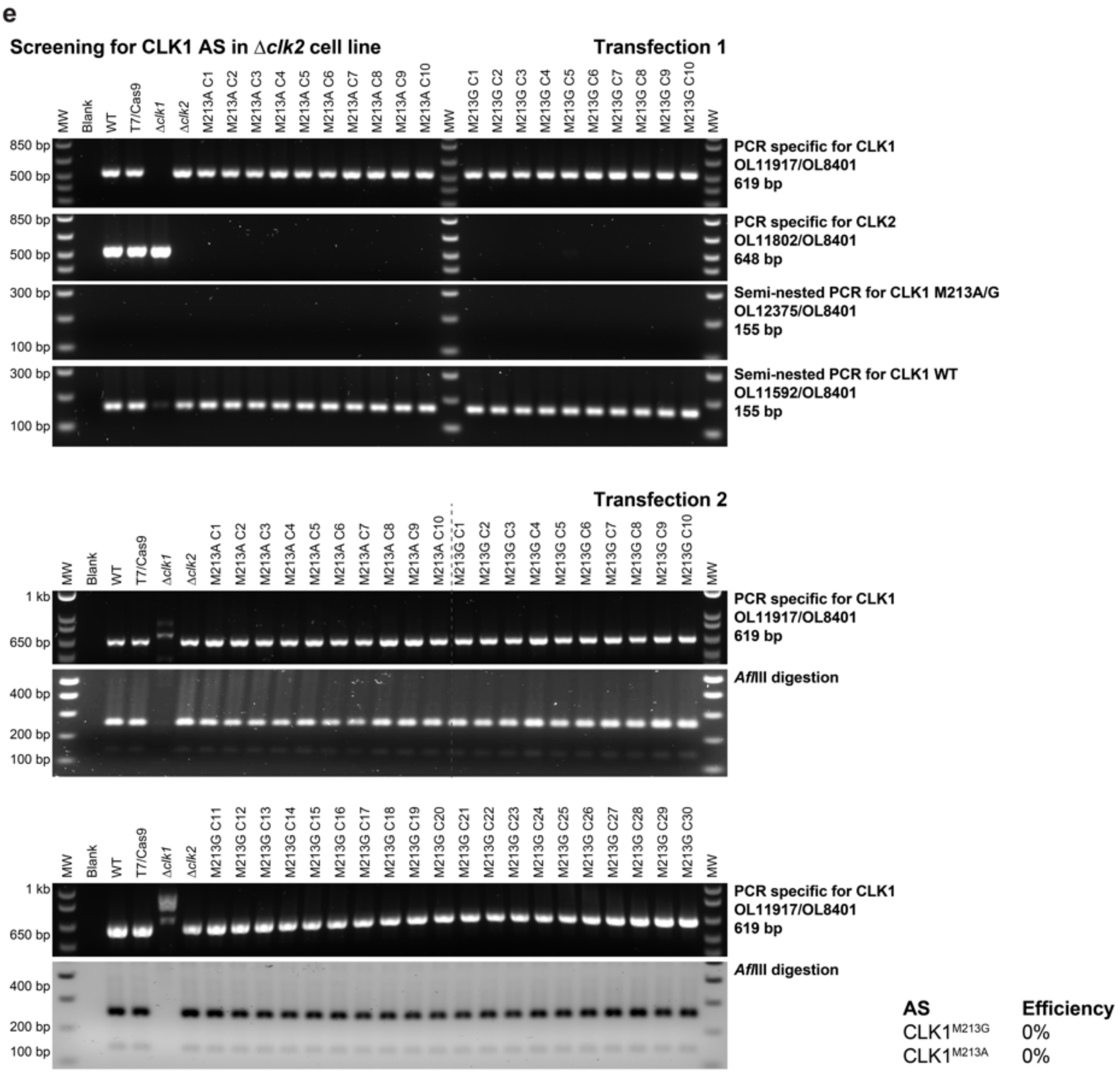

**Supplementary Fig 4. CRISPR-Cas9–mediated engineering of analog-sensitive CLK1/CLK2 in *Leishmania*.**

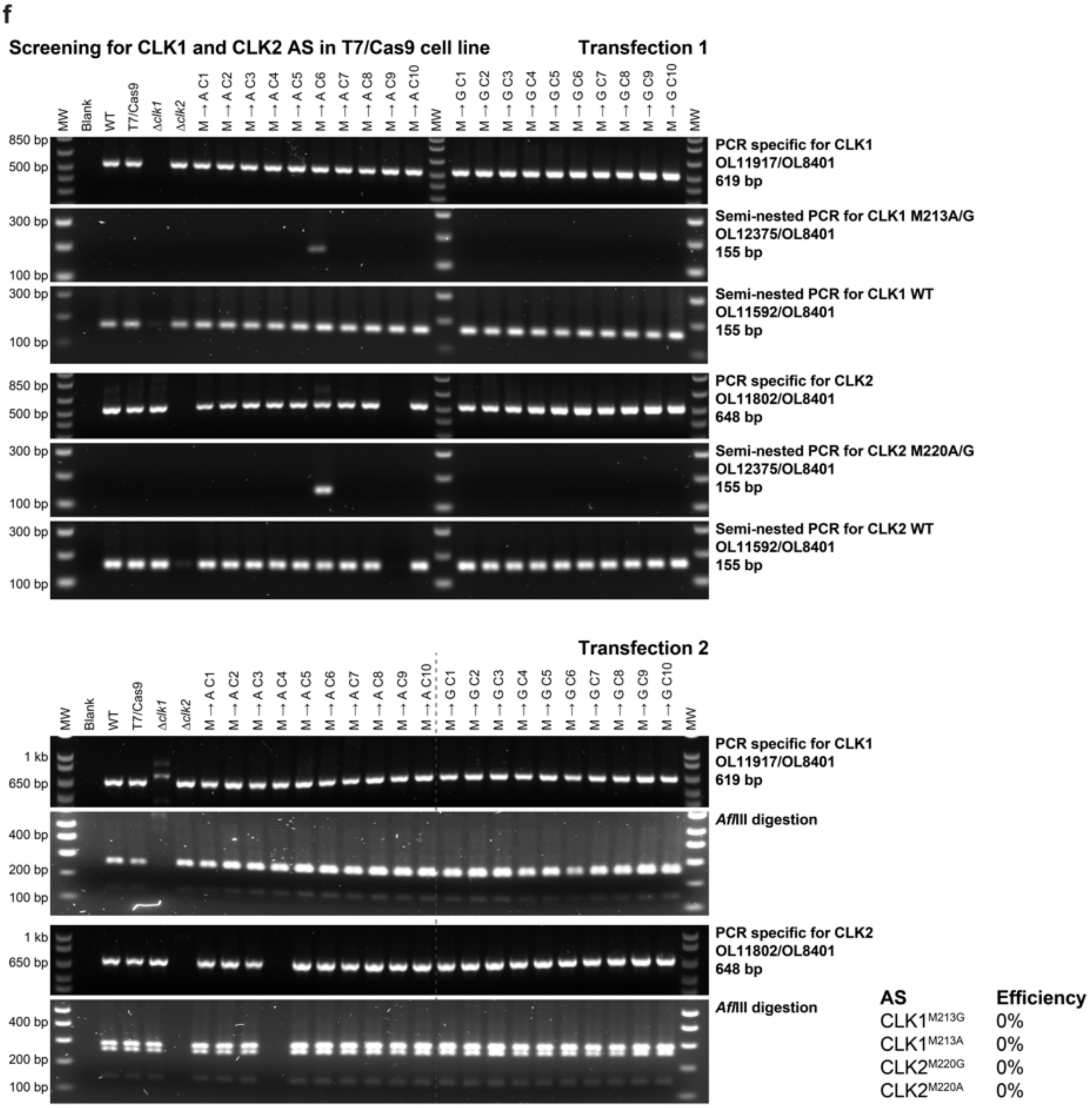

**Supplementary Fig 4. CRISPR-Cas9–mediated engineering of analog-sensitive CLK1/CLK2 in *Leishmania*.** (a) Sequence aligment was performed using the Clustal Omega algorithm in CLC Genomics Workbench v22. The consensus line graph indicating sequence conservation is shown below the alignment. The kinase domains were predicted using InterPro domain analysis [3], and the corresponding amino acid sequences are highlighted in blue. (b) Schematic of the CRISPR-Cas9 strategy used to engineer analog-sensitive kinases by substituting the CLK1 and CLK2 gatekeeper methionine (M) with glycine (G) or alanine (A). Linear DNA fragments for *in vivo* transcription of two single guide RNAs (sgRNAs), and a 120 bp DNA repair template (DRT) containing silent recoding mutations and the gatekeeper substitution were used. The mutations eliminated a *Afl*III restriction site, enabling genotypic screening of edited clones. PAM, protospacer adjacent motif; HDR, homology-directed repair. (c) Genotyping workflow (grey box) used to screen analog-sensitive clones. Genotyping was also performed using a semi-nested PCR: the first PCR was designed over the specific N-terminus for CLK1 (OL11917/OL8401) and CLK2 (OL11802/OL8404) followed by a second PCR that kept the reverse primer (OL8401) and using a specific primer to detect the wildtype genome (OL11592) or the recoded genome (OL12375). (d – e) Genotypic screening of individual clones (C1 – C30) for each gatekeeper mutation attempted in CLK1 and/or CLK2 genes within the *Δclk1*(c) *Δclk2* (d) or T7/Cas9 (e) cell lines. The editing efficiency for generating analog-sensitive mutants is indicated in the lower right corner.

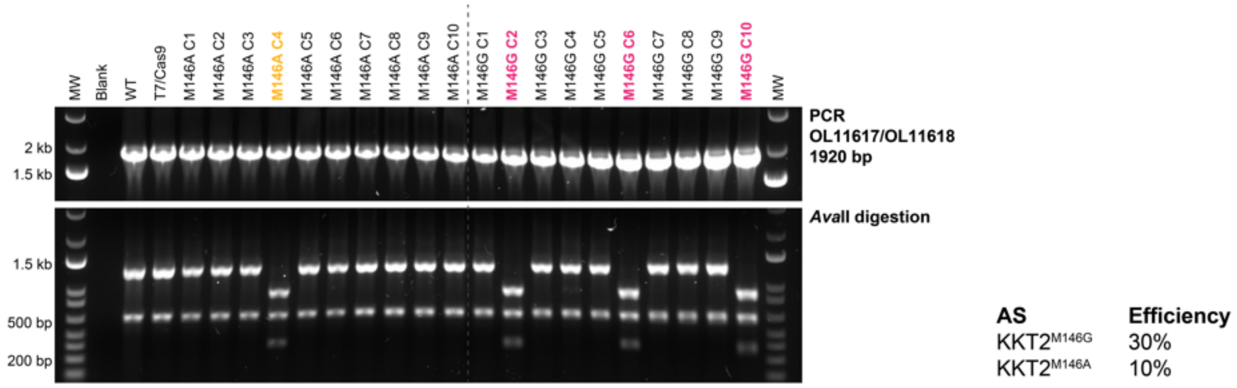

**Supplementary Fig 5. CRISPR-Cas9–mediated engineering of analog-sensitive KKT2 in *Leishmania*.** Genotypic screening of ten clones (C1 – C10) for each gatekeeper mutation introduced in KKT2. Genotyping results are color-coded as follows: black, wild-type; magenta, KKT2$^{M146G}$; yellow, KKT2$^{M146A}$. The editing efficiency for generating analog-sensitive mutants in this experiment is indicated in the lower right corner.

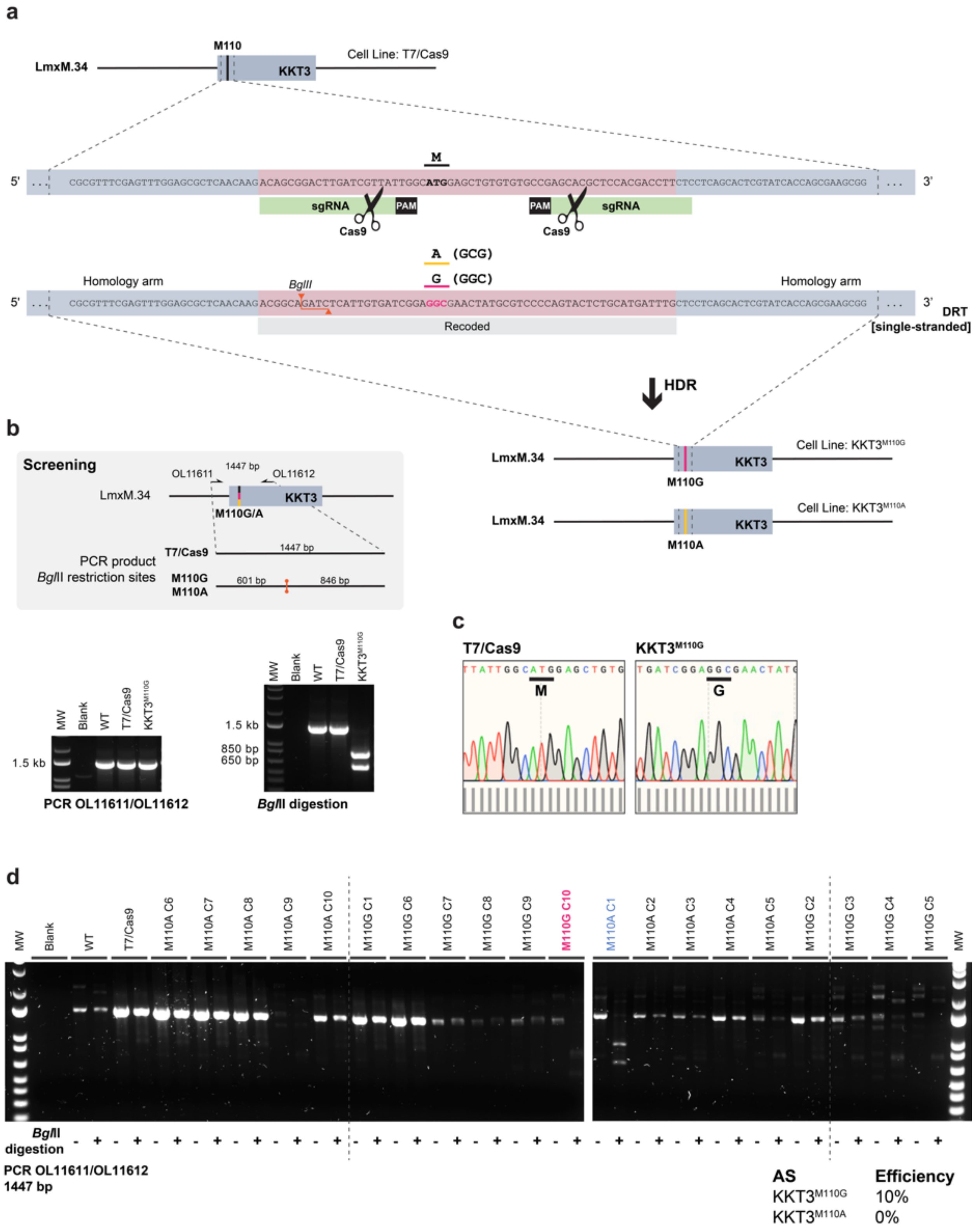

**Supplementary Fig 6. CRISPR-Cas9–mediated engineering of analog-sensitive KKT3 in *Leishmania*.** (a) Schematic of the CRISPR-Cas9 strategy used to engineer analog-sensitive kinases by substituting the KKT3 gatekeeper methionine (M) with glycine (G) or alanine (A). Linear DNA fragments for *in vivo* transcription of two single guide RNAs (sgRNAs), and a 120 bp single stranded DNA repair template (DRT) containing silent recoding mutations and the gatekeeper substitution were used. The mutations introduced a *Bgl*II restriction site, enabling genotypic screening of edited clones. PAM, protospacer adjacent motif; HDR, homology-directed repair. (b) Genotyping workflow (top grey box) and PCR-restriction digest results (bottom) for selected analog-sensitive clones. (c) Sanger sequencing of the engineered KKT3 locus confirms the substitution of the gatekeeper methionine with glycine in the KKT3$^{M110G}$ line. Sequencing chromatograms were visualized in SnapGene v7.2; bar graphs below indicate per-base quality scores. (d) Genotypic screening of ten clones (C1 – C10) for each gatekeeper mutation introduced in KKT3. Genotyping results are color-coded as follows: black, wild-type; magenta, $KKT3^{M110G}$; blue, $KKT3^{M110A}$. The editing efficiency for generating analog-sensitive mutants in this experiment is indicated in the lower right corner.

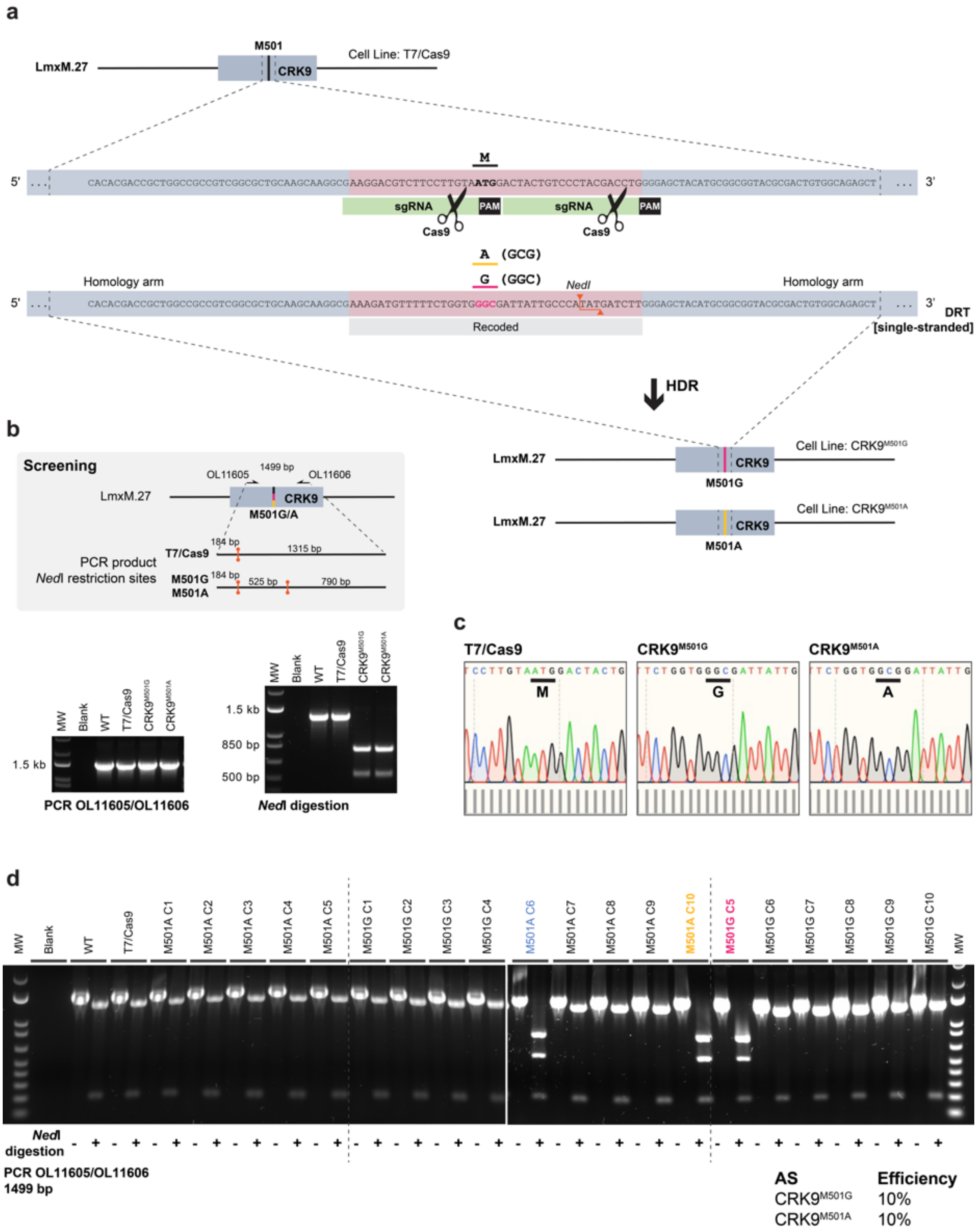

**Supplementary Fig 7. CRISPR-Cas9–mediated engineering of analog-sensitive CRK9 in *Leishmania*.** (a) Schematic of the CRISPR-Cas9 strategy used to engineer analog-sensitive kinases by substituting the CRK9 gatekeeper methionine (M) with glycine (G) or alanine (A). Linear DNA fragments for *in vivo* transcription of two single guide RNAs (sgRNAs), and a 120 bp single stranded DNA repair template (DRT) containing silent recoding mutations and the gatekeeper substitution were used. The mutations introduced a *Ned*I restriction site, enabling genotypic screening of edited clones. PAM, protospacer adjacent motif; HDR, homology-directed repair. (b) Genotyping workflow (top grey box) and PCR-restriction digest results (bottom) for selected analog-sensitive clones. (c) Sanger sequencing of the engineered CRK9 locus confirms the substitution of the gatekeeper methionine with glycine or alanine in the CRK9$^{M501G}$ and CRK9$^{M501A}$ lines, respectively. Sequencing chromatograms were visualized in SnapGene v7.2; bar graphs below indicate per-base quality scores. (d) Genotypic screening of ten clones (C1 – C10) for each gatekeeper mutation introduced in CRK9. Genotyping results are color-coded as follows: black, wild-type; magenta, $CRK9^{M501G}$; yellow, $CRK9^{M501A}$; blue, $CRK9^{M501A}$. The editing efficiency for generating analog-sensitive mutants in this experiment is indicated in the lower right corner.

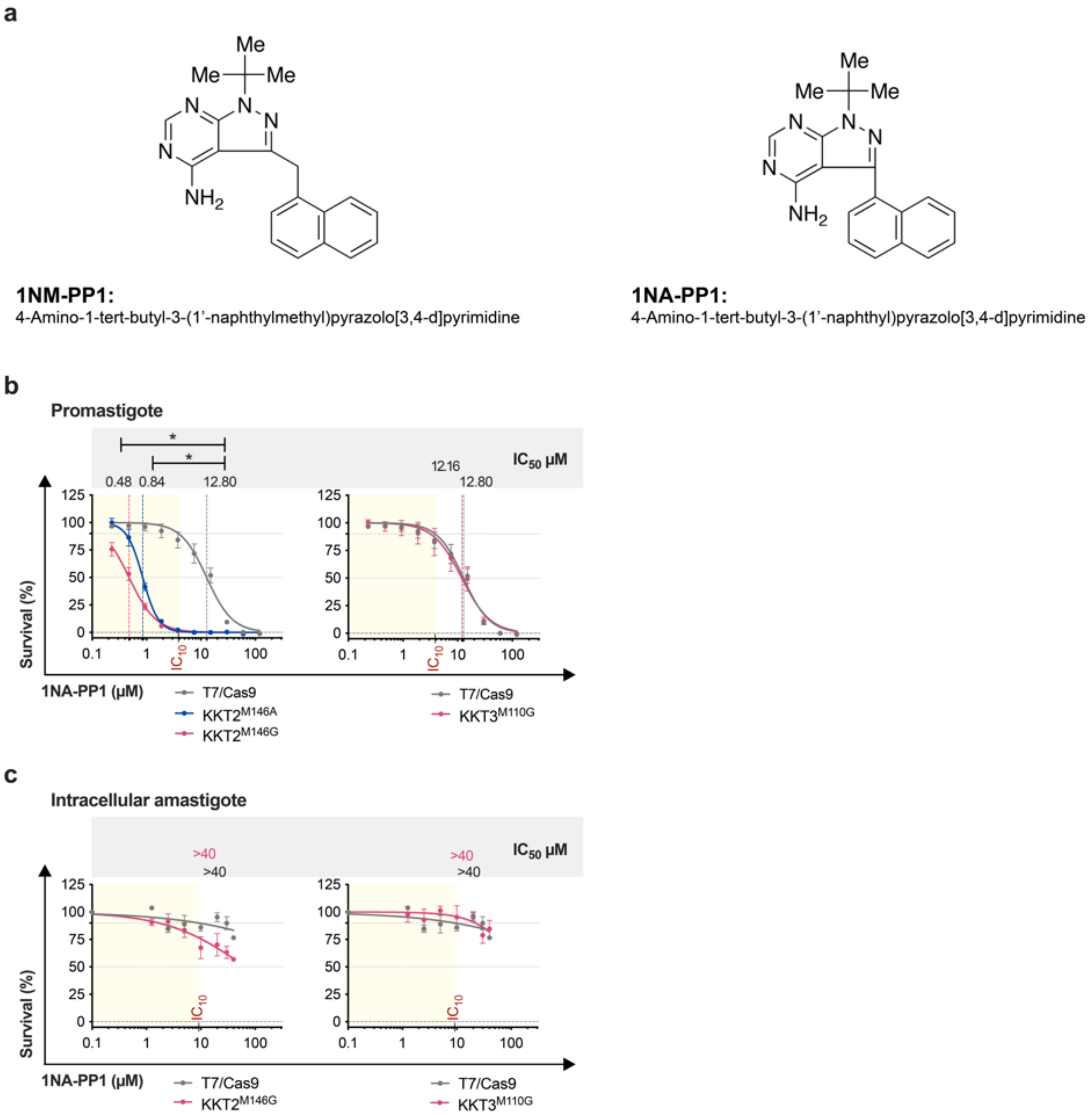

**Supplementary Fig 8. Susceptibility of *L. mexicana* lines expressing analog-sensitive variants of KKT2 and KKT3 to 1NA-PP1.** (a) Chemical structures of the bumped kinase inhibitors used in this study. Parasite viability under 1NA-PP1 treatment was assessed in *L. mexicana* analog-sensitive lines and wild-type T7/Cas9. (b-c) Dose-response curves were fitted using GraphPad Prism v10.4.1, with viability normalized to untreated controls (set at 100% for each cell line). Statistical significance was evaluated using unpaired two-tailed Student's t-tests (*p-value <0.05). The $IC_{10}$ value denotes the concentration of 1NA-PP1 that reduces the viability of the wild-type T7/Cas9 line by 10%. (b) Susceptibility of promastigotes to 1NA-PP1 was measured using a resazurin-based viability assay. Data represent mean ± SEM from three biological replicates. (c) Susceptibility of intracellular amastigotes to 1NA-PP1 was assessed by quantifying the percentage of infected macrophages. Data represent mean ± SEM from four biological replicates. Susceptibility of *L. mexicana* lines expressing AS variants of CRK9 was previously published by Jones N.G. et al., 2023 [4].

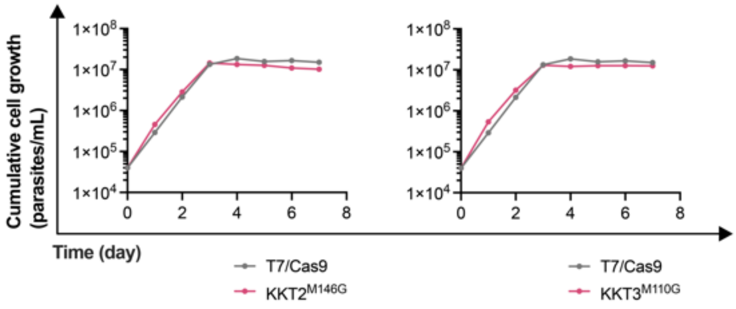

**Supplementary Fig 9. Growth curves of *L. mexicana* analog-sensitive kinase mutants and parental line.** Growth kinetics of *L. mexicana* promastigotes were assessed for the parental T7/Cas9 line and analog-sensitive kinase mutant lines (KKT2$^{M146G}$ and KKT3$^{M110G}$). Growth curve of *L. mexicana* line expressing AS variants of CRK9$^{M501G}$ was previously published by Jones N.G. et al., 2023 [4]. Cultures were initiated at a density of $4\times10^4$ cells mL$^{-1}$ in HOMEM medium supplemented with 10% heat-inactivated fetal bovine serum, and cumulative cell densities were measured daily by manual counting using a Neubauer chamber. Growth rates were calculated from the logarithmic phase of the growth curve (0 – 96 h) and are reported as mean ± SEM: T7/Cas9, 1.53 ± 0.007; KKT2$^{M146G}$, 1.45 ± 0.008; KKT3$^{M110G}$, 1.43 ± 0.022; and CRK9$^{M501G}$, 1.43 ± 0.006. Statistical analysis using the non-parametric Kruskal–Wallis test revealed no significant differences in growth rates between the AS mutants and the parental T7/Cas9 line.

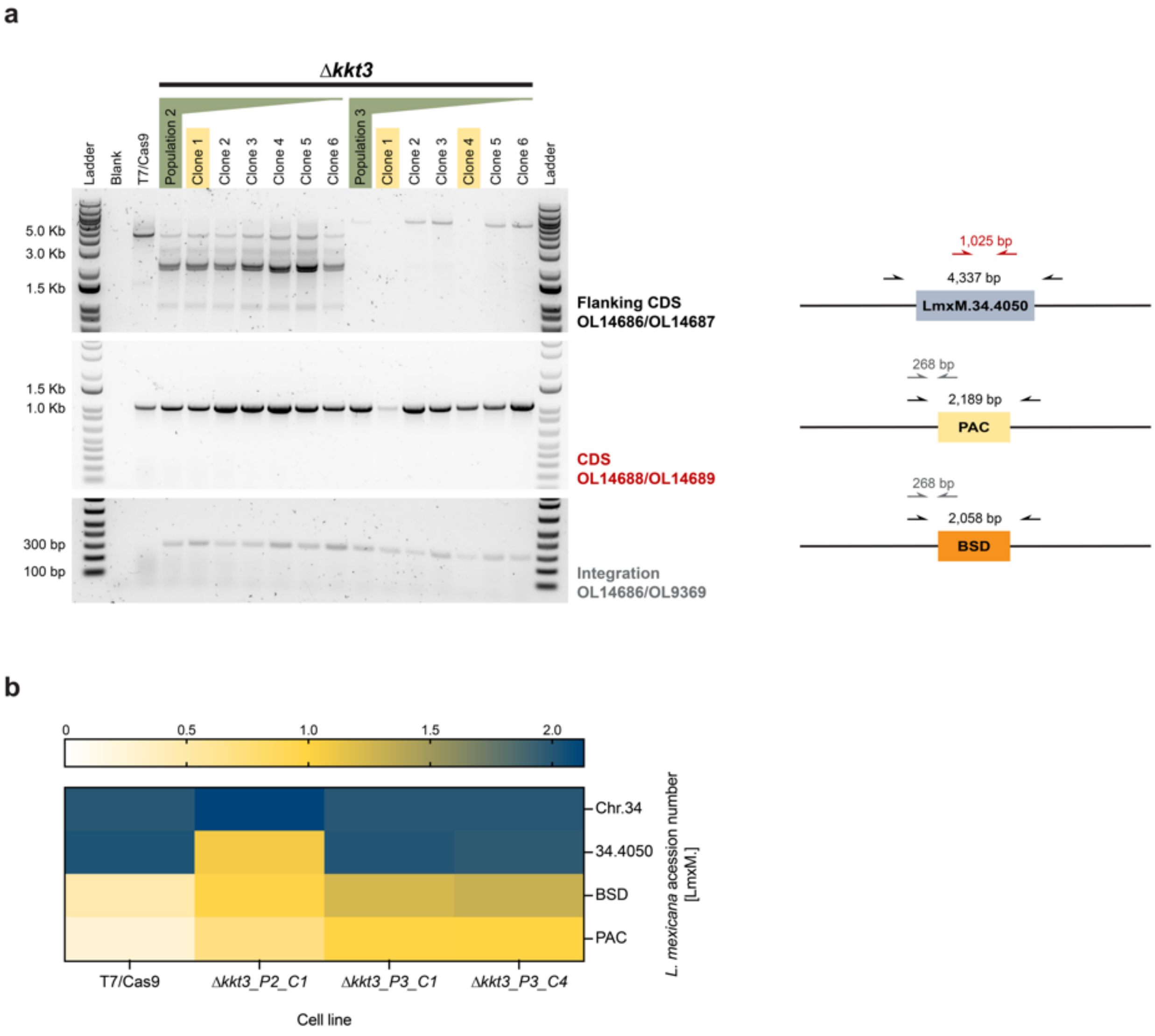

**Supplementary Fig 10. Requirement of KKT3 protein kinase for *L. mexicana* promastigote survival.** Three independent CRISPR-Cas9 transfections were performed to delete the KKT3 gene (LmxM.34.4050) using repair templates carrying blasticidin (*BSD*) and puromycin (*PAC*) resistance markers in combination with linear DNA fragments for *in vivo* transcription of two specific single guide RNAs. Following drug selection, viable populations emerged from transfections 2 and 3, and clones were subsequently isolated. (a) PCR amplification was conducted to assess the presence of the *KKT3* coding sequence (CDS) and integration of the drug resistance cassettes. The PCR strategy and expected amplicon sizes are illustrated in the schematic on the right. Clones highlighted in yellow were selected for whole-genome sequencing (WGS) using Illumina technology. (b) WGS analysis of Δ*kkt3* mutants. Illumina reads from clone 1 of population 2 (P2_C1), and clones 1 and 4 of population 3 (P3_C1 and P3_C4), were aligned to the reference genome of the T7/Cas9 parental cell line [5]. The heatmap shows the copy-number status of chromosome 34 and the *KKT3* locus. Chromosome ploidy was estimated by normalizing read depth to the average coverage of the four longest disomic chromosomes (Chr.08, Chr.20, Chr.33 and Chr.34), which were set to a baseline of 2. Gene copy number was estimated by the ratio of the gene coverage by its chromosome coverage, multiplied by the chromosome ploidy. WGS data for all analyzed mutants have been deposited in the SRA under project ID PRJNA1303394.

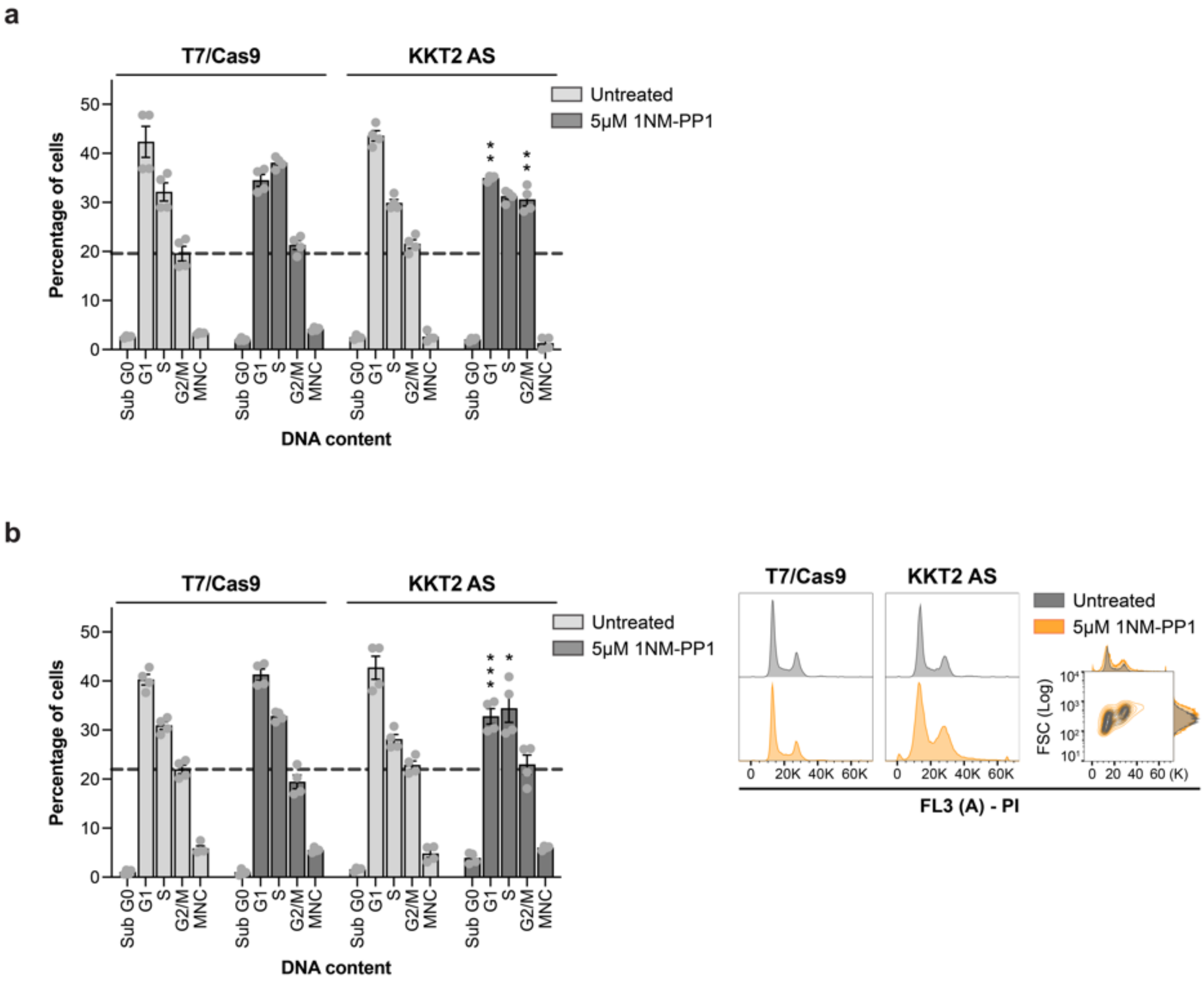

**Supplementary Fig 11. Effect of KKT2 kinase activity inhibition on *Leishmania* cell cycle progression.** The parental T7/Cas9 line and untreated parasites expressing KKT2 AS variant (KKT2$^{M146G}$) cultured under the same conditions were used as controls. Cell cycle analysis of cells stained with propidium iodide (PI) after 6 h (a) or 24 h (b) of treatment with 5 µM 1NM-PP1. Cell cycle phase quantification was performed using the Watson model algorithm in FlowJo v10.10.0. The left panel presents the percentage of cells in each cell cycle phase, while the right panel displays a representative cell cycle histogram. Adjacent histograms showing DNA content and forward scatter (FSC) were used to assess cell size across different cell cycle stages. Probability p-values were calculated using two-tailed Student's t-tests, comparing the percentage of cells in each phase between treated and untreated populations (* p < 0.05; **p < 0.01; *** p < 0.001). Data are mean ± SEM of four biological replicates.

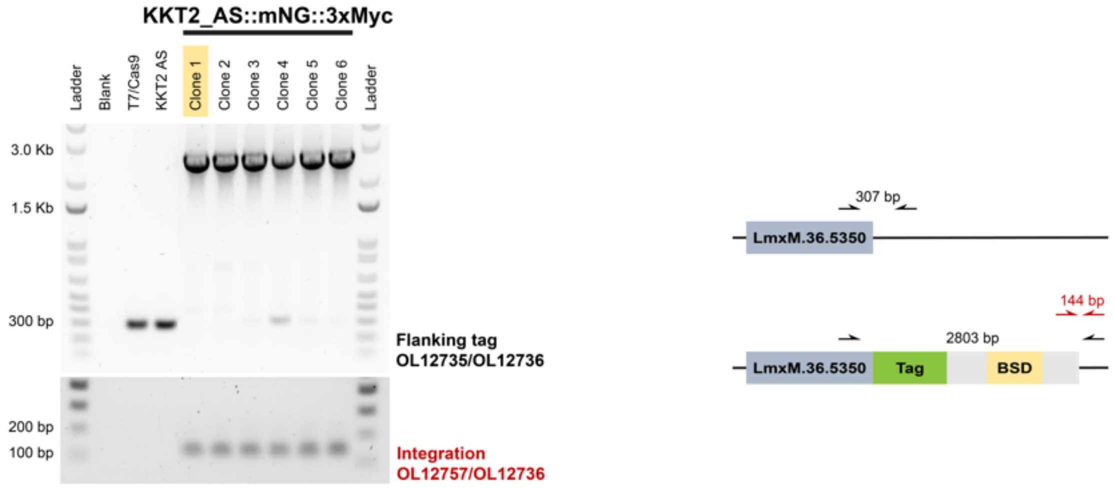

**Supplementary Fig 12. PCR screening of the KKT2_AS::mNG::3xMyc cell line.** The cell line expressing KKT2 AS variant (KKT2$^{M146G}$) was engineered using CRISPR-Cas9 to endogenously tag the C-terminus of both alleles of KKT2 AS with mNeonGreen (mNG) fused to the 3xMyc epitope, resulting in the KKT2_AS::mNG::3xMyc cell line. A repair template containing a blasticidin resistance marker and linear DNA fragments for *in vivo* transcription of the single guide RNA was transfected into the parental line. Following an overnight recovery period, transfected cells were selected with blasticidin and subsequently cloned by serial dilution. PCR amplification was performed to confirm the integration of the repair template at the correct locus in both alleles. The diagram on the right illustrates the PCR strategies and the expected DNA product sizes for each amplification. The clone highlighted in yellow was selected for subsequent experiments.

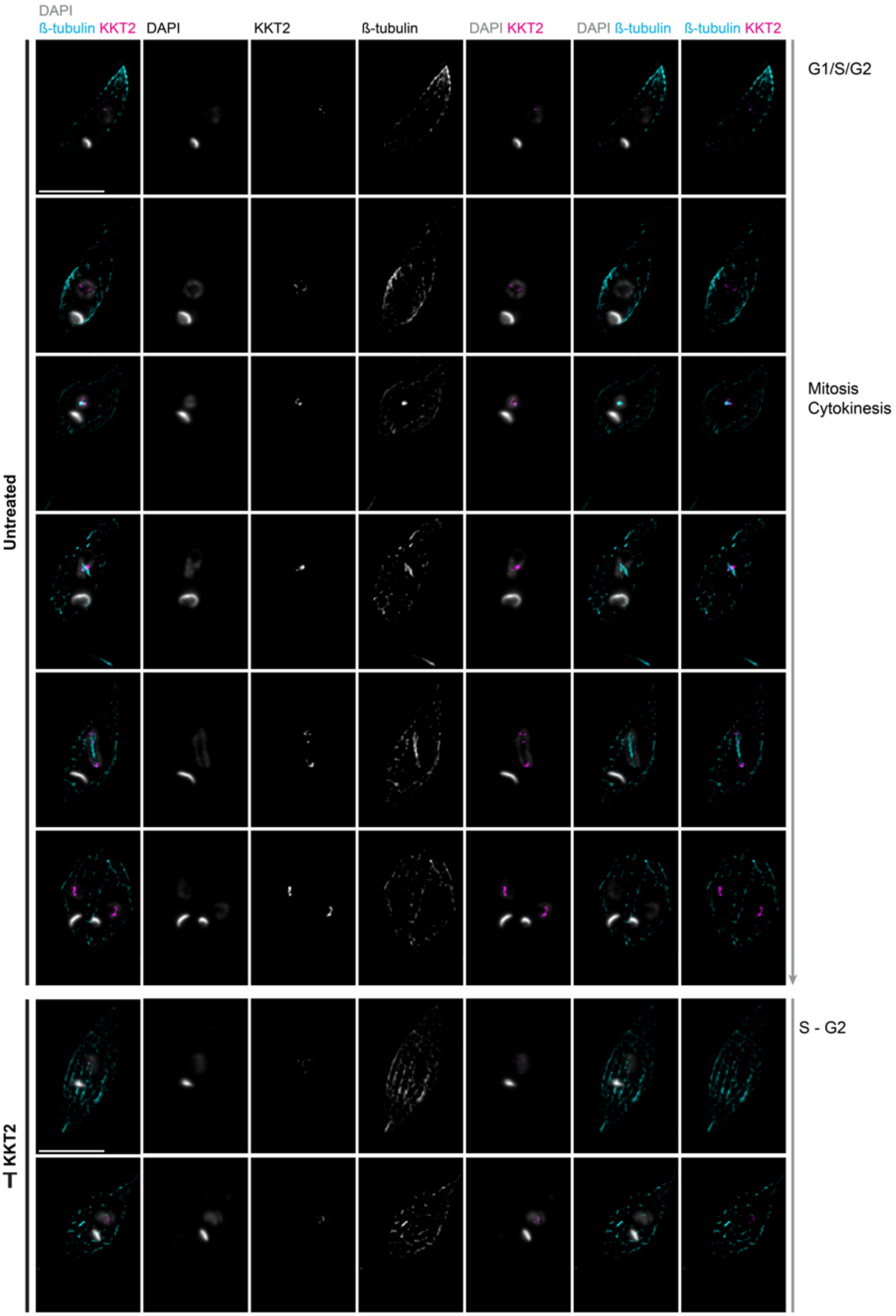

**Supplementary Fig 13. Spatial and temporal distribution of *L. mexicana* KKT2 throughout the cell cycle.** High-resolution fluorescence microscopy was performed on KKT2 AS::mNG::3xMyc promastigotes cultured with or without 10 µM 1NM-PP1 for 6 hours. Cells were stained with KMX-1 antibody to detect β-tubulin, anti-Myc antibody to visualize KKT2, and counterstained with DAPI for DNA visualization. Representative fluorescence micrographs show the localization of KKT2 and β-tubulin across different *Leishmania* cell cycle stages. The colours used for each marker are indicated to the top of the images. Scale bars, 5 μm.

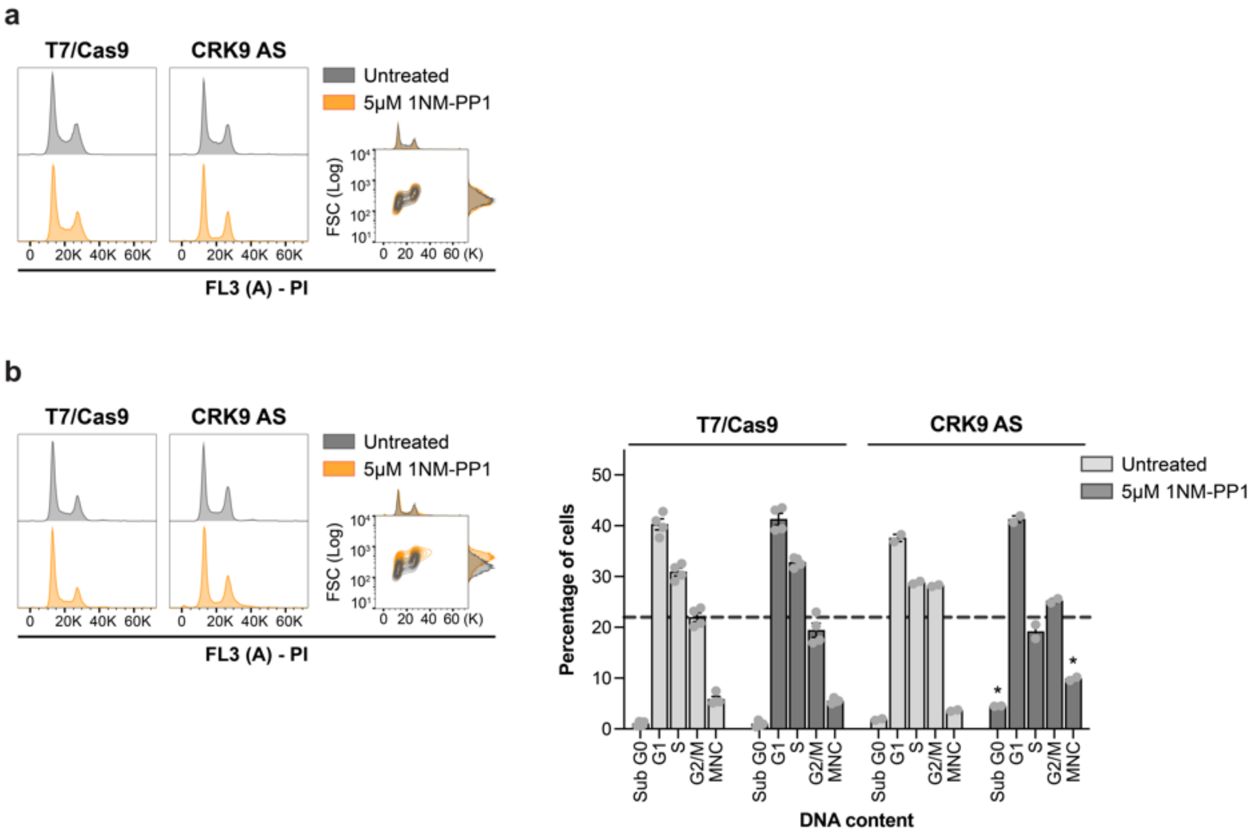

**Supplementary Fig 14. Effect of CRK9 kinase activity inhibition on *Leishmania* cell cycle progression.** The parental T7/Cas9 line and untreated parasites expressing CRK9 AS variant (CRK9$^{M501G}$) cultured under the same conditions were used as controls. Cell cycle analysis of cells stained with propidium iodide (PI) after 6 hours (a) or 24 hours (b) of treatment with 5 µM 1NM-PP1. Cell cycle phase quantification was performed using the Watson model algorithm in FlowJo v10.10.0. The left panel displays a representative cell cycle histogram, with adjacent histograms showing DNA content and forward scatter (FSC) were used to assess cell size across different cell cycle stages. The right panel presents the percentage of cells in each cell cycle phase. Probability p-values were calculated using two-tailed Student's t-tests, comparing the percentage of cells in each phase between treated and untreated populations (* $p < 0.05$; **$p < 0.01$; *** $p < 0.001$; **** $p < 0.0001$). Data are mean ± SEM of four biological replicates.

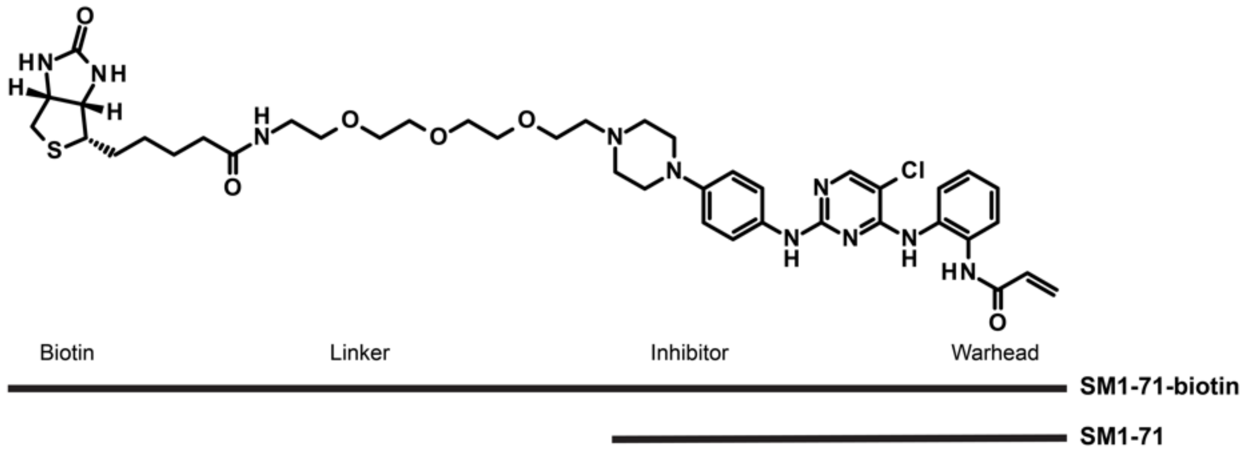

**Supplementary Fig 15. SM1-71 probes.** Chemical structures of SM1-71 and its biotinylated analog.

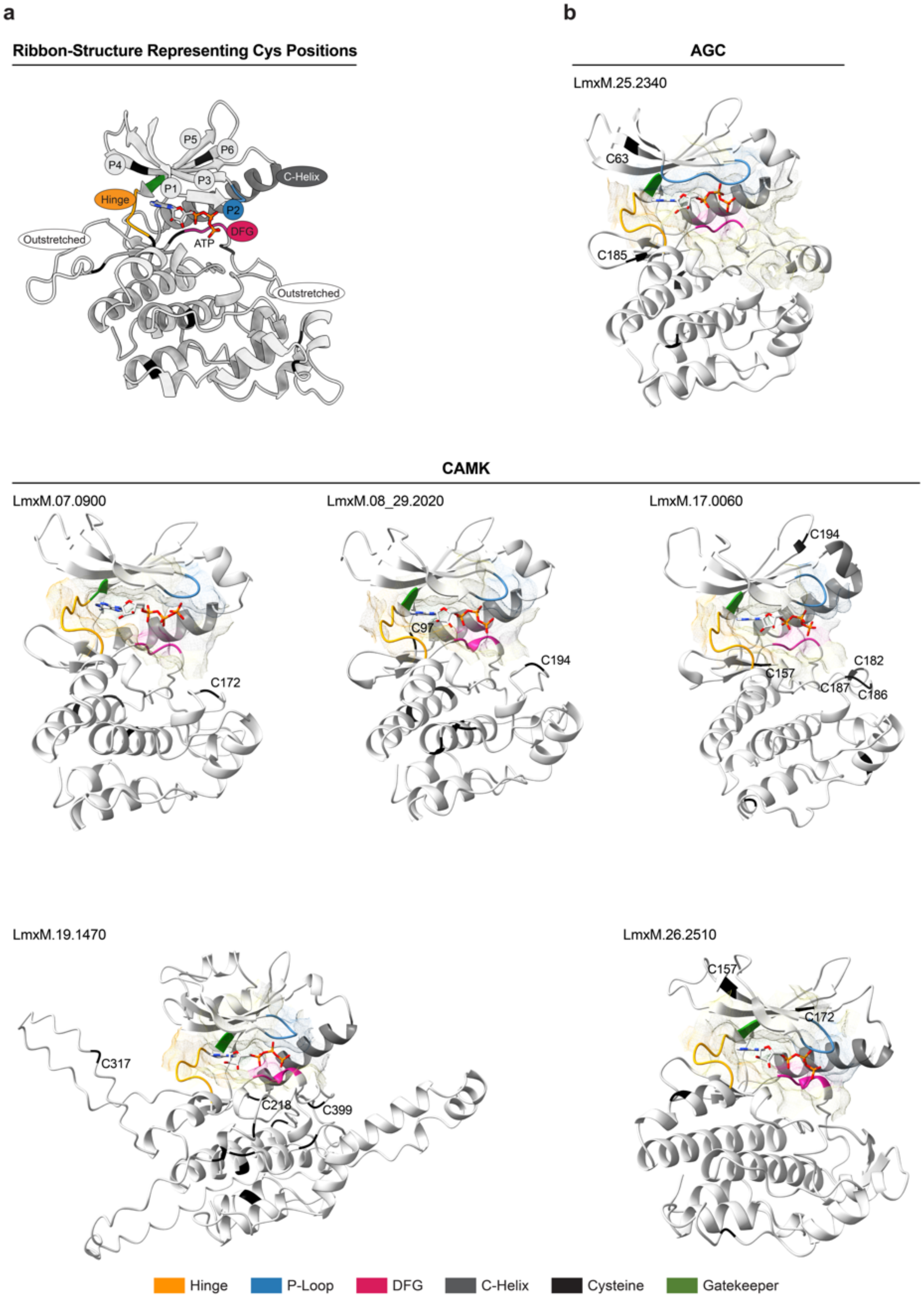

**Supplementary Fig 16. Predicted structural models of kinase domains from *L. mexicana* protein kinases enriched by the multi-targeted acrylamide-modified probe, SM1-71-biotin.**

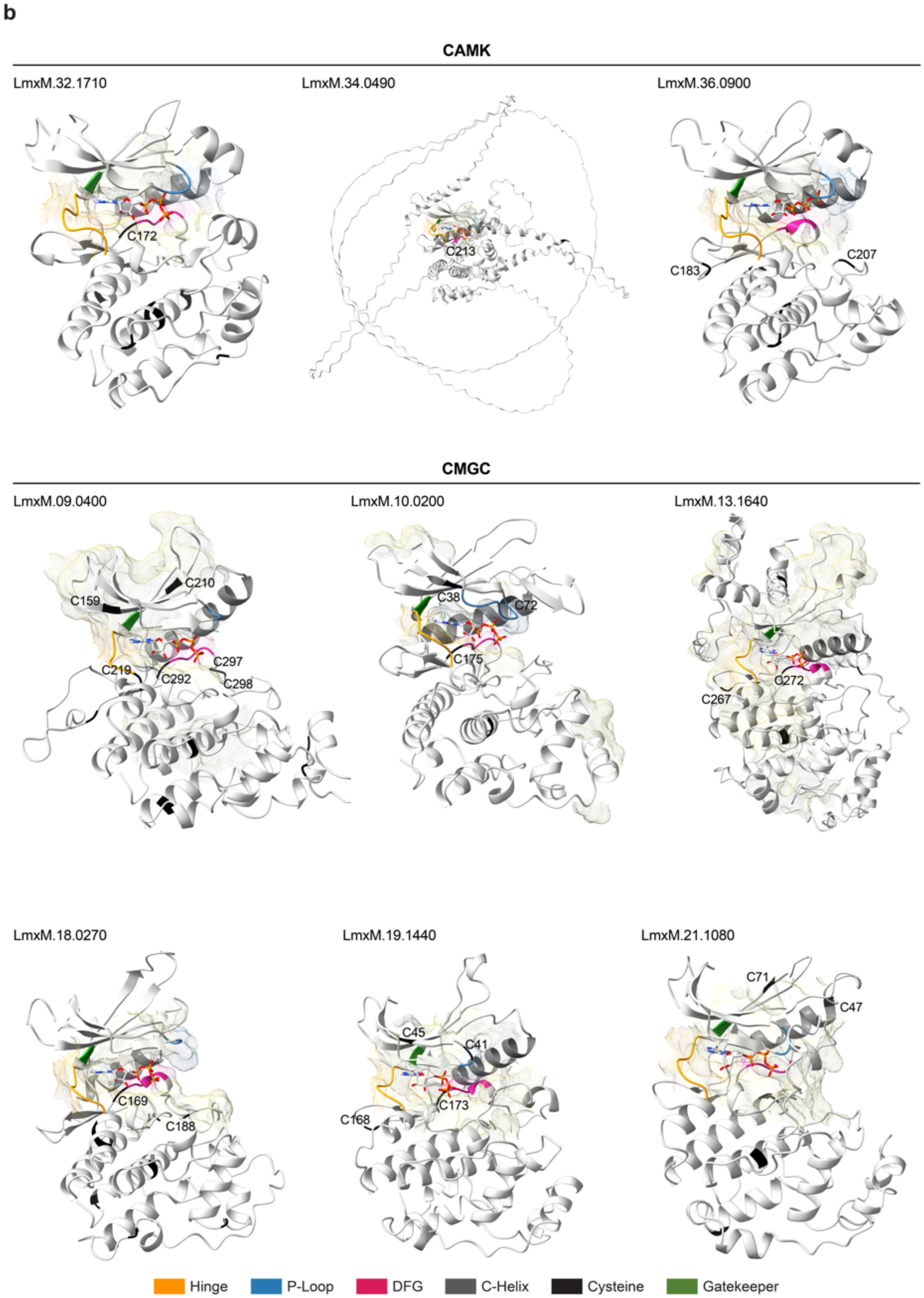

**Supplementary Fig 16. Predicted structural models of kinase domains from *L. mexicana* protein kinases enriched by the multi-targeted acrylamide-modified probe, SM1-71-biotin.**

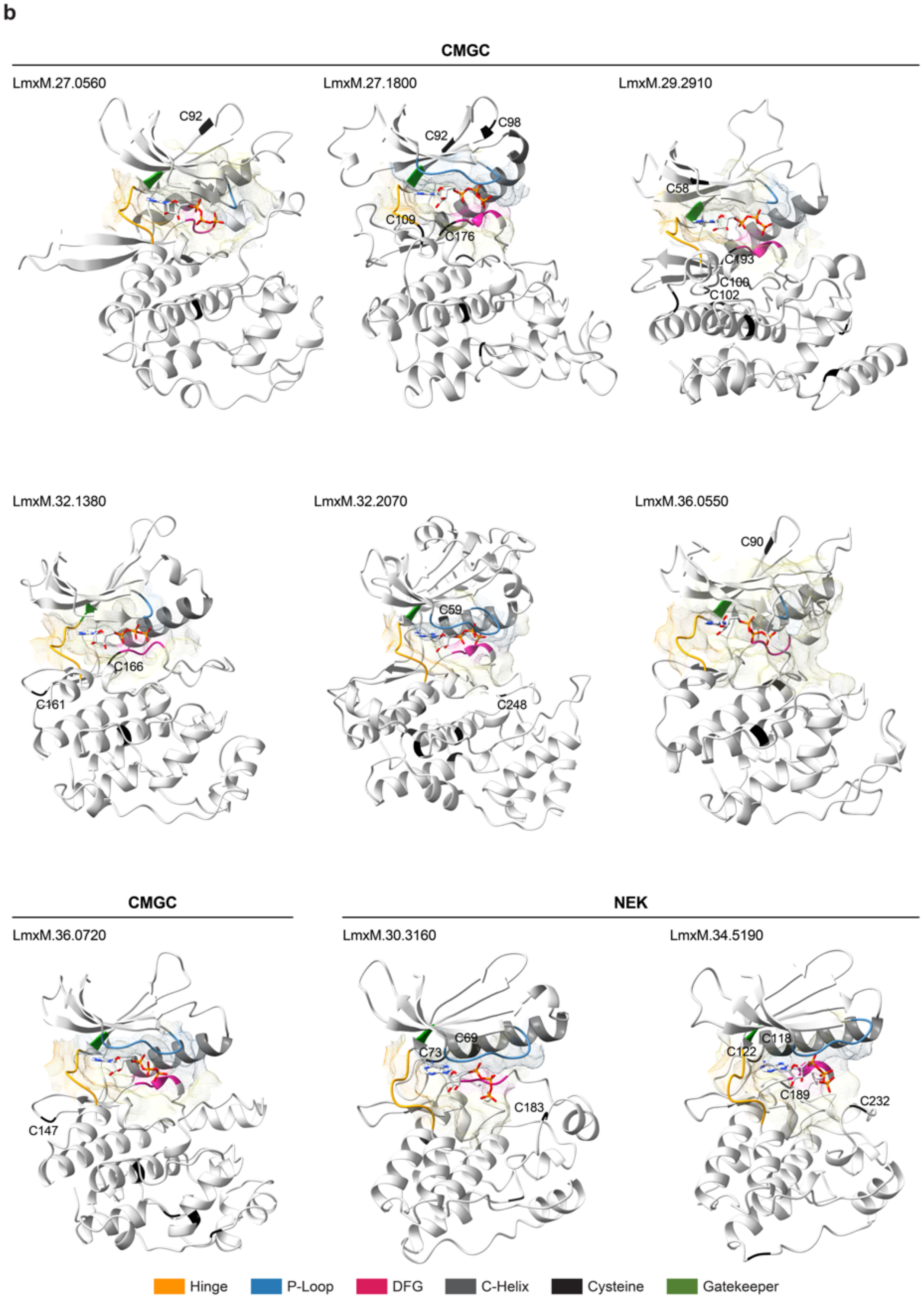

**Supplementary Fig 16. Predicted structural models of kinase domains from *L. mexicana* protein kinases enriched by the multi-targeted acrylamide-modified probe, SM1-71-biotin.**

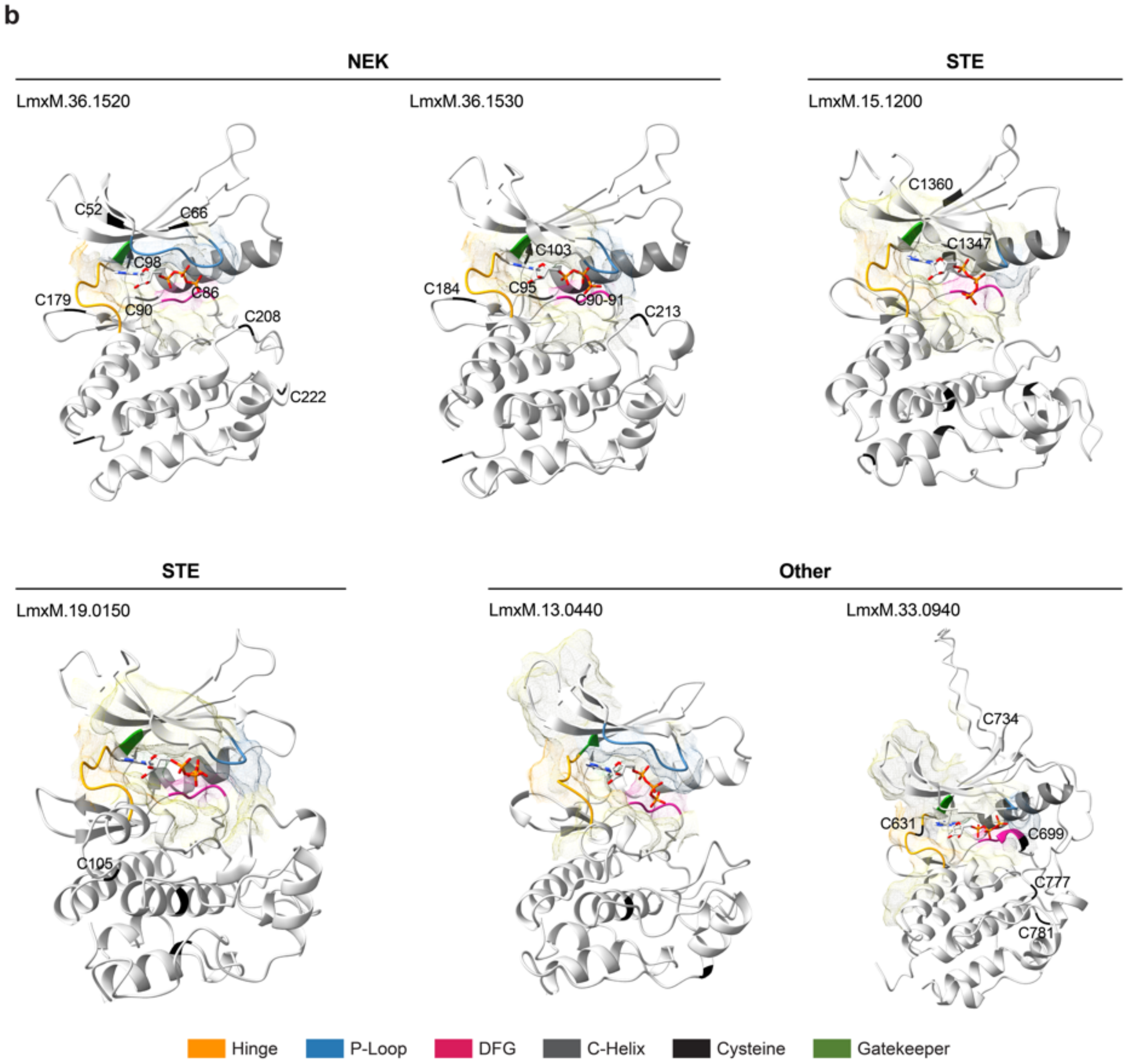

**Supplementary Fig 16. Predicted structural models of kinase domains from *L. mexicana* protein kinases enriched by the multi-targeted acrylamide-modified probe, SM1-71-biotin.** Structural models of the kinase domains were generated using AlphaFold 3 [6] and visualized with ChimeraX v1.9. The ATP ligand is displayed as a stick model, with heteroatom-based colouring. Residues located within 6 Å of the ATP-binding site are represented as a semitransparent surface overlay. AlphaFold confidence metrics, including the predicted template modelling score (pTM) and the interface predicted template modelling score (ipTM) for kinase-ATP interaction, are provided in Supplementary Data 2. (a) Ribbon diagram indicating the positions of cysteine residues within the kinase domain. (b) Predicted structural models of kinase domains from *L. mexicana* protein kinases identified as enriched by the multi-target acrylamide-based probe SM1-71-biotin.

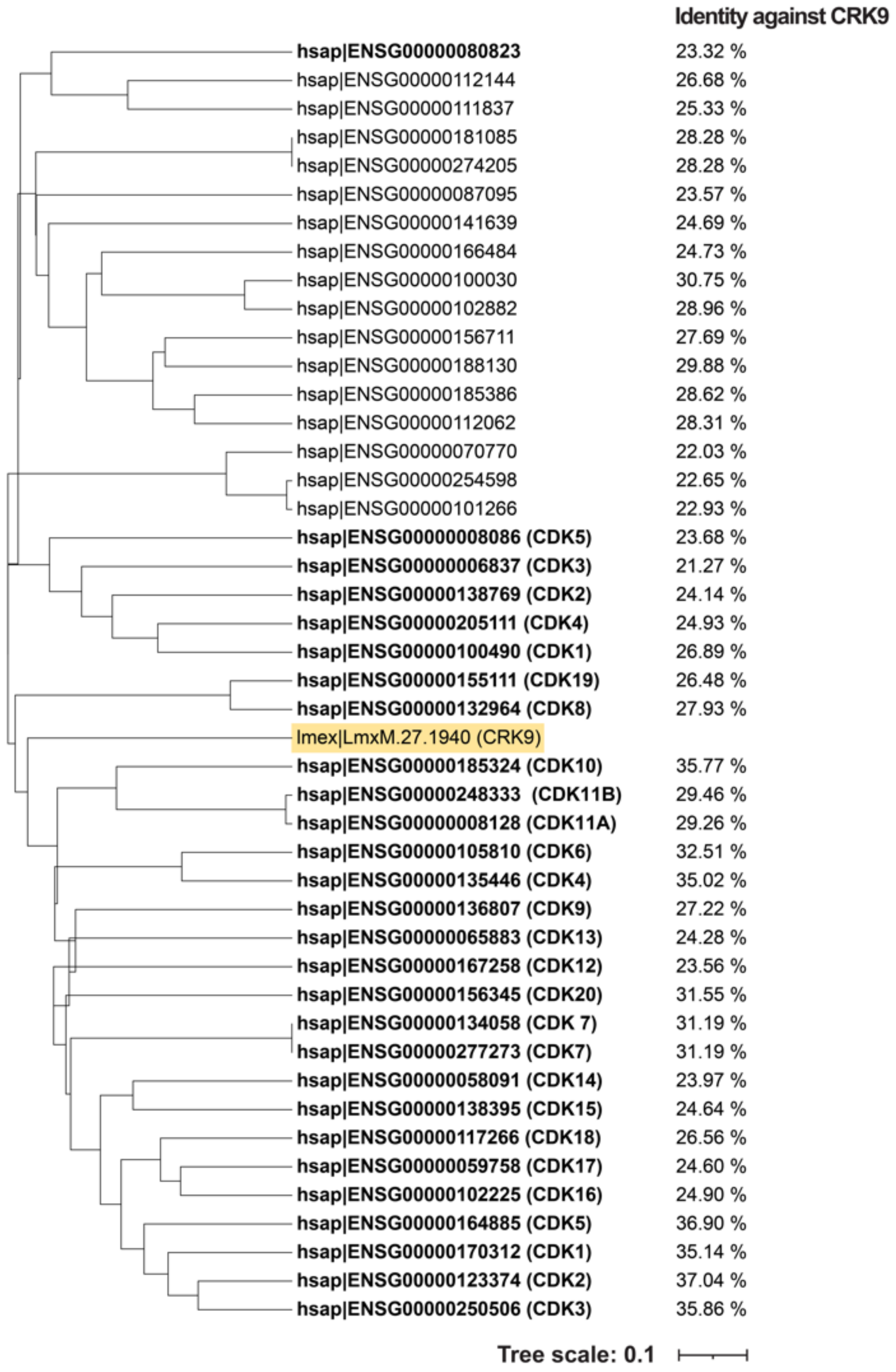

**Supplementary Fig 17. Phylogenetic analysis of human CDKs with sequence similarity to *L. mexicana* CRK9.** Phylogenetic tree visualization of *L. mexicana* CRK9 (LmxM.27.1940) and selected human CDKs, as identified using OrthoMCL BD (https://orthomcl.org/orthomcl/app). The *L. mexicana* CRK9 protein is highlighted in yellow, and human cyclin-dependent kinases (CDKs) are indicated in bold. Protein sequences were aligned using Clustal Omega (https://www.ebi.ac.uk/jdispatcher/msa/clustalo), and the percentage identity relative to *L. mexicana* CRK9 is shown on the right side of the tree.
